# Measurement-Dependent Parameter Recoverability in Neural Cardiovascular LPN Inference

**DOI:** 10.64898/2026.10.29.762665

**Authors:** Nanlong Sun, Yue Li, Sanjian Zhang, Yurui Chen, Lei Shi

## Abstract

Estimating cardiovascular lumped-parameter network (LPN) parameters from limited hemodynamic observations is a potentially non-unique inverse problem in cardiovascular model personalization. We investigated how parameter recoverability depends on measurement availability and noise, and whether poor recovery reflects limitations of neural inference or insufficient observational information. We developed Mada (measurement-availability and difficulty-aware), a probabilistic neural framework that provides parameter estimates and predictive uncertainty under incomplete observations. Using a closed-loop, four-chamber cardiovascular LPN with time-varying elastance, we generated a synthetic population of 650,000 cases comprising 20 chamber-mechanics and vascular parameters and up to 30 hemodynamic and timing observations. Under complete observations, Mada achieved macro *R*^2^ = 0.862, maintaining competitive accuracy across neural baselines while providing parameter-specific uncertainty. Recoverability varied substantially across parameters, with atrial active elastances remaining particularly difficult. For mask-aware Mada, macro *R*^2^ ranged from 0.066 with basic clinical measurements to 0.751 with a rich clinical set and 0.842 with all measurements, while uncertainty generally increased as informative measurements were removed. Bounded direct optimization achieved substantially smaller hemodynamic residuals but did not consistently recover the generating parameters. Complementary profile and local-sensitivity analyses revealed case-specific parameter compensation and weak observational directions, while target-specific fine-tuning offered little improvement for difficult atrial parameters. Increasing measurement noise degraded both parameter recovery and predictive-interval coverage, and similar trends were observed with a simpler probabilistic ResMLP. These findings demonstrate that accurate reconstruction of observed hemodynamics does not necessarily imply accurate recovery of the underlying LPN parameters, highlighting the importance of assessing parameter recoverability and predictive uncertainty when physiologically meaningful parameter estimates are sought.

## 1 Introduction

Lumped-parameter models, also referred to as zero-dimensional (0D) or lumped-parameter network (LPN) models, provide a computationally efficient representation of cardiovascular dynamics through systems of ordinary differential equations describing pressure, flow, vascular resistance, compliance, and chamber mechanics [1–4]. Despite their reduced dimensionality, these models can reproduce important characteristics of the systemic and pulmonary circulations and have been increasingly used for patient-specific hemodynamic analysis, multiscale modeling, and cardiovascular model personalization [5–8]. A central requirement for such personalization is the reliable inference of physiologically meaningful model parameters from measurements available for an individual subject.

Cardiovascular model personalization therefore requires solving an inverse problem: estimating latent quantities such as chamber elastances, unstressed volumes, vascular resistances, and compliances from observed pressures, volumes, flows, and related hemodynamic variables. Previous studies have addressed this problem using optimization, system identification, sensitivity analysis, sequential estimation, data assimilation, and Bayesian inference [7, 9– 13], with more recent work emphasizing efficient personalization and uncertainty-aware parameter inference [14, 15]. These studies demonstrate the feasibility of cardiovascular parameter estimation, but they also reveal a fundamental difficulty: the same observation set may constrain some physiological parameters strongly while providing substantially less information about others.

This difficulty is closely related to parameter identifiability. Structural identifiability describes whether a parameter can theoretically be determined uniquely from ideal observations, whereas practical identifiability additionally depends on measurement selection, noise, parameter sensitivity, and finite data [16–18]. Limited or weakly informative observations may therefore allow substantially different parameter combinations to produce similar modeled hemodynamics [10–13]. Poor recovery, however, may also arise from limitations of the inference procedure itself, because optimization and learned inverse approximations can introduce additional error in high-dimensional nonlinear inverse problems [19]. In this study, we use the term *inverse recoverability* to describe the empirical predictability of an individual LPN parameter from a specified hemodynamic observation space. Unlike formal structural identifiability, recoverability is evaluated for a finite parameter population and a particular measurement set and is therefore both parameter dependent and measurement dependent.

Measurement availability and quality are consequently central to practical parameter recovery. Clinical hemodynamic information may be obtained from arterial or intracardiac pressure measurements, catheterization, echocardiography, chamber-volume assessment, and derived circulatory indices, but the available measurements vary across subjects and clinical settings. Previous personalization studies have shown that measurement selection and information content can strongly affect parameter estimation [7, 10, 12, 13]. A useful personalization framework should therefore accommodate incomplete observation sets and remain interpretable as physiologically informative measurements are removed or corrupted by noise.

When sufficiently large ensembles of forward simulations are available, learning-based methods provide an efficient alternative to repeated case-wise optimization by learning an amortized mapping from hemodynamic observations to physiological parameters. Because multiple LPN parameters are inferred simultaneously, individual targets may exhibit substantially different learning difficulty, as commonly encountered in multi-output and multi-task regression [20–27]. Modern neural architectures provide flexible representations for such nonlinear mappings [28–31], but increased model capacity cannot recover physiological information that is insufficiently constrained by the observations. Recent cardiovascular studies illustrate complementary learning-based approaches. Salvador et al. [32] developed a latent neural ordinary differential equation surrogate of a whole-heart electromechanical model to accelerate repeated sensitivity, parameter-inference, and uncertainty-quantification calculations. Tong et al. [33] introduced InVAErt for amortized inference of hemodynamic LPNs and explicitly represented multiple parameter combinations compatible with observed data. These developments demonstrate the potential of neural methods for efficient cardiovascular inverse inference while also emphasizing the need to determine which individual parameters are actually supported by the available observations.

Uncertainty estimation is particularly important in this setting because deterministic inverse regression produces a point estimate even when the observations provide only weak constraints on a parameter. Probabilistic neural regression provides a means of estimating parameter-specific predictive uncertainty [34, 35], while approximate Bayesian inference, Monte Carlo dropout, and deep ensembles provide alternative uncertainty-estimation strategies [36–38]. Such uncertainty estimates are useful only when they meaningfully reflect prediction error and remain appropriately calibrated [39–42]. For cardiovascular personalization, parameter-specific uncertainty may therefore provide complementary information about whether an estimated parameter is well supported by the available physiological information. Its reliability must also be reassessed when measurement availability or noise differs from the conditions under which calibration was performed.

To investigate these issues, we developed Mada (measurement-availability and difficulty-aware), a probabilistic neural inverse framework that produces parameter estimates together with parameter-specific predictive uncertainty while accommodating incomplete observations. Mada incorporates an empirical validation-derived recoverability descriptor and a mask-aware formulation so that a single model can operate across different measurement-availability patterns. We evaluated the framework using a synthetic population of 650,000 cardiovascular simulations spanning broad physiologically plausible variation in chamber mechanics, unstressed volumes, and systemic and pulmonary vascular properties. Each case contains known generating values for 20 inverse LPN parameters together with up to 30 hemodynamic observations. This large controlled population enables direct assessment of parameter recovery, systematic variation of measurement availability and noise, and comparison between learned inverse inference and forward-model-based optimization.

The study was designed to address three related questions. First, how heterogeneous is empirical recoverability across individual cardiovascular LPN parameters? Second, how do measurement availability, physiological information content, and measurement noise affect parameter recovery and predictive uncertainty? Third, when a parameter remains difficult to recover, to what extent does the limitation arise from the learned inverse approximation rather than insufficient constraint from the available observations?

We address these questions through complementary learning-based and physics-based analyses. Neural inverse performance and parameter-specific predictive uncertainty are evaluated across complete, incomplete, clinically motivated, and noise-perturbed observation conditions. We further compare neural inference with bounded direct optimization using the original dataset-generation LPN forward solver and fixed configuration. We also use target-specific and profile-based analyses to investigate persistent recovery limitations. Finally, additional neural baselines distinguish findings specific to Mada from more general properties of the observation-to-parameter inverse problem.

## 2 Methods

### 2.1 Cardiovascular LPN Data and Inverse Problem

The synthetic population used in this study was generated using the closed-loop, four-chamber cardiovascular lumped-parameter network (LPN) shown in Fig. 1. The model comprises atrial and ventricular time-varying elastance elements, pressure-dependent valve resistances, and systemic and pulmonary resistance–compliance–inertance compartments, following standard formulations [1–3] and our previous implementation [43]. The model structure is summarized in Appendix A.1.

**Figure 1.**
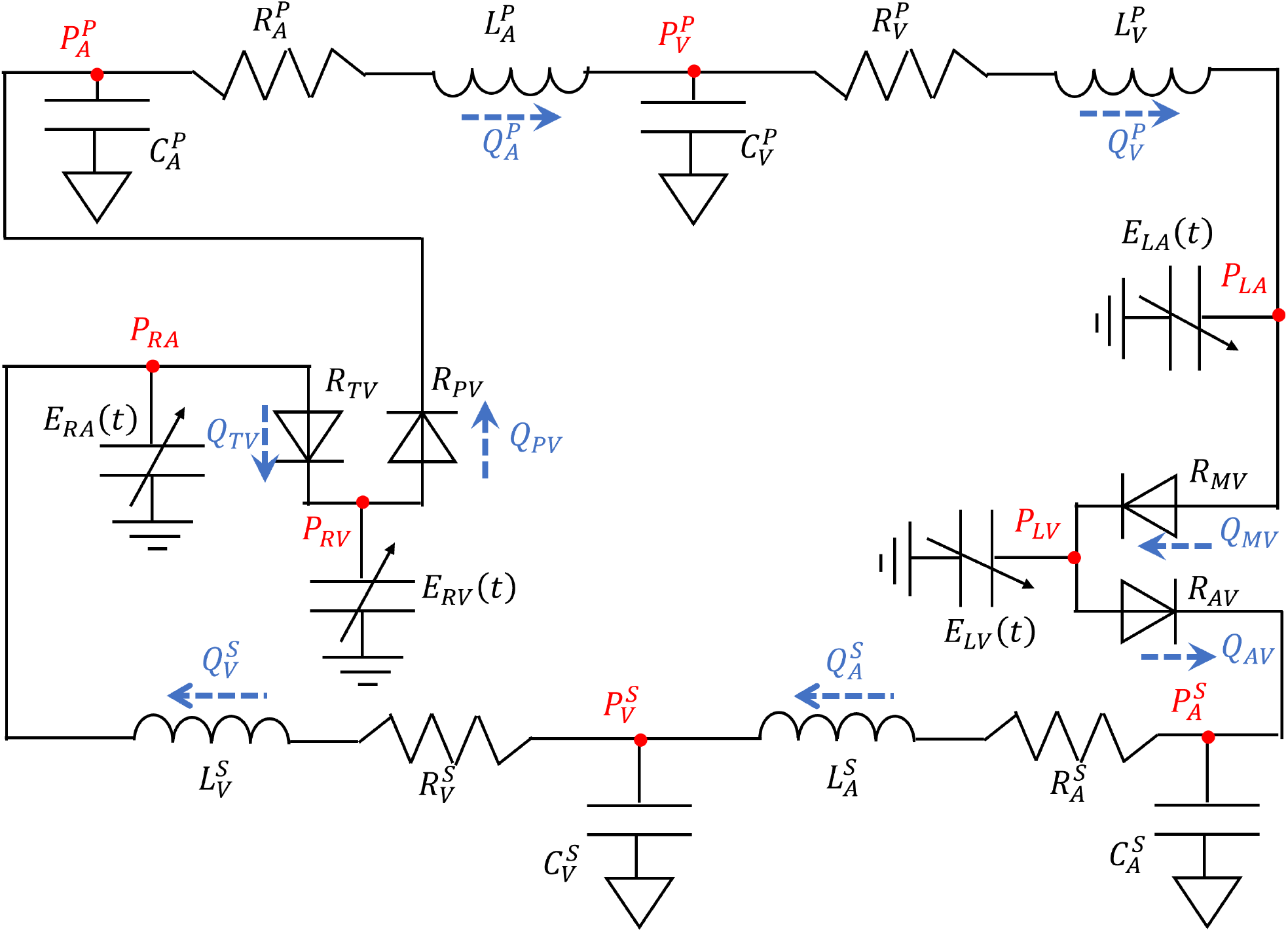
Closed-loop cardiovascular LPN used to generate the synthetic population. The left and right atria (LA, RA) and ventricles (LV, RV) are represented by time-varying elastance elements, while the systemic (S) and pulmonary (P) circulations contain arterial and venous resistance, compliance, and inertance elements. Valve resistances regulate directional flow between adjacent compartments. Only the selected chamber-mechanics and vascular parameters described below were treated as inverse targets.

For each simulated case, the physiological parameter vector is

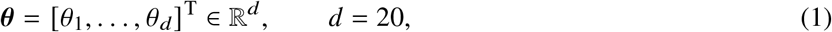

where *θ*_*j*_ denotes the *j*th LPN parameter. The parameter population was selected to span broad physiologically plausible variation in the modeled cardiovascular system. The realized parameter ranges in the final synthetic population are reported in Table 6 in Appendix A.2.

The case-specific timing vector is

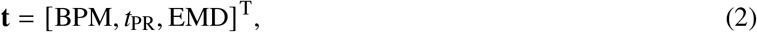

where BPM is heart rate in beats per minute, *t*_PR_ is the PR interval, and EMD is electromechanical delay.

For each sampled pair (***θ*, t**), the forward LPN solver produces 27 modeled hemodynamic quantities, **F**(***θ***; **t**) *∈* ℝ^27^.

Together with the three timing quantities, these form the complete *p* = 30 dimensional observation vector

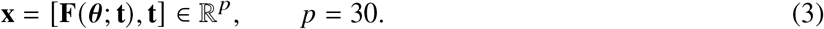

The 20 inverse targets comprise active and passive chamber elastances, unstressed chamber volumes, and systemic and pulmonary vascular resistances and compliances. The 30 observations comprise pressures, chamber volumes, stroke volumes, ejection fractions, global hemodynamic indices, ventricular stroke work, and cardiac timing variables (Table 1).

**Table 1.**
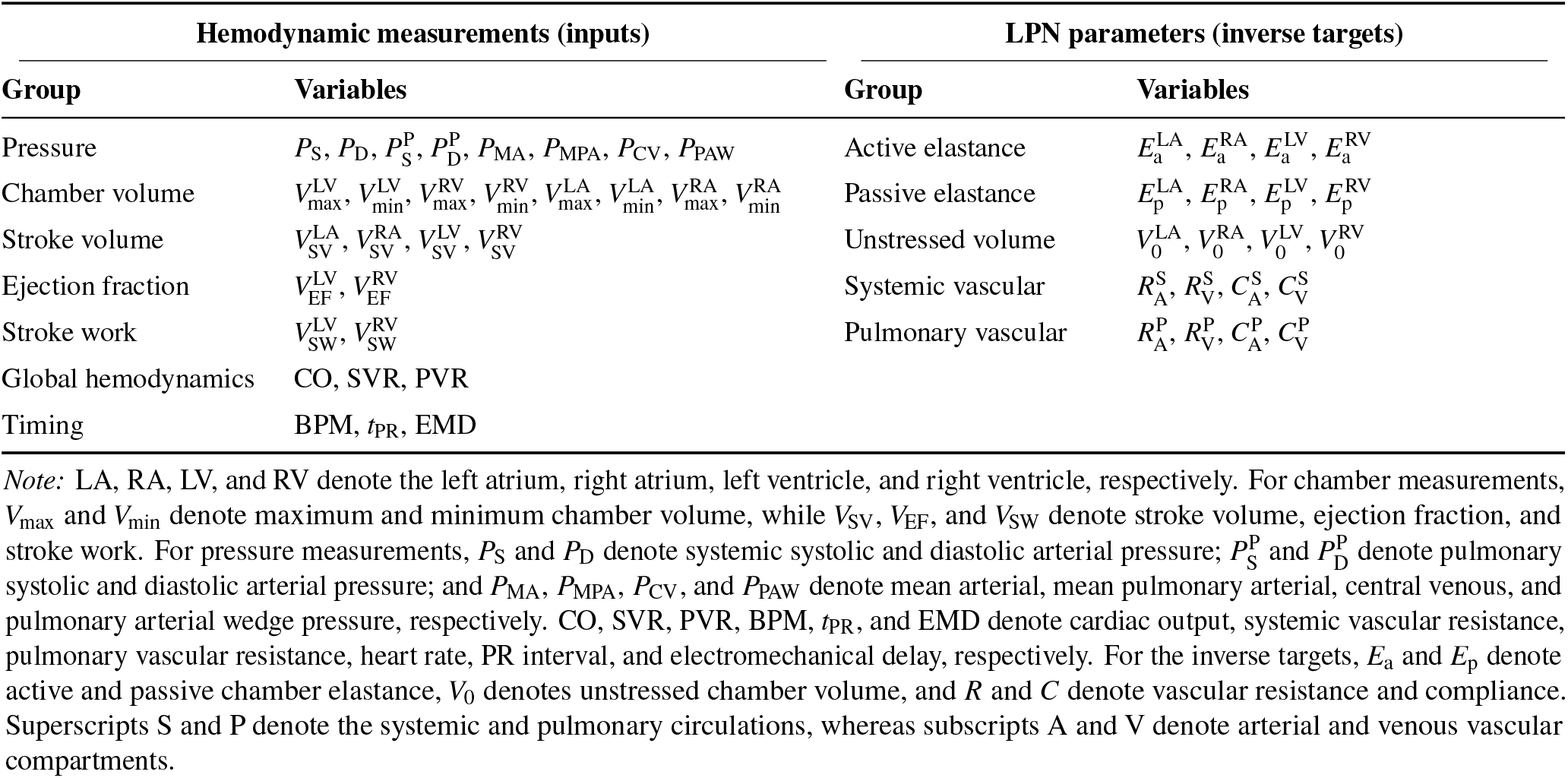
Physiological grouping of the 30 hemodynamic measurements used as model inputs and the 20 LPN parameters used as inverse-prediction targets.

The inverse-learning dataset is

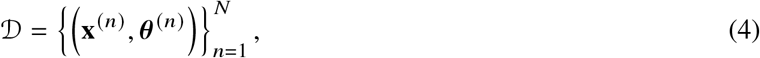

where *n* indexes simulated cases and *N* = 650,000 is the total number of samples. A fixed 80%/10%/10% partition was used for training, validation, and testing, yielding 520,000, 65,000, and 65,000 samples, respectively. The same partition was retained for all model development and evaluation.

The complete-observation inverse problem is to estimate

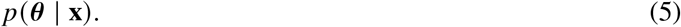

To represent incomplete observations, a binary availability mask **m** *∈* {0, 1}^*p*^ is introduced, where *m*_*i*_ = 1 indicates that measurement *x*_*i*_ is available and *m*_*i*_ = 0 indicates that it is missing. With **x**_obs_ = **x** *⊙* **m**, where *⊙* denotes element-wise multiplication, the general inverse problem becomes

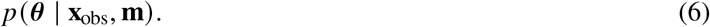

All normalization statistics were estimated exclusively from the training subset. For observation *i* and inverse target *j*,

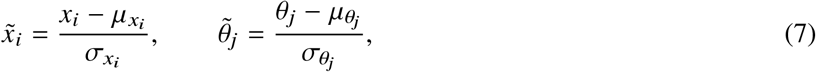

where *i* = 1, …, *p, j* = 1, …, *d*, and *µ* and *σ* denote the corresponding training-set mean and standard deviation. Tildes denote normalized quantities. Neural-network optimization was performed in this normalized space. All reported recoverability results should be interpreted conditional on the synthetic population considered here.

### 2.2 Mada Architecture

Mada uses a two-stage framework that separates empirical characterization of parameter recoverability from probabilistic inverse prediction. Stage 1 trains a deterministic reference predictor and uses its validation performance to estimate parameter-wise recoverability scores and characteristic residual scales. Stage 2 then trains a separate probabilistic predictor while holding the Stage 1 model fixed. The Stage 1 prediction enters Stage 2 only as an empirical reference through a recoverability-weighted regularization term. The overall framework is illustrated in Fig. 2.

**Figure 2.**
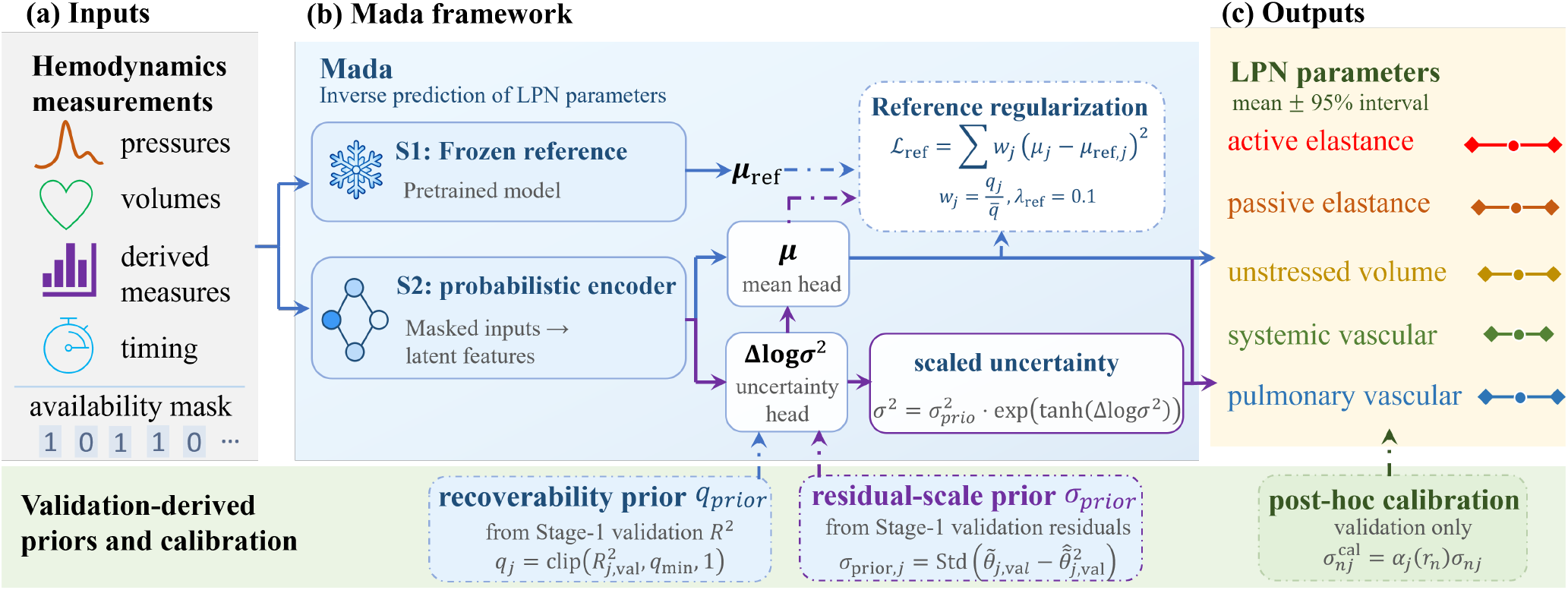
Overview of the proposed Mada framework for uncertainty-aware inverse estimation of cardiovascular LPN parameters. A deterministic Stage 1 reference predictor is first trained, and its validation predictions are used to derive parameter-wise recoverability scores and residual-scale priors. During Stage 2, the reference predictor remains fixed while a separate probabilistic predictor estimates the final parameter mean and sample-specific predictive uncertainty. Recoverability enters the Stage 2 objective through a weighted penalty on deviation of the final prediction from the fixed Stage 1 reference, while validation-derived residual scales provide parameter-specific centers for the predictive variance model. For incomplete observations, normalized measurements are accompanied by an explicit availability mask. Validation-only calibration is subsequently applied to predicted uncertainty. The recoverability score is an empirical measure of inverse predictability rather than a formal structural-identifiability index.

For the mask-aware model, the normalized measurements and their availability mask are combined as

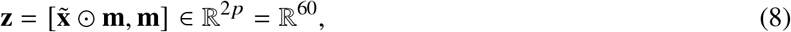

where **z** is the network input. The full-observation model instead uses 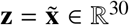, without concatenating an all-ones mask. The corresponding Stage 1 and Stage 2 models use the same input representation within each variant.

#### 2.2.1 Reference Prediction and Validation-Derived Priors

In Stage 1, a deterministic dense multilayer perceptron *f*_ref_ is trained on the training subset using an *L*_2_ prediction loss. For input **z**, it produces the normalized parameter estimate

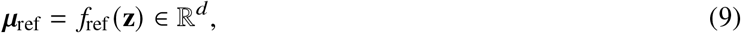

where ***µ***_ref_ is the 20-dimensional reference prediction. After Stage 1 training, *f*_ref_ is frozen and is not updated during Stage 2 optimization.

The trained reference predictor is then evaluated on the validation subset. For target *j*, its validation coefficient of determination 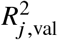 is used as an empirical measure of inverse recoverability:

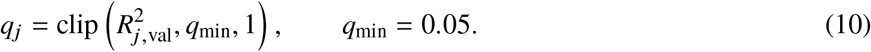

The recoverability-prior vector

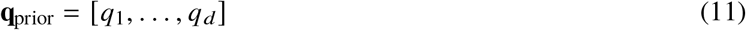

therefore summarizes the relative empirical recoverability of the *d* = 20 targets. Larger *q*_*j*_ indicates stronger predictive support from the available observation space. The lower bound *q*_min_ prevents low or negative validation *R*^2^ values from producing a vanishing score.

The same validation predictions define a characteristic residual scale for each target,

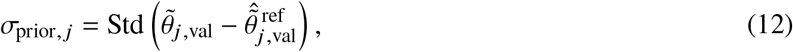

where 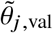 is the normalized validation target and 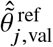 is the corresponding Stage 1 prediction. Thus, *q*_*j*_ characterizes point-prediction recoverability, whereas *σ*_prior, *j*_ characterizes the typical Stage 1 residual scale. Both are derived exclusively from validation data.

#### 2.2.2 Probabilistic Prediction and Predictive Uncertainty

Stage 2 uses a separate probabilistic predictor. The input **z** is projected to a 256-dimensional latent space and passed through fully connected residual blocks,

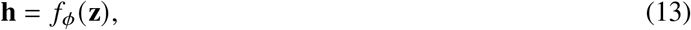

where *f*_*ϕ*_ denotes the shared nonlinear encoder and **h** its latent representation. A mean head *f*_*µ*_ maps **h** directly to the final normalized parameter estimate,

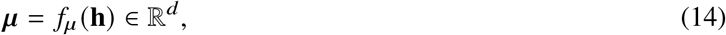

where *µ*_*j*_ is the predictive mean for target *j* . The Stage 1 reference prediction ***µ***_ref_ enters the Stage 2 objective through the regularization term defined in Sec. 2.2.3.

In parallel, an uncertainty head *f*_*σ*_ predicts a sample-specific adjustment vector,

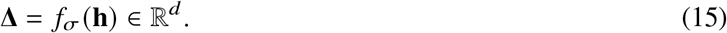

Mada represents the normalized inverse targets using conditionally independent heteroscedastic Gaussian distributions,

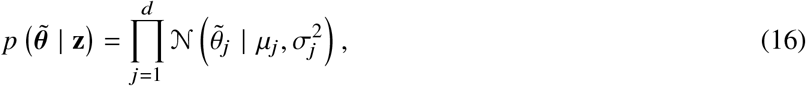

where 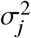 is the predicted variance for target *j* in normalized parameter space. Its log variance is centered on the validation-derived Stage 1 residual scale,

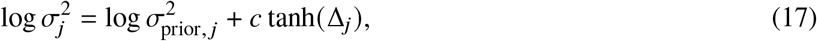

where *c* bounds the sample-specific departure from the prior scale in log-variance space. The full-observation model used *c* = 1.0. For the mask-aware model, *c* = 2.5 was used to permit broader uncertainty adaptation under information loss. The resulting log variance was additionally clipped to [−7, 4] for numerical stability.

#### 2.2.3 Recoverability-Weighted Reference Regularization

For *N* training samples and *d* = 20 targets, the Stage 2 probabilistic predictor is optimized using the heteroscedastic Gaussian negative log-likelihood

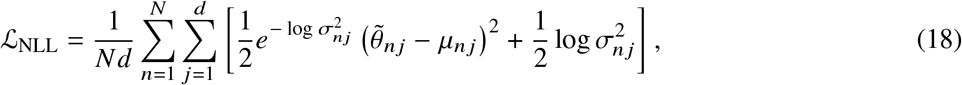

where 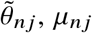, and 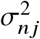 denote the normalized target, predicted mean, and predicted variance, respectively, for sample *n* and target *j* .

The Stage 1 recoverability scores are normalized to unit mean:

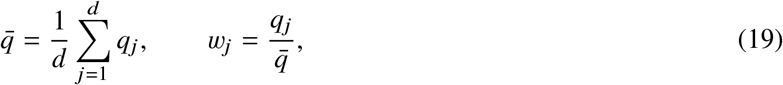

where 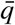 is the mean recoverability score and *w*_*j*_ is the corresponding target-specific reference weight.

The Stage 2 mean is then penalized for deviating from the fixed Stage 1 reference,

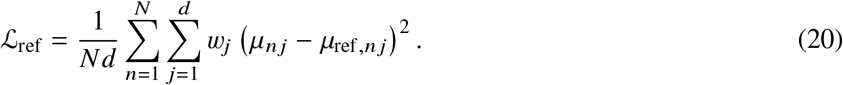

Targets that are more reliably predicted by Stage 1 therefore receive a stronger penalty for departure from the reference, whereas targets with lower empirical recoverability are less strongly constrained. The penalty acts directly on the final Stage 2 predictive mean.

The complete Stage 2 objective is

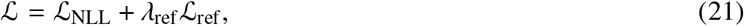

where *λ*_ref_ ≥ 0 controls the strength of the reference regularization.

### 2.3 Training, Calibration, and Evaluation

Separate full-observation and mask-aware variants were trained using the same data partition and training-derived normalization statistics. The full-observation models received complete inputs throughout training. For the mask-aware models, 25% of samples retained complete observations, whereas for the remaining 75% a missing fraction

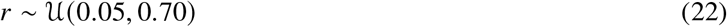

was sampled and individual measurements were randomly masked subject to at least one observation remaining available. The same mask-generation distribution was used for the mask-aware Stage 1 reference and Stage 2 probabilistic predictors.

After Stage 1 training, the reference predictor was frozen and excluded from the Stage 2 optimizer. All neural models were implemented in PyTorch and optimized using AdamW. For mask-aware Mada, the initial learning rate was 3 × 10−4, batch size was 256, and weight decay was 10−4. Training was allowed for up to 300 epochs with an early-stopping patience of 30 epochs. The encoder used a hidden dimension of 256, two residual blocks, and dropout probability 0.1. The uncertainty-adjustment output layer was initialized to zero so that the initial predictive variance equaled the validation-derived residual scale in Eq. (12).

The full-observation and mask-aware Stage 2 predictors contained 420,668 and 428,348 trainable parameters, respectively. The difference of 7,680 parameters corresponds to the additional 30 × 256 input-projection weights associated with the availability mask.

The reference-regularization coefficient was selected using a matched validation sweep,

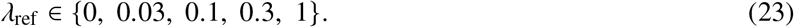

All candidates used the same Stage 1 reference, Stage 2 initialization, training split, stochastic sequence, optimizer settings, and checkpoint criterion. Checkpoints were selected using the lowest unregularized validation NLL. The sweep selected *λ*_ref_ = 0.1, which was subsequently fixed for both the full-observation and mask-aware Mada models. The complete matched sweep is reported in Appendix D.4.

Predictive uncertainty was calibrated using validation data only. Let ℓ index the evaluation condition, corresponding to a prescribed random-missingness level, a removed physiological measurement group, or a fixed observation scenario. For condition ℓ and inverse target *j*, an initial validation scale ratio was defined as

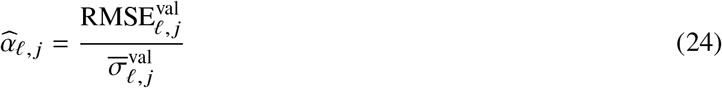

where 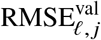 is the validation root-mean-square prediction error and

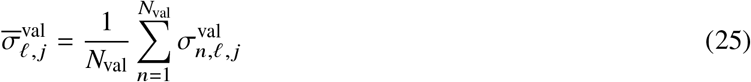

is the corresponding mean uncalibrated predictive standard deviation over the *N*_val_ validation cases. Both quantities were computed in the same parameter space. For the dedicated full-observation model, the applied factor was 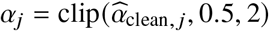.

For mask-aware random and structured measurement removal, a least-squares linear fit of 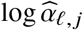 against the five prescribed validation missingness fractions *r*_ℓ_ *∈* { 0, 0. 1, 0. 3, 0 5, 0. 7 } gave an intercept *a*_*j*_ and slope *b*_*j*_ . The applied factor was *α*_*n*,ℓ, *j*_ = exp(*a*_*j*_ + *b*_*j*_*r*_*n*,ℓ_ ), where *r*_*n*,ℓ_ is the actual fraction of unavailable measurements in sample *n*. For structured removal this fraction is fixed by the removed group. For each fixed observation scenario, the validation ratio was instead clipped to [0.4, 2.5] and applied separately to each target. The dedicated full-observation and fixed-scenario factors are constant across samples.

The calibrated test-set standard deviation was then

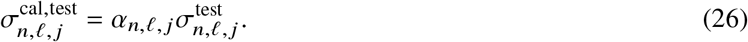

No test-set targets were used for model selection or uncertainty calibration.

The two-stage mask-aware training procedure is summarized in Algorithm 1.

#### Algorithm 1

Two-stage training procedure for mask-aware Mada

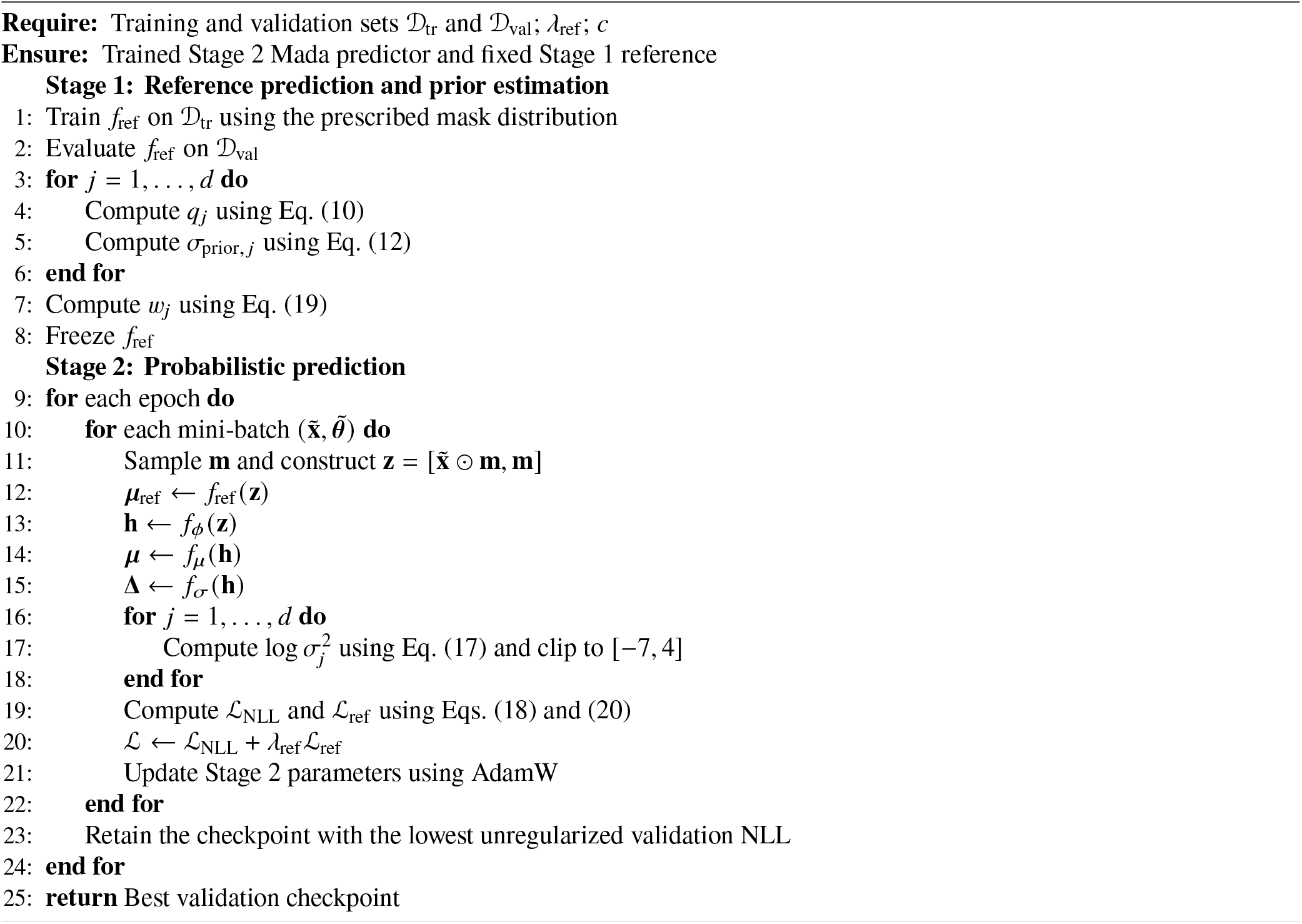

Model performance was evaluated using parameter-wise and macro-averaged coefficient of determination (*R*^2^), normalized root-mean-square error (NRMSE), and Pearson correlation coefficient (PCC). Predictive uncertainty was evaluated using the mean predictive standard deviation, the ratio of mean predictive standard deviation to empirical root-mean-square error (Std./RMSE), and 95% prediction-interval coverage probability (PICP_95_). Complete metric definitions are provided in Appendix E.

### 2.4 Measurement-Availability Experiments

Measurement availability was evaluated using three complementary protocols.

First, random measurements were removed at test-time fractions of 0%, 10%, 30%, 50%, and 70%. These same missingness levels were used for validation-based uncertainty calibration.

Second, seven physiological measurement groups—pressure, chamber volume, stroke volume, ejection fraction, global hemodynamics, stroke work, and timing—were removed one at a time. For removed group *g* and inverse target *j*, the recoverability loss was defined as

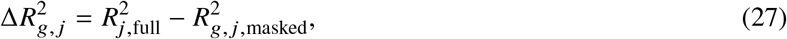

where 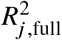 is the test-set *R*^2^ with all measurements available and 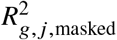 is the corresponding value after group *g* is removed. The predictive-uncertainty inflation factor was

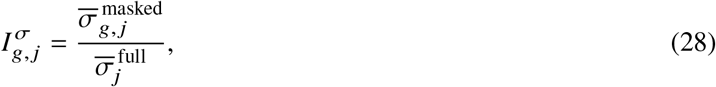

where the numerator and denominator are the corresponding mean predictive standard deviations. These quantities are empirical measurement–parameter dependency measures rather than formal structural-identifiability indices.

For the correlation analysis, the absolute change in normalized calibrated predictive uncertainty was additionally defined as

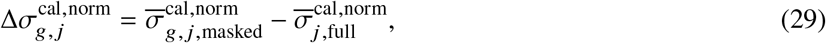

where the two terms are the mean normalized calibrated predictive standard deviations for target *j* after removal of group *g* and under complete observations, respectively.

Third, the frozen mask-aware Mada checkpoint was evaluated under six fixed observation scenarios: basic clinical (systemic systolic and diastolic pressure plus heart rate; 3/30), BP + chamber volumes (10/30), Echo-like (15/30), hemodynamic (eight pressures plus cardiac output; 9/30), rich clinical (23/30), and full synthetic (30/30). No scenario-specific retraining or fine-tuning was performed. Because these fixed masks differed from the random masks used during training, scenario-specific multiplicative uncertainty calibration was estimated from the validation set using Eq. (B1). Detailed observation-set definitions are provided in Appendix B.3.4.

### 2.5 Diagnostic Analyses of Poor Recoverability

The neural experiments identify which parameters are difficult to recover but do not by themselves distinguish limitations of the observation space from limitations of the learned inverse mapping. Four complementary diagnostics were therefore performed.

First, a physics-based inverse benchmark estimated the 20 LPN parameters through bounded optimization using the same forward solver and model configuration employed to generate the synthetic dataset.

The three timing variables **t** = [BPM, *t*_PR_, EMD]^T^ were retained as known case-specific inputs, while the remaining 27 hemodynamic quantities contributed to the standardized forward residual. For modeled output *k*,

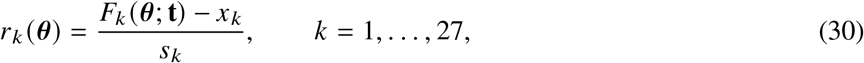

where *F*_*k*_ (***θ***; **t**) is the *k*th modeled hemodynamic output, *x*_*k*_ is the corresponding synthetic observation, and *s*_*k*_ is its training-set standard deviation.

The bounded inverse estimate was

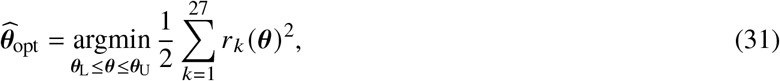

where ***θ***_L_ and ***θ***_U_ are the original lower and upper generating bounds used for inverse estimation.

Parameter recovery was summarized using the range-normalized absolute error

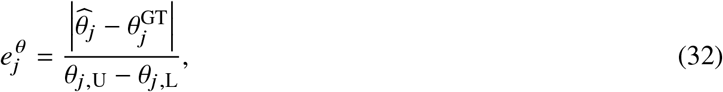

where 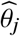 is the recovered value, 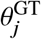 is the generating value, and *θ*_*j*,L_ and *θ*_*j*,U_ are the original lower and upper generating bounds used in the direct-optimization benchmark.

Observation-space consistency was quantified using the normalized root-mean-square forward error

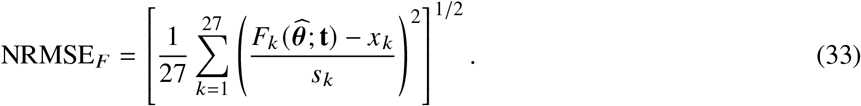

A five-case diagnostic and a broader 20-case space-filling cohort were evaluated. Optimizer settings and initialization rules are provided in Appendix C.1.

Second, target-specific single-output residual multilayer perceptron (ResMLP) fine-tuning was performed for 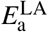 and 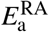 to test whether their poor recovery was primarily caused by interference among the 20 simultaneously predicted targets.

Third, profile-style compensation analyses were performed for a retained representative case (Case 07). The atrial active elastances were fixed at prescribed values while a targeted set of physiologically related parameters was re-optimized. One- and two-dimensional profiles were used to determine whether substantially different atrial active-elastance values could be compensated by other parameters while preserving similar hemodynamic outputs.

Fourth, for the five representative optimization cases, the normalized forward Jacobian at the generating parameter vector was analyzed by singular-value decomposition. The resulting singular-value spectrum, spectral conditioning, and effective rank were used to characterize weak local parameter directions.

The single-output, profile, and local-sensitivity protocols are described in Appendices C.4, C.5, and D.6, respectively.

### 2.6 Robustness and Architecture Diagnostics

Two additional analyses tested the robustness and architectural specificity of the primary findings.

First, a modality-dependent measurement-noise experiment perturbed pressure, chamber-volume, timing, flow, and stroke-work quantities in physical units before applying the original training normalization. The frozen full-observation Mada model was evaluated without noisy-data retraining or recalibration. The same noise realization was also used in a five-case original-LPN optimization diagnostic. A common-amplitude sweep scaled all prescribed perturbations by

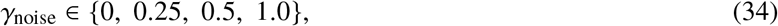

while retaining the same stochastic realization, allowing degradation of parameter accuracy and uncertainty calibration to be examined without changing the underlying noise pattern. Perturbation levels, original-LPN noise procedures, and amplitude-sweep details are provided in Appendices D.1 and D.5.

Second, mask-aware Mada was compared with a simpler probabilistic ResMLP using the same mask-aware ResMLP backbone but excluding the two Mada-specific components: Stage 1 recoverability-weighted reference regularization and the *σ*_prior_-centered uncertainty formulation. The simpler model used direct mean and log-variance heads and was trained with the same heteroscedastic Gaussian NLL. Architecture and training details for this architecture comparison are provided in Appendix D.3.

## 3 Results

### 3.1 Parameter-Wise Recoverability Under Complete Observations

Table 2 summarizes full-observation performance for a standard multilayer perceptron (MLP), a residual multilayer perceptron (ResMLP), an FT-Transformer, a heteroscedastic Gaussian negative-log-likelihood (Gaussian-NLL) model, and the dedicated full-observation Mada model (details of the other models are summarized in Sec. A.3). The deterministic ResMLP achieved the highest macro *R*^2^ (0.871), followed by Mada (0.862). Mada achieved a macro NRMSE of 0.276 and macro PCC of 0.921, compared with 0.272 and 0.927 for ResMLP. The MLP, FT-Transformer, and Gaussian-NLL baselines achieved macro *R*^2^ values of 0.835, 0.815, and 0.819, respectively. Thus, the dedicated full-observation Mada model retained point-prediction accuracy close to the strongest deterministic baseline while additionally providing parameter-specific predictive uncertainty.

**Table 2.** Full-observation inverse-prediction performance and model complexity on the held-out test set. Boldface identifies the best point-prediction metrics (*R*^2^, NRMSE, and PCC). For Mada, the parameter count refers to the trainable Stage 2 predictor; the frozen Stage 1 reference used during training is excluded. Mada uncertainty is reported after validation-only calibration, whereas Gaussian-NLL uncertainty is uncalibrated; therefore Std./RMSE and PICP_95_ are reported descriptively and are not used for direct model ranking.

| Model | Key architectural features | Params | Macro $R^2$ | Macro NRMSE | Macro PCC | Std./RMSE | PICP<sub>95</sub> |
| --- | --- | --- | --- | --- | --- | --- | --- |
| MLP | Standard MLP | 0.146M | 0.835 | 0.326 | 0.905 | — | — |
| ResMLP | Residual backbone | 0.544M | **0.871** | **0.272** | **0.927** | — | — |
| FT-Transformer | Feature-attention architecture | 0.276M | 0.815 | 0.358 | 0.893 | — | — |
| Gaussian-NLL | Heteroscedastic predictive uncertainty | 0.151M | 0.819 | 0.341 | 0.897 | 1.207 | 0.986 |
| **Full-observation Mada (ours)** | Frozen reference; reference regularization; predictive uncertainty; recoverability prior | 0.421M | 0.862 | 0.276 | 0.921 | **1.045** | **0.974** |

Aggregate metrics concealed substantial parameter-wise variation (Fig. 3). The atrial active elastances were the most difficult targets, with *R*^2^ = 0.351 for 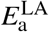 and 0.361 for 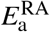. In contrast, 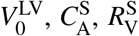 and 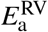 achieved *R*^2^ = 1.000, 1.000, 0.997, and 0.997, respectively.

**Figure 3.**
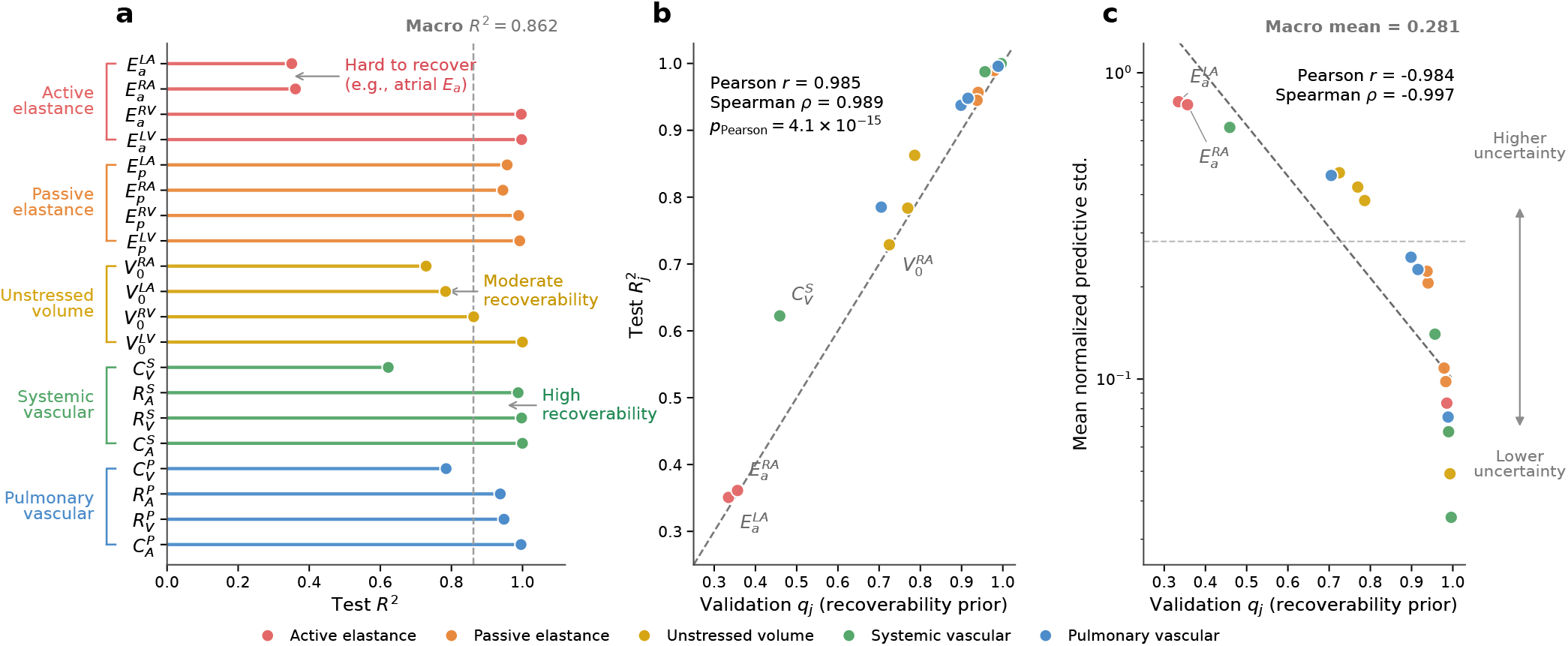
Parameter-wise inverse recoverability and recoverability-prior behavior under complete observations. (a) Test-set 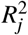 across the 20 LPN parameters. (b) Relationship between the validation-derived parameter-wise recoverability score *q*_*j*_ and held-out test 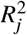, where *r* and *ρ* denote the Pearson and Spearman correlation coefficients, respectively, and *p*_Pearson_ denotes the two-sided Pearson-correlation *p*-value. (c) Relationship between recoverability and parameter-specific predictive uncertainty. The recoverability score is an empirical predictability descriptor rather than a formal structural-identifiability measure. Because *q*_*j*_ and *σ*_prior, *j*_ are both derived from Stage 1 validation residuals, and Stage 2 predictive variance is centered on *σ*_prior, *j*_, their cross-parameter association is partly induced by construction and is interpreted descriptively rather than as independent uncertainty validation.

The validation-derived recoverability score was strongly associated with held-out test-set recoverability: across the 20 parameters, the Pearson correlation between *q*_*j*_ and test-set 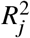 was *r* = 0.985, and the Spearman rank correlation was *ρ* = 0.989. The matched regularization sweep showed that weak recoverability-weighted reference regularization provided a modest improvement in aggregate predictive accuracy.

After validation-only calibration, Mada achieved a mean Std./RMSE ratio of 1.045 and mean PICP_95_ of 0.974. The most difficult and easiest targets were broadly consistent across architectures (Table 8). Coverage exceeded the nominal 0.95 level on average; calibration did not make every parameter’s uncertainty ratio equal to one.

The mask-aware Mada model used in the measurement-availability experiments was trained across variable missingness patterns and is therefore evaluated separately below.

### 3.2 Measurement Availability Determines Recoverability and Uncertainty

The mask-aware model showed a progressive loss of inverse accuracy as random measurements were removed. Macro *R*^2^ decreased from 0.842 with complete observations to 0.800, 0.690, 0.535, and 0.337 at 10%, 30%, 50%, and 70% missingness, respectively. Over the same range, the validation-calibrated mean normalized predictive standard deviation increased from 0.314 to 0.866 (Fig. 4). Thus, point-prediction accuracy decreased while reported predictive uncertainty increased as information was removed.

**Figure 4.**
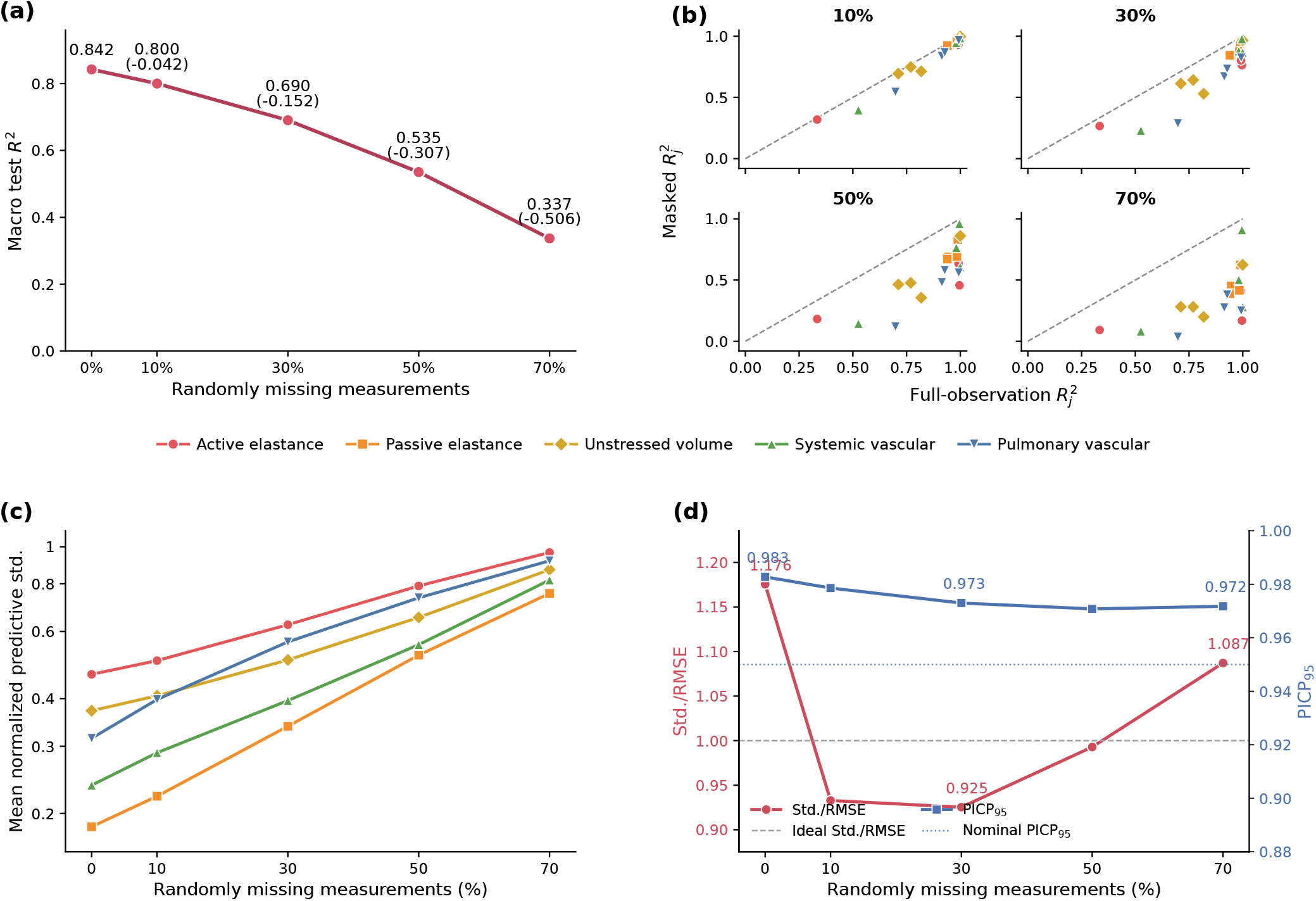
Robustness of mask-aware Mada to random measurement missingness. (a) Macro test *R*^2^; parenthesized values indicate the change in macro *R*^2^ relative to complete observations. (b) Parameter-wise recoverability relative to complete observations. (c) Mean validation-calibrated normalized predictive standard deviation across LPN parameter groups. (d) Missingness-conditioned uncertainty calibration, reported using Std./RMSE and PICP_95_.

At 70% missingness, mean PICP_95_ was 0.972 and the mean Std./RMSE ratio was 1.087. PICP_95_ remained approximately 0.97–0.98 across the evaluated missingness levels. These results describe aggregate calibration; they do not imply nominal coverage for every target.

Structured measurement removal showed that physiological content mattered more than measurement count alone (Table 3 and Fig. 5). Pressure removal caused the largest macro-averaged recoverability loss (0.275), followed by chamber-volume removal (0.111) and stroke-work removal (0.076). Pressure measurements were especially important for systemic and pulmonary vascular parameters; chamber volumes strongly constrained unstressed chamber volumes; and stroke work strongly constrained active ventricular mechanics. Validation-calibrated uncertainty inflation broadly followed the same dependency structure.

**Table 3.**
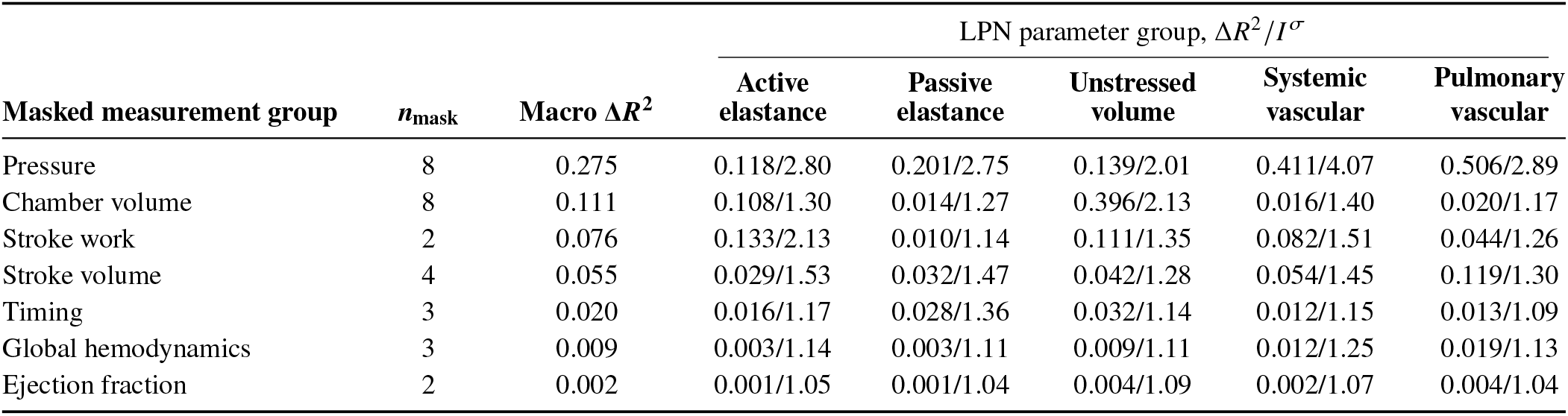
Effect of masking physiological measurement groups on parameter recoverability and predictive uncertainty. Here, *n*_mask_ denotes the number of measurements removed in each physiological group. For each LPN parameter group, entries are reported as mean recoverability loss Δ*R*^2^ / mean calibrated uncertainty inflation *I* ^*σ*^. The macro Δ*R*^2^ column summarizes the overall loss across all inferred parameters.

| Masked measurement group | $n_{\text{mask}}$ | Macro $\Delta R^2$ | LPN parameter group, $\Delta R^2 / I^\sigma$ | | | | |
| --- | --- | --- | --- | --- | --- | --- | --- |
|  |  |  | Active elastance | Passive elastance | Unstressed volume | Systemic vascular | Pulmonary vascular |
| Pressure | 8 | 0.275 | 0.118/2.80 | 0.201/2.75 | 0.139/2.01 | 0.411/4.07 | 0.506/2.89 |
| Chamber volume | 8 | 0.111 | 0.108/1.30 | 0.014/1.27 | 0.396/2.13 | 0.016/1.40 | 0.020/1.17 |
| Stroke work | 2 | 0.076 | 0.133/2.13 | 0.010/1.14 | 0.111/1.35 | 0.082/1.51 | 0.044/1.26 |
| Stroke volume | 4 | 0.055 | 0.029/1.53 | 0.032/1.47 | 0.042/1.28 | 0.054/1.45 | 0.119/1.30 |
| Timing | 3 | 0.020 | 0.016/1.17 | 0.028/1.36 | 0.032/1.14 | 0.012/1.15 | 0.013/1.09 |
| Global hemodynamics | 3 | 0.009 | 0.003/1.14 | 0.003/1.11 | 0.009/1.11 | 0.012/1.25 | 0.019/1.13 |
| Ejection fraction | 2 | 0.002 | 0.001/1.05 | 0.001/1.04 | 0.004/1.09 | 0.002/1.07 | 0.004/1.04 |

**Figure 5.**
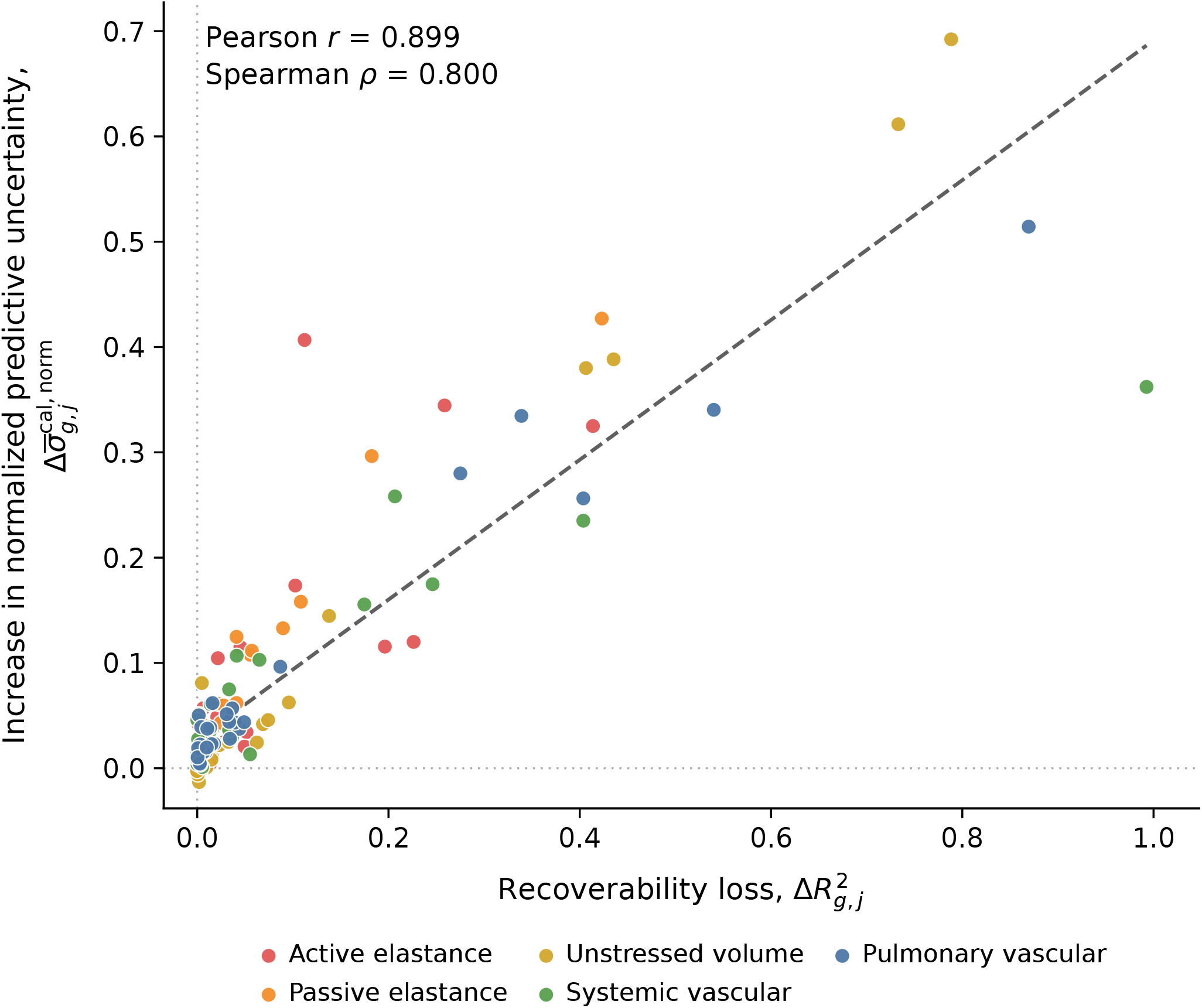
Relationship between parameter-level recoverability loss 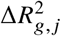 and the corresponding change in normalized calibrated predictive uncertainty 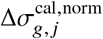 under structured measurement removal. Each point represents one masked-measurement-group–parameter pair. Colors denote LPN parameter groups, and the dashed line indicates the least-squares trend.

The fixed observation scenarios reinforced the same conclusion (Table 4). Basic clinical measurements (systemic systolic/diastolic pressure and heart rate) yielded macro *R*^2^ = 0.066. BP + chamber volumes, Echo-like, and hemodynamic sets produced macro *R*^2^ values of 0.389, 0.374, and 0.394, respectively. The rich clinical set reached 0.751, compared with 0.842 for all 30 measurements. The Echo-like set contained more measurements than the BP + chamber-volume set but performed slightly worse in aggregate, consistent with the importance of pressure information demonstrated by the structured-removal experiment. The atrial active elastances remained poorly recovered even under the full synthetic observation set.

**Table 4.**
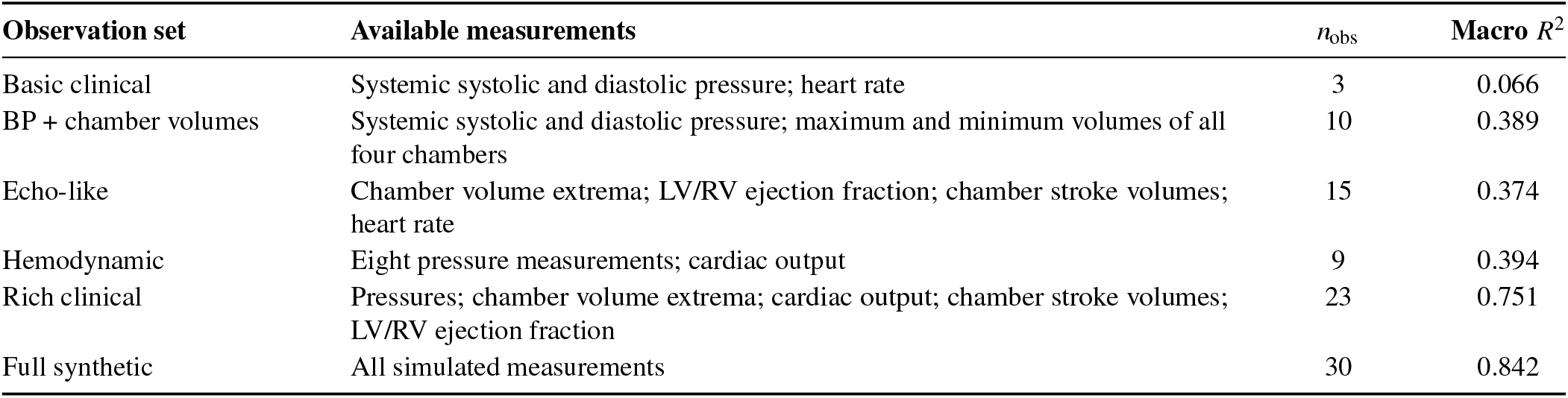
Performance of mask-aware Mada across six fixed, clinically motivated observation sets. Here, *n*_obs_ denotes the number of available measurements in each observation set. No scenario-specific retraining or fine-tuning was performed. The full-synthetic row represents mask-aware Mada evaluated with all measurements available and is distinct from the dedicated full-observation Mada model reported in Table 2.

| Observation set | Available measurements | $n_{\text{obs}}$ | Macro $R^2$ |
| --- | --- | --- | --- |
| Basic clinical | Systemic systolic and diastolic pressure; heart rate | 3 | 0.066 |
| BP + chamber volumes | Systemic systolic and diastolic pressure; maximum and minimum volumes of all four chambers | 10 | 0.389 |
| Echo-like | Chamber volume extrema; LV/RV ejection fraction; chamber stroke volumes; heart rate | 15 | 0.374 |
| Hemodynamic | Eight pressure measurements; cardiac output | 9 | 0.394 |
| Rich clinical | Pressures; chamber volume extrema; cardiac output; chamber stroke volumes; LV/RV ejection fraction | 23 | 0.751 |
| Full synthetic | All simulated measurements | 30 | 0.842 |

Restricted fixed observation sets also increased calibrated predictive uncertainty. Mean normalized predictive standard deviation was highest for the basic clinical scenario and decreased substantially for the rich clinical and full synthetic observation sets. Scenario-specific validation calibration maintained near-nominal interval coverage in most settings, although the basic clinical set retained undercoverage. Calibration details are summarized in Appendix B.3.

### 3.3 Parameter Recovery Is Distinct From Hemodynamic Reconstruction

Across the 20-case held-out diagnostic cohort, selected TRF achieved mean cross-case *R*^2^ = 0.588 over seven selected difficult parameters 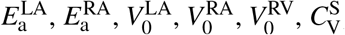and 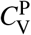, compared with 0.601 for full-observation Mada, 0.194 for Dogbox, and −0.001 for L-BFGS-B. For each optimizer and case, the result with the lower forward NRMSE from two fixed initializations was selected, without reference to parameter error. The median forward NRMSE was 0.002 for selected TRF versus 0.054 for Mada, while the corresponding median mean range-normalized parameter errors were 0.069 and 0.075, respectively. Thus, TRF achieved substantially better hemodynamic reconstruction while providing broadly comparable parameter recovery on this cohort. All 40 TRF runs converged; detailed optimizer and runtime results are provided in Appendix C.2.

Atrial active-elastance recovery remained heterogeneous. LA and RA cross-case *R*^2^ values were 0.236 and 0.352 for selected TRF, compared with 0.139 and 0.441 for Mada. Figure 6 separates the forward and parameter errors.

**Figure 6.**
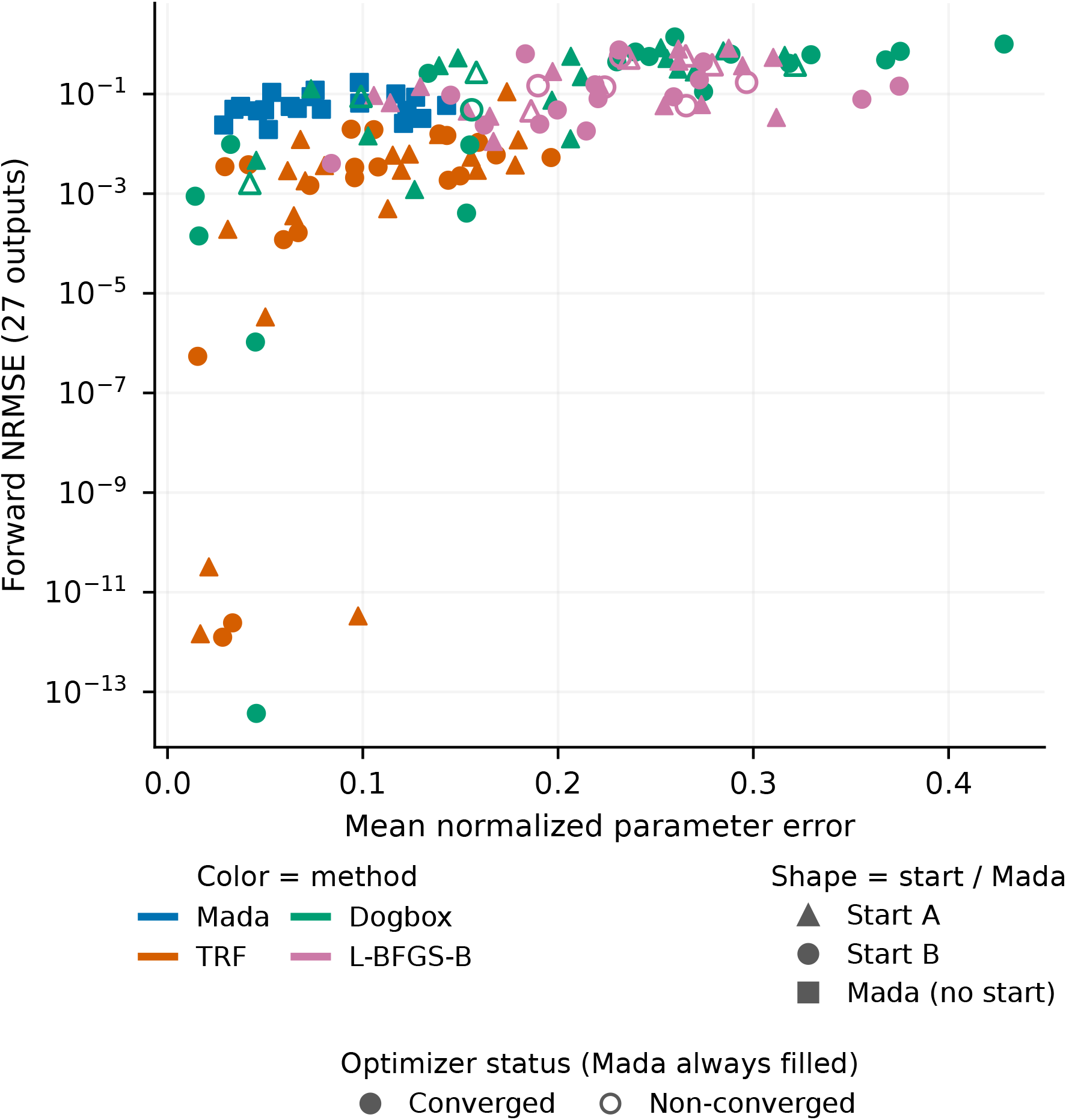
Parameter error and original LPN forward-solver NRMSE for 20 cases: all 20 Mada estimates and both starts for each optimizer (40 runs each). Colors identify methods. Triangles denote start A; circles start B; filled squares Mada (no starts). Filled/open optimizer markers indicate convergence/non-convergence. Parameter errors use generating ranges. Mada is the dedicated full-observation model (Appendix C.1).

The distinction between hemodynamic reconstruction and parameter recovery is also evident at the individual-case level. Cases 15 and 06 exhibited relatively small forward NRMSE values of 0.011 and 0.002, respectively, while their mean range-normalized parameter errors were 0.159 and 0.144. The largest atrial active-elastance errors reached 0.460 and 0.858, respectively. These results illustrate that relatively close hemodynamic reconstruction can coexist with appreciable parameter-recovery errors.

Target-specific single-output fine-tuning provided little improvement in atrial active-elastance prediction. Predicting each target independently increased full-test *R*^2^ by only 0.002 for 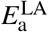 and 0.007 for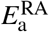, suggesting that their poor recovery was not primarily caused by interference among the 20 simultaneously predicted targets.

The retained Case 07 profiles showed the broadest low-error compensation for RA active elastance. At forward NRMSE ≤ 0.02, 6/7 RA grid positions qualified, compared with 1/7 LA positions and 4/7 LV-control positions. Four of 25 joint LA–RA combinations qualified, with minimum forward NRMSE 0.003 (Fig. 7).

**Figure 7.**
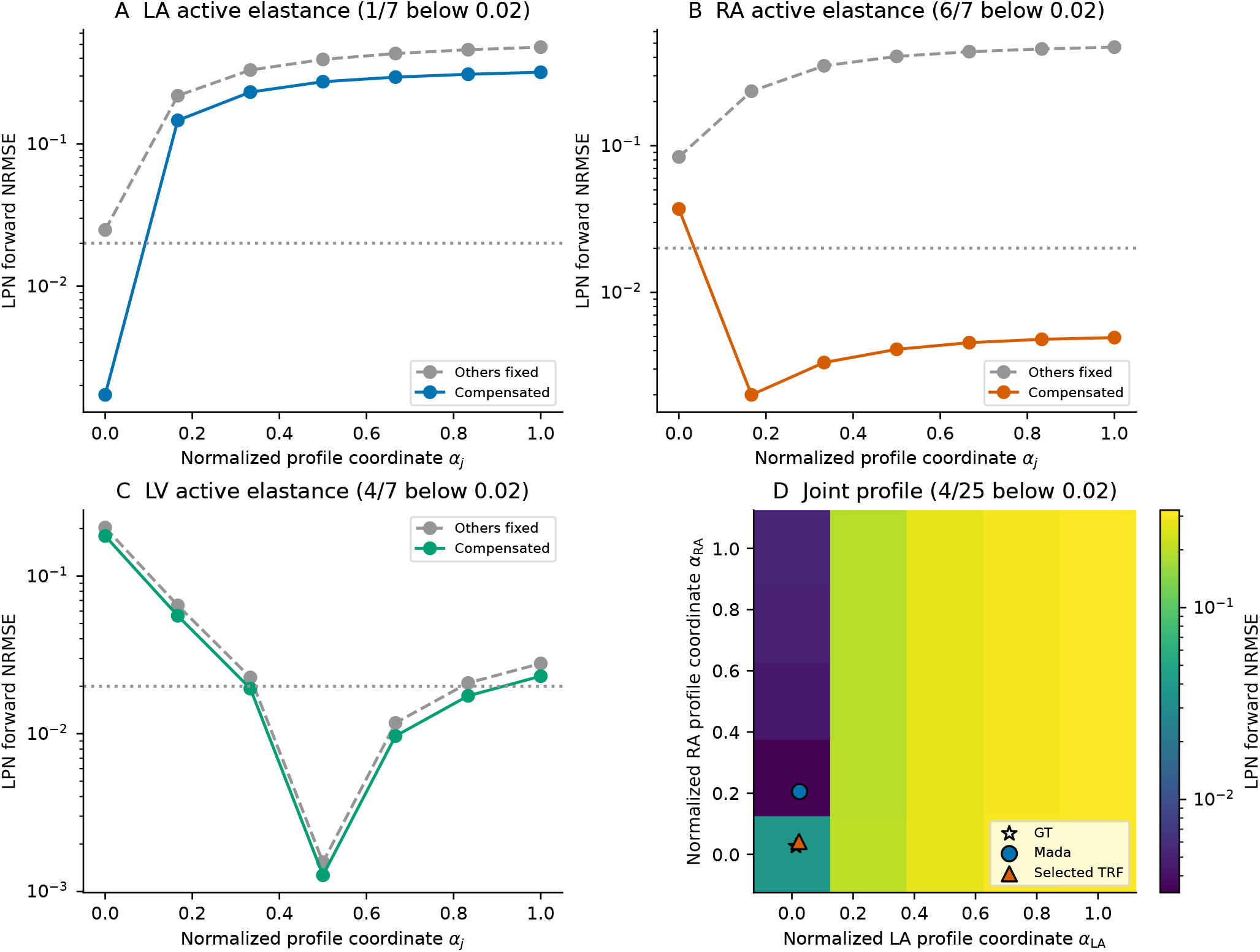
Case 07 compensation profiles. Panels A–C compare fixed and locally compensated LA, RA, and LV active elastance; the NRMSE threshold 0.02 is met at 1/7, 6/7, and 4/7 sampled positions. Panel D shows the joint LA–RA grid (4/25 qualifying points). The normalized profile coordinates *α*_*j*_ span fixed physical grid bounds, whereas free-parameter optimization uses generating bounds.

The five normalized forward Jacobians were strongly ill-conditioned: at *h* = 0.01, condition numbers ranged from 1.91 × 107 to 1.33 × 109, with effective ranks 15–17 of 20. Effective ranks were unchanged across the tested steps, although some weakest directions were step-sensitive. The spectra and detailed loadings are reported in Appendix D.6 and Tables 18–20.

### 3.4 Robustness to Measurement Noise and Model Architecture

The measurement-noise amplitude sweep showed a progressive distribution shift as the prescribed perturbations increased. At *γ*_noise_ = 0, 0.25, 0.5, and 1.0, macro *R*^2^ decreased from 0.862 to −0.053, −0.749, and −1.610, while macro NRMSE increased from 0.276 to 0.959, 1.279, and 1.586. Uncertainty calibration deteriorated substantially: the mean Std./RMSE ratio decreased from 1.046 to 0.281, 0.226, and 0.190, and PICP_95_ decreased from 0.974 to 0.474, 0.386, and 0.310. Thus, uncertainty calibrated on clean synthetic observations did not remain calibrated after the noise-induced input shift (Table 5; Appendix D.5).

**Table 5.** Performance of the frozen full-observation Mada model with increasing measurement-noise amplitude. Estimates are projected to training extrema; uncertainty retains clean-validation calibration.

| $\gamma_{\text{noise}}$ | Macro $R^2$ | Macro NRMSE | Std./RMSE | PICP<sub>95</sub> |
| --- | --- | --- | --- | --- |
| 0 | 0.862 | 0.276 | 1.046 | 0.974 |
| 0.25 | -0.053 | 0.959 | 0.281 | 0.474 |
| 0.50 | -0.749 | 1.279 | 0.226 | 0.386 |
| 1.00 | -1.610 | 1.586 | 0.190 | 0.310 |

**Table 6.**
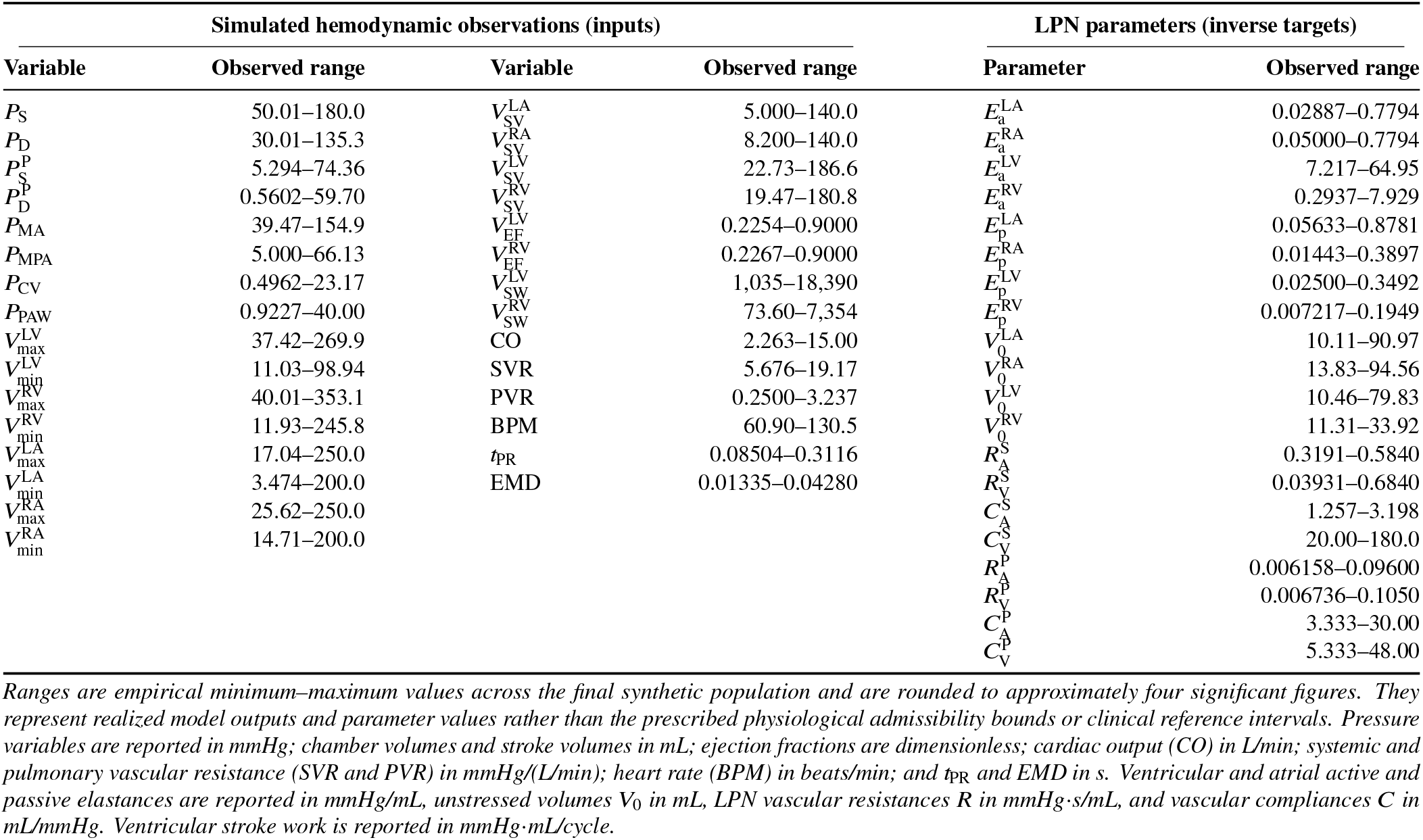
Observed ranges of the 30 simulated hemodynamic observations used as model inputs and the 20 LPN parameters used as inverse-prediction targets in the final 650,000-sample synthetic cardiovascular dataset.

In the five-case original-LPN noise diagnostic, median clean-reference forward NRMSE increased from 0.002 under clean inversion to 0.264 after noisy inversion. The prescribed perturbations therefore changed both the fitted parameters and their agreement with the clean generating observations.

Using the same measurement-availability conditions evaluated in Sec. 3.2, mask-aware Mada achieved macro *R*^2^ = 0.842 with complete observations and 0.337 at 70% random missingness, compared with 0.794 and 0.312, respectively, for the simple probabilistic ResMLP (Fig. 8). Across the same fixed observation sets, both models improved as observation content became richer, although the ordering of the intermediate sets differed. Parameter-wise differences were heterogeneous, and the uncertainty metrics did not consistently favor either model.

**Figure 8.**
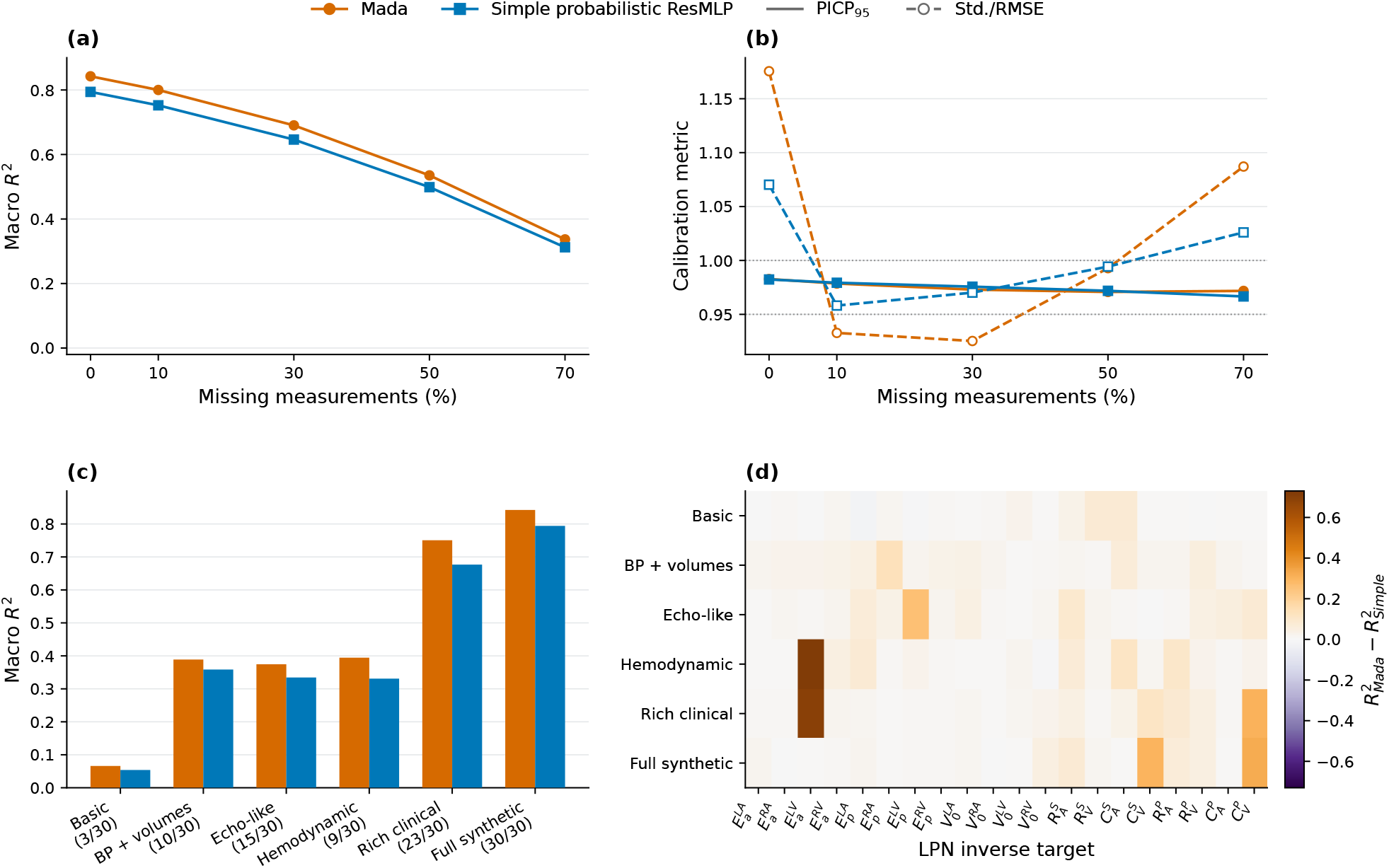
Comparison between mask-aware Mada and a simple mask-aware probabilistic ResMLP. (a) Macro *R*^2^ under random missingness. (b) Uncertainty calibration. (c) Macro *R*^2^ across fixed observation sets. (d) Parameter-wise *R*^2^ difference.

## 4 Discussion

This study investigated uncertainty-aware inverse estimation of cardiovascular LPN parameters under variable measurement availability. The results show that inverse recoverability is strongly parameter dependent and measurement dependent, and that predictive uncertainty, direct optimization, and local sensitivity analysis provide complementary views of this difficulty. In particular, accurate hemodynamic reconstruction does not necessarily imply accurate recovery of the generating physiological parameters.

### 4.1 Information-Limited and Inference-Limited Recovery

The direct-optimization results distinguish hemodynamic consistency from parameter recoverability. TRF achieved substantially smaller forward residuals than neural inference while providing broadly comparable parameter recovery on the diagnostic cohort. However, close agreement with the observed hemodynamic summaries did not always imply recovery of the generating parameter vector. In general, distinct parameter vectors may satisfy

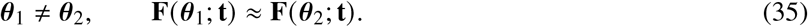

The profile and Jacobian analyses provide complementary evidence for this behavior. Some parameters could be displaced substantially while compensatory changes in other parameters preserved similar modeled hemodynamics, and the normalized forward Jacobians contained weak local parameter directions. At the same time, strong parameter recovery remained possible for other cases. Together with the limited improvement from target-specific single-output fine-tuning, these findings support heterogeneous, case-dependent inverse difficulty rather than a uniform failure of either neural inference or direct optimization.

These diagnostics should not be interpreted as establishing global structural non-identifiability. The profile analyses explored a finite grid and a targeted compensation subset, while the weakest local Jacobian directions showed some sensitivity to the finite-difference step. Rather, they provide case-specific evidence of practical ambiguity within the evaluated observation space and help explain why some parameters remain difficult to recover even when forward reconstruction is accurate. Future work could combine the present recoverability- and uncertainty-aware framework with generative inverse methods, such as InVAErt [33], to represent multiple compatible parameter combinations more explicitly.

Among the traditional optimizers evaluated here, TRF was the most reliable, whereas Dogbox and L-BFGS-B showed greater sensitivity to numerical termination and initialization. Neural and optimization-based inference therefore offer different advantages. Neural inference amortizes the cost of learning across a population and provides rapid parameter estimates together with predictive uncertainty, whereas direct optimization performs case-specific residual minimization through repeated LPN evaluations. Neither low forward residual nor strong average population-level prediction alone establishes that an individual physiological parameter is tightly constrained. Accurate hemodynamic reconstruction can nevertheless be sufficient when parameter identification is not the modeling objective.

### 4.2 Measurement Content and Implications for Personalization

The measurement-availability experiments show that recoverability depends on physiological information content rather than measurement count alone. Pressure measurements were particularly important for vascular parameters, chamber volumes strongly informed unstressed chamber volumes, and stroke-work information contributed substantially to ventricular active-mechanics recovery. The effects also differed among parameters within the same physiological group, indicating that measurement requirements should be considered at the parameter level rather than only by broad model category.

The fixed observation scenarios reinforce this interpretation. Sparse observation sets did not support uniformly reliable recovery across all 20 parameters, and adding measurements was most useful when the added information constrained previously weak aspects of the inverse problem. The rich clinical observation set substantially improved recovery of many ventricular and vascular parameters, yet the atrial active elastances remained difficult even with the complete synthetic observation set. Additional measurements of the same type may therefore provide limited benefit for these parameters. Measurements more directly related to atrial contraction, such as atrial pressure–volume behavior, atrial strain, or transvalvular flow, may be more informative targets for future study.

Structured measurement removal should also be interpreted conditionally. A small recoverability loss after removal of a measurement group does not imply that the measurements are physiologically unimportant; their information may be redundant with measurements that remain available. Conversely, a large loss indicates that the retained observations provide insufficient replacement information for that parameter. These experiments therefore characterize empirical measurement–parameter dependencies rather than causal relationships or formal structural-identifiability properties.

For LPN personalization, the practical question is consequently not only how to estimate a parameter, but whether the available observations support its estimation. Parameter-specific uncertainty may help identify parameters that can be estimated with confidence, those that require stronger regularization or prior information, and those for which a range of compatible values may be more appropriate than a single estimate. The present uncertainty estimates, however, were calibrated within the synthetic population and should not yet be interpreted as clinical confidence intervals. Because the mask-aware model was trained primarily with randomly generated measurement masks, the absolute degradation observed under structured and fixed clinical masks may reflect both physiological information loss and mask-pattern distribution shift. The within-count contrasts nevertheless demonstrate that measurement identity carries information beyond measurement number alone.

### 4.3 Predictive Uncertainty, Measurement Noise, and the Role of Architecture

Under clean observations, validation-based calibration provided a useful parameter-specific uncertainty scale, and reduced measurement availability was generally accompanied by both poorer point prediction and greater reported uncertainty.

Missing and noisy measurements had distinct effects on inverse prediction. Under missingness, the mask-aware model explicitly represented unavailable observations, and validation-calibrated uncertainty maintained broadly stable interval coverage despite declining parameter-recovery accuracy. In contrast, measurement noise altered observed values without explicitly indicating their reduced reliability to the model, leading to substantial degradation in both parameter recovery and prediction-interval coverage. These findings highlight the importance of considering both measurement availability and measurement quality in uncertainty-aware cardiovascular LPN personalization.

The architecture diagnostics further clarify the role of the Mada-specific formulation. The validation-derived recoverability scores were highly informative descriptors of parameter-wise inverse difficulty and generalized strongly to held-out data. Weak recoverability-weighted reference regularization produced a modest improvement in the matched full-observation sweep, with stronger regularization providing no additional benefit. The sweep varied regularization strength while keeping *q*_*j*_ fixed, so it does not establish an advantage over uniform reference weighting. The reference penalty acts as a weak target-specific inductive prior: well-recoverable parameters are more strongly anchored to the reference estimate, whereas poorly recoverable parameters retain greater flexibility.

The simpler probabilistic ResMLP reproduced the major parameter-difficulty patterns and the degradation associated with reduced measurement availability. This supports the interpretation that the principal physiological trends arise from the observation-to-parameter inverse problem rather than from the Mada architecture itself.

### 4.4 Limitations and Future Directions

The present study used a controlled synthetic population generated and evaluated with the same cardiovascular LPN, enabling direct assessment of parameter recovery but limiting generalization beyond the assumed model and parameter distributions. Differences in cardiac activation, valve behavior, vascular representation, and physiological parameter ranges could alter parameter sensitivities, compensatory relationships, and predictive uncertainty. Furthermore, the fixed observation scenarios approximate clinical measurement availability without fully capturing variations in acquisition conditions and measurement quality. Future work should examine whether the observed recoverability patterns persist across disease-specific populations, more detailed cardiovascular models, including whole-circulation formulations such as those of Mynard and Smolich [3], and independent forward models and patient data.

The measurement-noise analysis used a prescribed perturbation model and a single noise realization. Future studies could extend this analysis to repeated noise realizations and more realistic, acquisition-specific measurement errors. Mada further assumes conditionally independent Gaussian predictive distributions for the inferred parameters. This representation does not capture posterior dependence, multimodality, or bounded and asymmetric inverse distributions. The profile analyses indicate that compensatory parameter changes can occur along coupled directions in parameter space, suggesting that multivariate or multimodal probabilistic formulations may better represent weakly constrained solutions.

Finally, the direct-optimization and local-sensitivity analyses were diagnostic rather than exhaustive. The optimization benchmark used a finite 20-case cohort, two fixed initializations, and local optimization algorithms, while the profile analysis examined a targeted compensation subset for a single retained case. More extensive exploration was limited by the computational cost of repeated LPN forward evaluations required for case-wise optimization and profile searches. These experiments therefore demonstrate practical ambiguity in selected regions of the inverse problem but do not characterize the complete solution set or establish population-wide structural non-identifiability. Broader population-level sensitivity analysis and more systematic exploration of compatible parameter sets would provide a natural extension as computational efficiency improves.

## 5 Conclusion

We developed Mada, a mask-aware probabilistic neural framework, to investigate parameter-specific recoverability and predictive uncertainty in cardiovascular LPN inverse inference under variable measurement availability. Our results demonstrated substantial heterogeneity in parameter recoverability, with several ventricular and vascular parameters recovered accurately while atrial active elastances remained comparatively difficult. Measurement content, rather than measurement count alone, strongly influenced parameter recovery, highlighting the importance of selecting physiologically informative observations. Complementary direct-optimization and sensitivity analyses showed that close hemodynamic reconstruction did not always guarantee accurate recovery of the generating parameters, with case-specific parameter compensation and weak observational directions contributing to inverse difficulty. Predictive uncertainty generally increased as informative measurements were removed, while measurement noise degraded both parameter recovery and prediction-interval coverage. Similar recoverability patterns observed with a simpler probabilistic ResMLP further supported the broader relevance of these findings across neural architectures. Overall, this study highlights the importance of integrating parameter-wise recoverability, measurement sufficiency, and predictive uncertainty to support more reliable cardiovascular LPN personalization.

## Data and Code Availability

Upon acceptance, the synthetic dataset and neural-network model code supporting this study will be made publicly available through a GitHub repository.

## Acknowledgment

Research reported in this publication was supported by the National Heart, Lung, and Blood Institute of the National Institutes of Health under Award No. R15HL181637. This material is also based upon work supported by the U.S. National Science Foundation under Award No. 2501698 and by the Kennesaw State University Grand Challenge Award. The content is solely the responsibility of the authors and does not necessarily represent the official views of the National Institutes of Health.

## Funding Data

- National Heart, Lung, and Blood Institute, National Institutes of Health (NIH), Award No. R15HL181637.
- U.S. National Science Foundation (NSF), Award No. 2501698.
- Kennesaw State University, Grand Challenge Award.

## A Cardiovascular LPN and Synthetic Population

### A. Cardiovascular LPN Governing Equations and Inferred Parameterization

The synthetic inverse-learning dataset was generated using the closed-loop cardiovascular lumped-parameter network (LPN) described below, in which the four cardiac chambers are coupled to systemic and pulmonary arterial and venous compartments. The same executable forward solver and fixed model configuration were retained for the physics-based diagnostic analyses in this study. The model follows the standard electrical-analogue formulation: pressure corresponds to voltage, flow to current, vascular resistance to a resistive element, compliance to a capacitive element, and inertance to an inductive element [1–3]. Cardiac chambers are represented by time-varying elastance elements and the four cardiac valves by pressure-dependent resistive elements. The overall network structure is shown in Fig. 1.

#### A.1.1 Closed-Loop Governing Equations

Let *V*_*I*_ (*t*) and *P*_*I*_ (*t*) denote the volume and pressure of chamber *I ∈* {LA, LV, RA, RV}. Conservation of volume in the four chambers gives

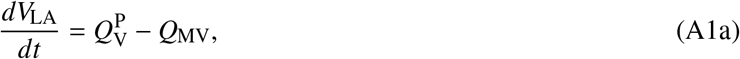

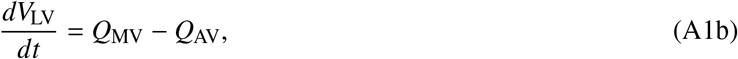

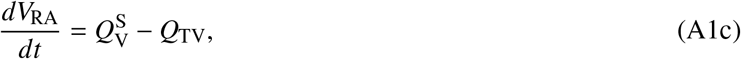

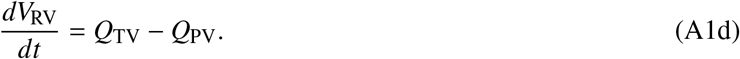

The systemic and pulmonary vascular compliances satisfy

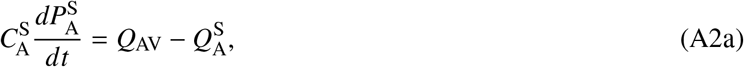

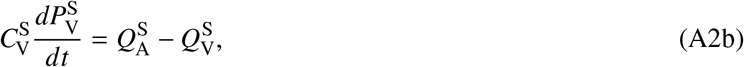

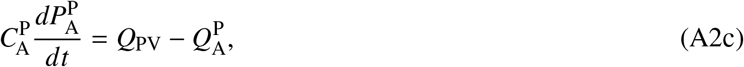

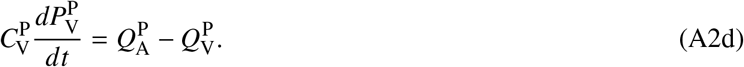

Flow through the systemic and pulmonary vascular segments is governed by the corresponding resistance–inertance relation,

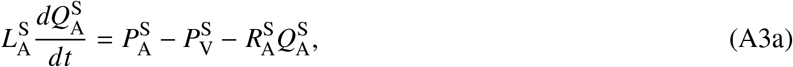

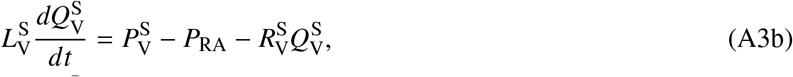

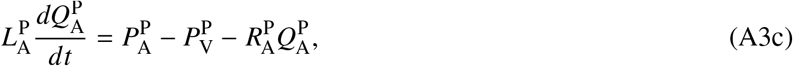

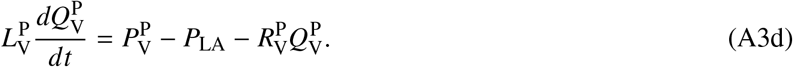

Here *Q, P*, and *V* denote flow rate, pressure, and volume, respectively, while *R, C*, and *L* denote vascular resistance, compliance, and inertance. The subscripts A and V denote arterial and venous vascular compartments.

The mitral (MV), aortic (AV), tricuspid (TV), and pulmonary (PV) valve flows are written as

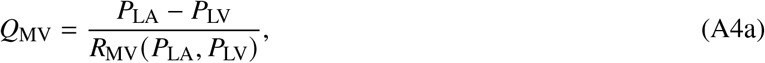

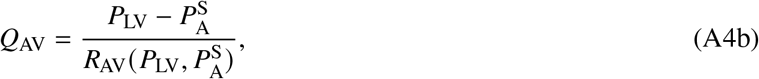

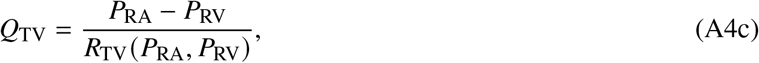

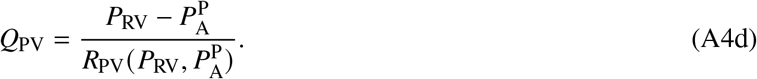

For valve *i*, the effective resistance is

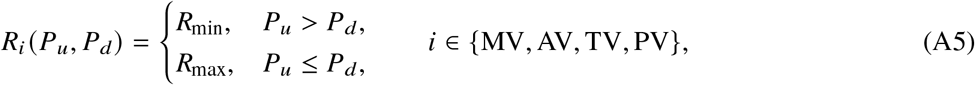

where *P*_*u*_ and *P*_*d*_ denote the upstream and downstream pressures, respectively. The quantities *R*_min_ and *R*_max_ denote the low-resistance open-valve state and high-resistance closed-valve state, respectively. This switching relation provides a reduced-order representation of valve opening and closure.

#### A.1.2 Time-Varying Chamber Elastance

The pressure in each cardiac chamber is determined by

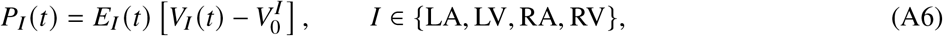

where 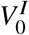 is the chamber unstressed volume. The time-varying elastance is decomposed into passive and active contributions,

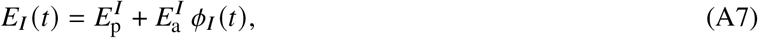

where 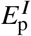 characterizes passive chamber stiffness, 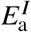 controls the active contractile contribution, and *ϕ*_*I*_ (*t*) is a normalized activation function.

The forward LPN uses a double-Hill chamber-activation form [44],

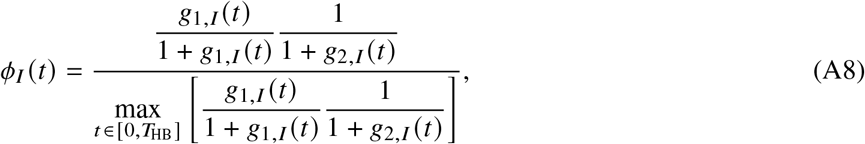

with

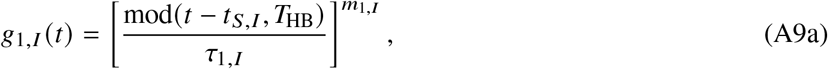

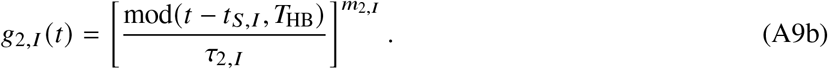

The cardiac-cycle duration is *T*_HB_ = 60/BPM. The parameters *t*_*S,I*_, *τ*_1,*I*_, *τ*_2,*I*_, *m*_1,*I*_, and *m*_2,*I*_ determine chamber-specific activation timing and shape. In the inverse experiments, BPM, *t*_PR_, and EMD were treated as known case-specific timing quantities rather than unknown LPN parameters. Thus, they were included in the 30-variable observation vector used by the neural models and retained as fixed inputs in the direct LPN optimization benchmark.

#### A.1.3 Inferred Parameterization

The inverse problem estimates a selected 20-dimensional subset of the LPN parameters. The chamber-related targets consist of four active elastances,

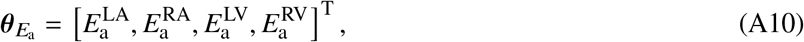

four passive elastances,

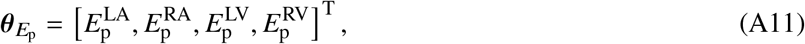

and four unstressed chamber volumes,

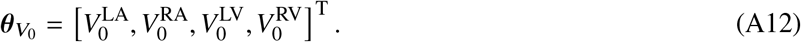

The systemic vascular targets are

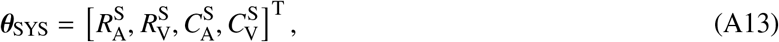

and the pulmonary vascular targets are

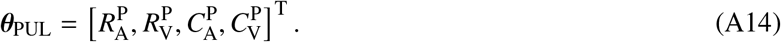

The complete inverse target vector is therefore

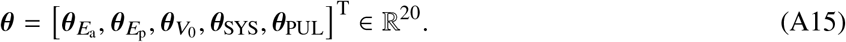

Valve resistances, vascular inertances, and activation-shape parameters were part of the forward LPN but were not included among the 20 inverse targets. This distinction is important because the recoverability results in the main text concern only the selected chamber-mechanics and systemic/pulmonary resistance–compliance parameters defined above.

### A.2 Characterization of the Synthetic Parameter Population

The inverse-learning experiments used a synthetic cardiovascular population of 650,000 cases. Its realized parameter and observation distributions are summarized below because empirical recoverability depends on the domain over which prediction is evaluated.

#### A.2.1 Realized Parameter Space

Table 6 summarizes the observed ranges of the simulated hemodynamic observations used as model inputs and the LPN parameters used as inverse-prediction outputs.

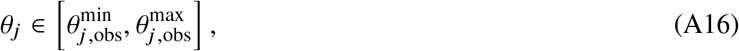

with observed midpoint

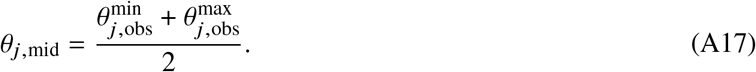

The midpoint summarizes the observed range and should not be interpreted as a physiological reference value.

The observed ranges define the realized parameter domain used for training, validation, and testing. Because regression difficulty depends on this domain, parameter-wise recoverability should be interpreted as empirical predictability within the present synthetic population rather than as an intrinsic property of each physiological parameter.

#### A.2.2 Correlation Structure of Synthetic Inputs and Parameter–Input Associations

Figure 9 reports input–input and parameter–input Spearman correlations. It describes redundancy among measurements and marginal associations between individual parameters and observations. These pairwise correlations do not establish independence, causality, or identifiability, because important nonlinear and multivariable relationships may remain.

**Figure 9.**
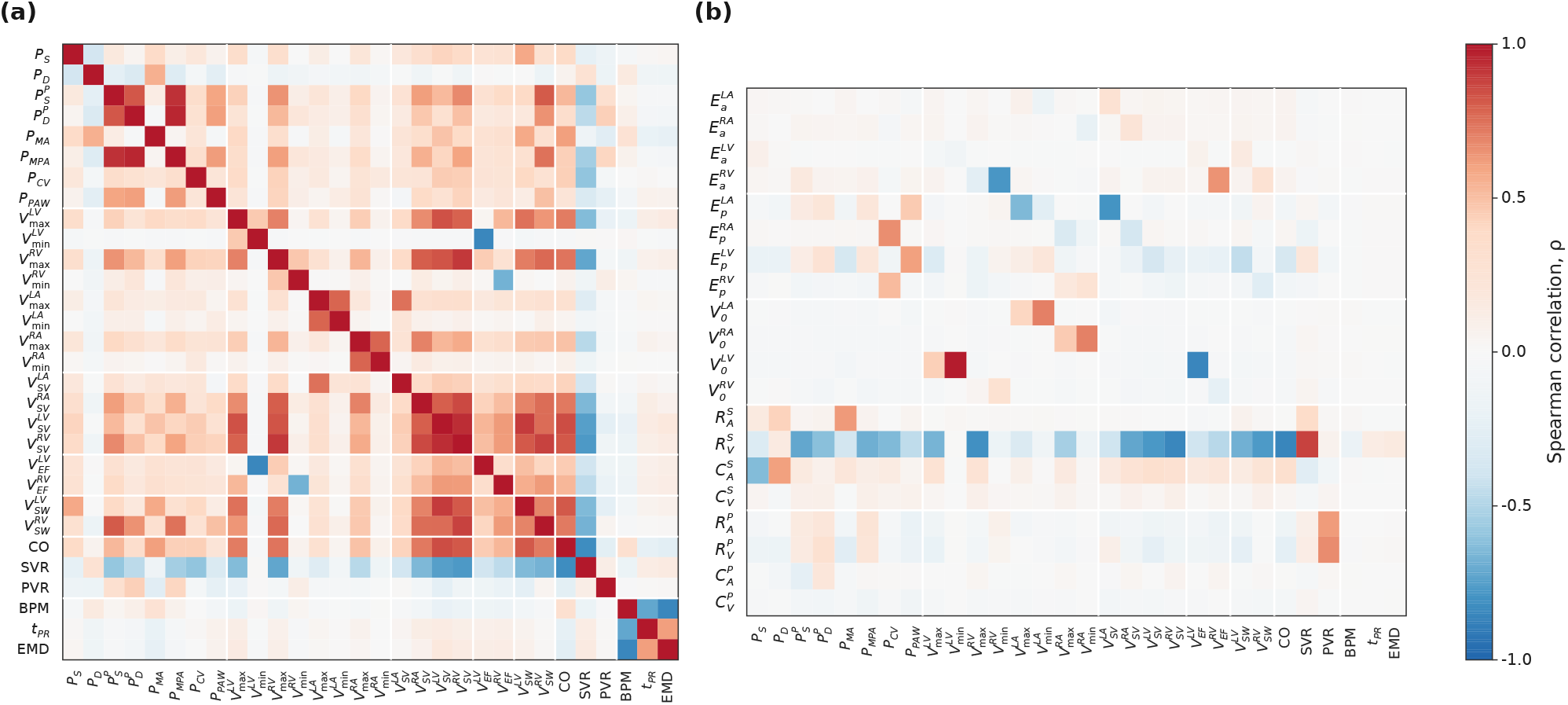
Correlation structure of the synthetic inverse-learning dataset. (a) Pairwise Spearman correlations among the 30 hemodynamic measurements used as model inputs. (b) Spearman correlations between the 20 LPN parameters (rows) and the 30 hemodynamic inputs (columns).

**Figure 10.**
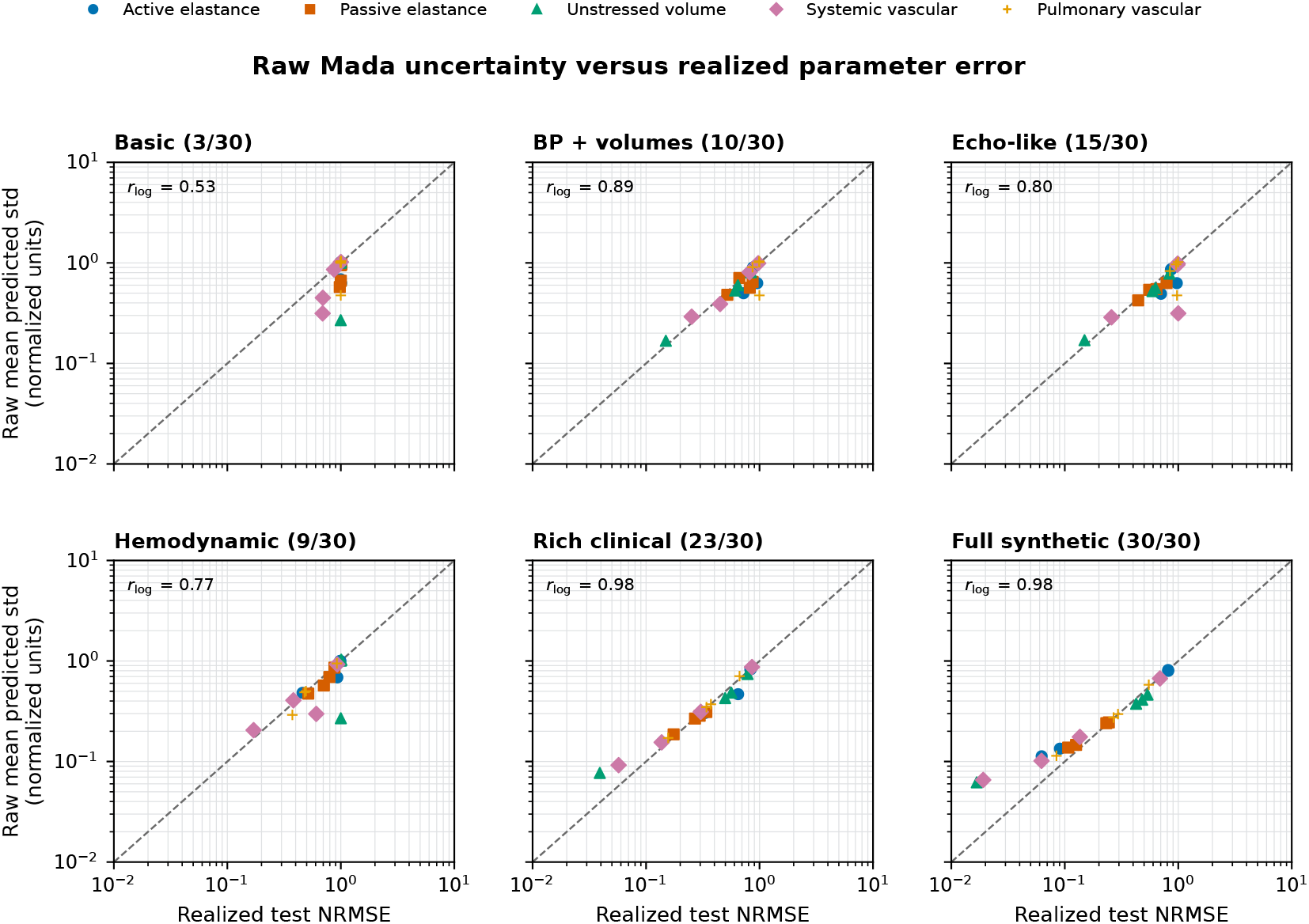
Correspondence between raw Mada predictive uncertainty and realized parameter-recovery error under fixed clinical observation sets. Each point represents one of the 20 inverse targets. The reported *r*_log_ is the Pearson correlation after logarithmic transformation of both quantities.

**Figure 11.**
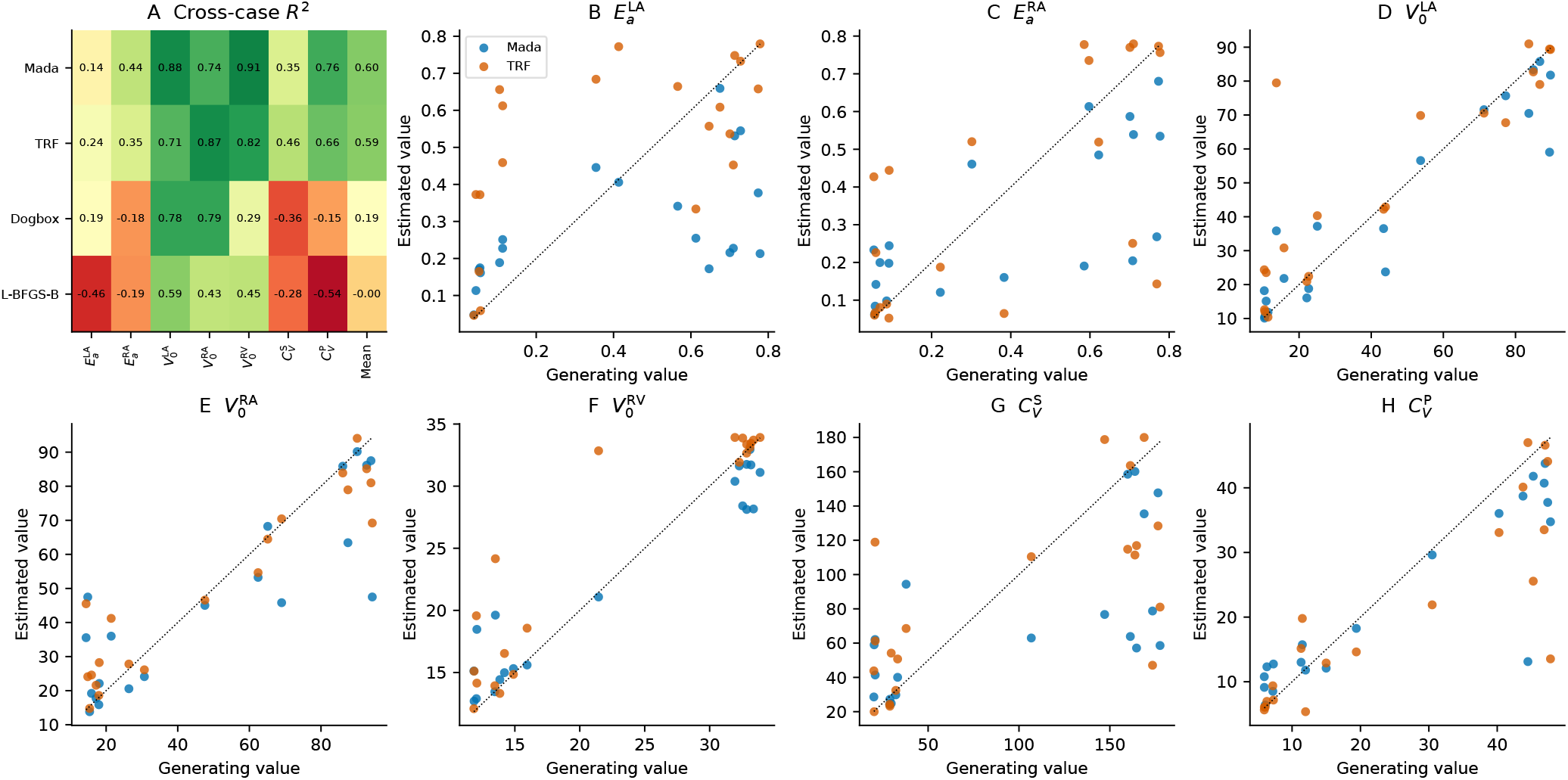
Cross-case recovery of seven emphasized parameters and their macro mean (A); generating values versus Mada and selected TRF estimates (B–H). For each optimizer, a start is selected solely by forward NRMSE.

**Figure 12.**
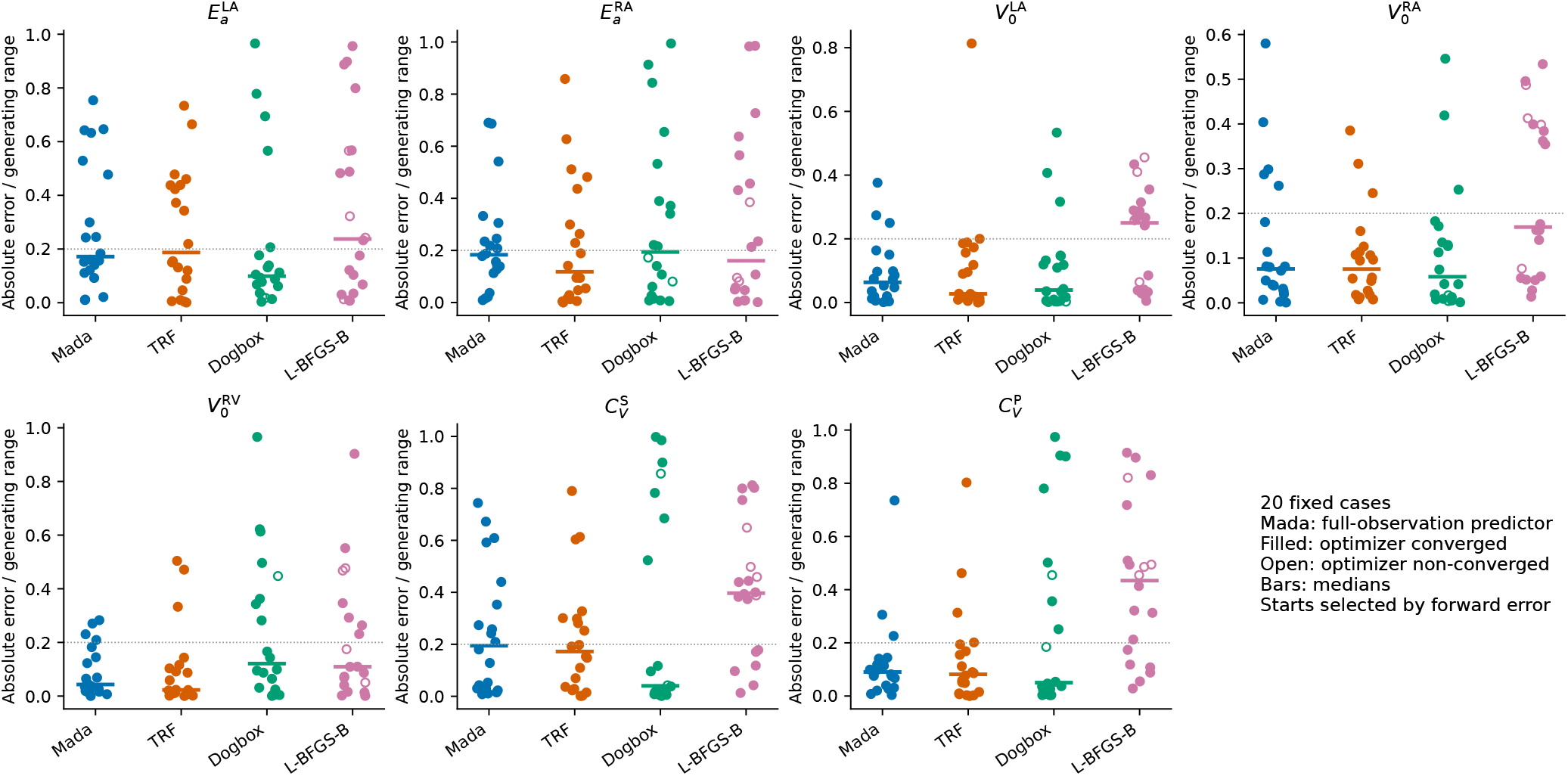
Range-normalized absolute errors for the seven emphasized targets. Filled markers denote optimizer convergence; open markers denote non-convergence from an evaluation limit or line-search failure. Horizontal bars show medians. Mada is the dedicated full-observation model. The 0.2 reference line is descriptive.

**Figure 13.**
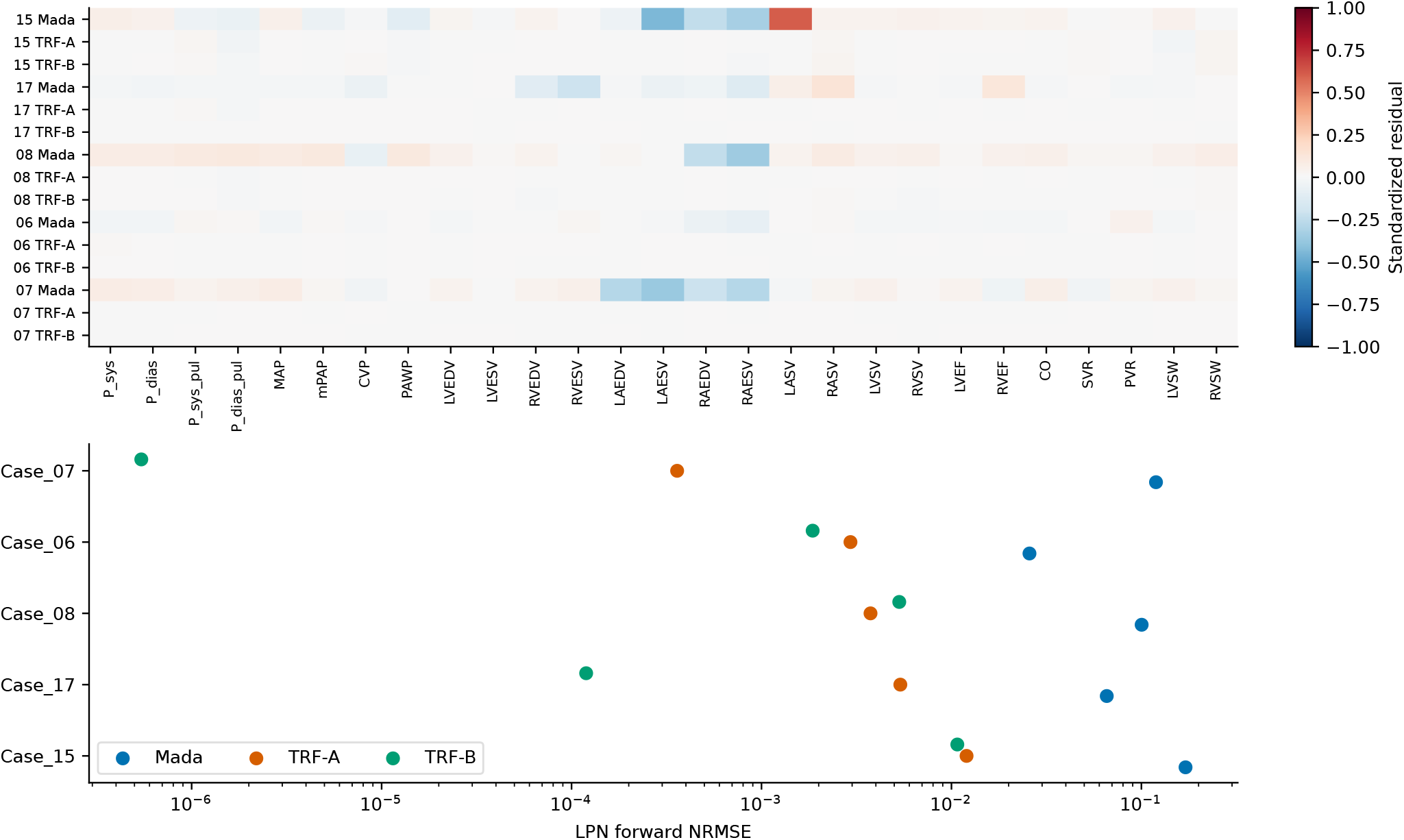
Five-case original LPN forward-solver comparison. Upper: standardized residuals of the 27 summary outputs. Lower: forward NRMSE for Mada and both optimized TRF starts.

**Figure 14.**
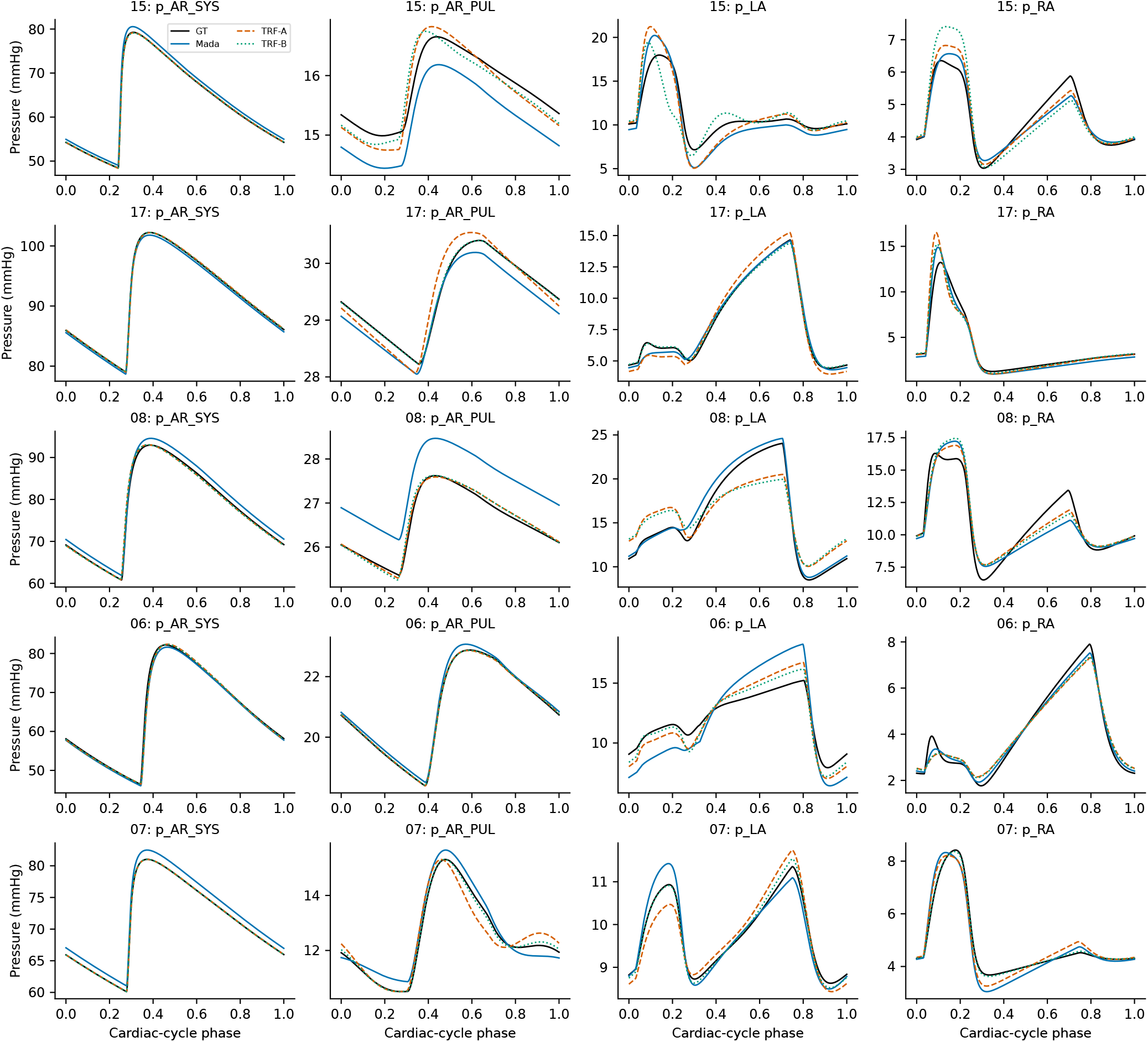
Final-cycle pressure comparisons for Cases 15, 17, 08, 06, and 07. Columns show systemic arterial, pulmonary arterial, LA, and RA pressures. GT denotes original-solver replay at generating parameters; the other curves use Mada and optimized TRF-A/B parameters.

**Figure 15.**
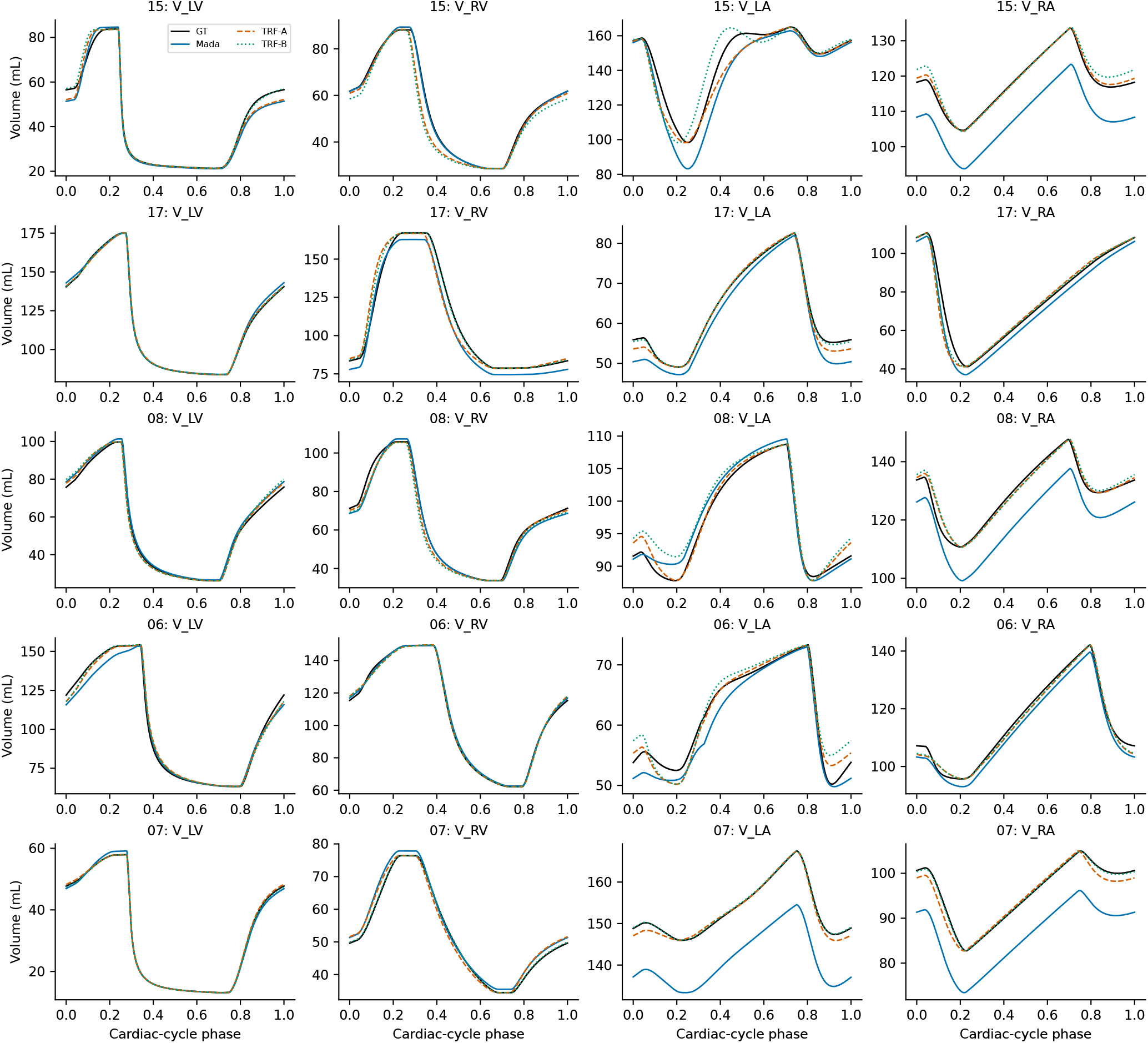
Final-cycle chamber-volume comparisons for the same five cases and parameter vectors. Columns show LV, RV, LA, and RA volume.

#### A.2.3 Implications for Interpretation of Recoverability

Conceptually, empirical recoverability can be viewed as

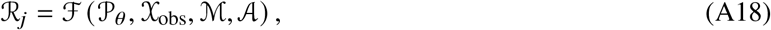

where *P*_*θ*_ denotes the parameter population, *X*_obs_ the observation space, *M* the mechanistic LPN, and *A* the inference procedure. Changing the parameter ranges or physiological dependence structure could alter both absolute error and the relative ranking of difficult and easy targets. The recoverability values reported here are therefore conditional on the synthetic population and should not be interpreted as universal structural-identifiability properties.

### A.3 Baseline Neural Architectures

To provide additional implementation details for the neural baselines reported in Table 2, we summarize their main architectural characteristics in Table 7. All models received the same 30-dimensional normalized full-observation input vector and predicted the same 20 normalized LPN parameters. The deterministic baselines produced point estimates directly, whereas the Gaussian-NLL model additionally predicted parameter-wise variances and was optimized using a heteroscedastic Gaussian negative log-likelihood. These architectures were used as representative neural baselines rather than as an exhaustive architecture search.

**Table 7.**
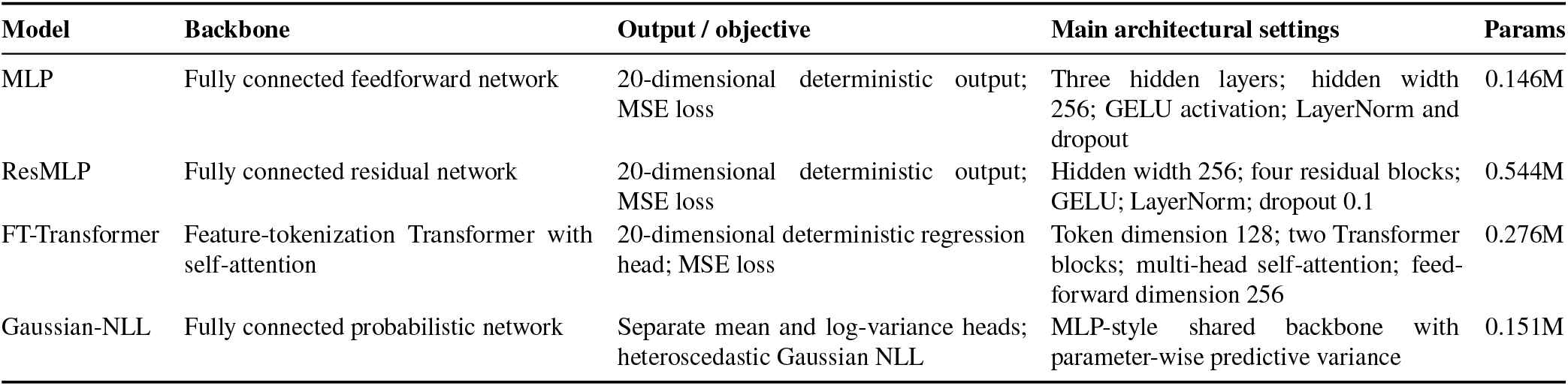
Architectural details of the non-Mada neural baselines used in the full-observation comparison in Table 2. All models used the same 30 normalized hemodynamic inputs and 20 normalized inverse targets.

**Table 8.**
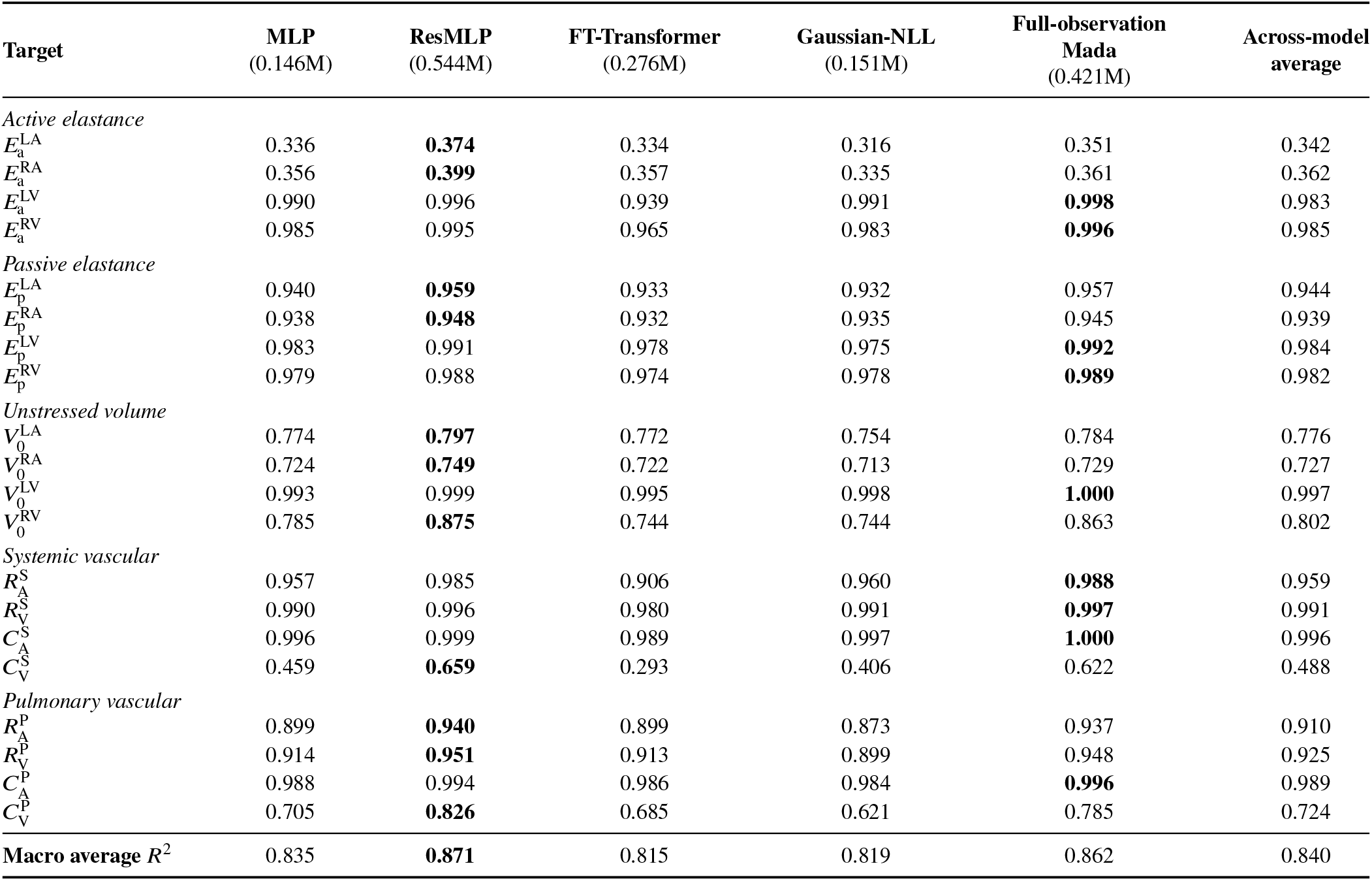
Cross-model parameter-wise inverse-prediction performance under complete observations. The Mada column corresponds to the dedicated full-observation Mada model rather than the mask-aware model evaluated with a complete mask. Entries are held-out test-set *R*^2^ values; boldface indicates the best result for each target. Numbers in parentheses below the model names denote trainable parameter counts in millions (M).

| Target | MLP<br>(0.146M) | ResMLP<br>(0.544M) | FT-Transformer<br>(0.276M) | Gaussian-NLL<br>(0.151M) | Full-observation<br>Mada<br>(0.421M) | Across-model<br>average |
| --- | --- | --- | --- | --- | --- | --- |
| *Active elastance* |  |  |  |  |  |  |
| $E_a^{LA}$ | 0.336 | **0.374** | 0.334 | 0.316 | 0.351 | 0.342 |
| $E_a^{RA}$ | 0.356 | **0.399** | 0.357 | 0.335 | 0.361 | 0.362 |
| $E_a^{LV}$ | 0.990 | 0.996 | 0.939 | 0.991 | **0.998** | 0.983 |
| $E_a^{RV}$ | 0.985 | 0.995 | 0.965 | 0.983 | **0.996** | 0.985 |
| *Passive elastance* |  |  |  |  |  |  |
| $E_p^{LA}$ | 0.940 | **0.959** | 0.933 | 0.932 | 0.957 | 0.944 |
| $E_p^{RA}$ | 0.938 | **0.948** | 0.932 | 0.935 | 0.945 | 0.939 |
| $E_p^{LV}$ | 0.983 | 0.991 | 0.978 | 0.975 | **0.992** | 0.984 |
| $E_p^{RV}$ | 0.979 | 0.988 | 0.974 | 0.978 | **0.989** | 0.982 |
| *Unstressed volume* |  |  |  |  |  |  |
| $V_0^{LA}$ | 0.774 | **0.797** | 0.772 | 0.754 | 0.784 | 0.776 |
| $V_0^{RA}$ | 0.724 | **0.749** | 0.722 | 0.713 | 0.729 | 0.727 |
| $V_0^{LV}$ | 0.993 | 0.999 | 0.995 | 0.998 | **1.000** | 0.997 |
| $V_0^{RV}$ | 0.785 | **0.875** | 0.744 | 0.744 | 0.863 | 0.802 |
| *Systemic vascular* |  |  |  |  |  |  |
| $R_A^S$ | 0.957 | 0.985 | 0.906 | 0.960 | **0.988** | 0.959 |
| $R_V^S$ | 0.990 | 0.996 | 0.980 | 0.991 | **0.997** | 0.991 |
| $C_A^S$ | 0.996 | 0.999 | 0.989 | 0.997 | **1.000** | 0.996 |
| $C_V^S$ | 0.459 | **0.659** | 0.293 | 0.406 | 0.622 | 0.488 |
| *Pulmonary vascular* |  |  |  |  |  |  |
| $R_A^P$ | 0.899 | **0.940** | 0.899 | 0.873 | 0.937 | 0.910 |
| $R_V^P$ | 0.914 | **0.951** | 0.913 | 0.899 | 0.948 | 0.925 |
| $C_A^P$ | 0.988 | 0.994 | 0.986 | 0.984 | **0.996** | 0.989 |
| $C_V^P$ | 0.705 | **0.826** | 0.685 | 0.621 | 0.785 | 0.724 |
| Macro average $R^2$ | 0.835 | **0.871** | 0.815 | 0.819 | 0.862 | 0.840 |

## B Neural Prediction, Uncertainty, and Measurement Availability

Two separately trained Mada variants are reported in this appendix. The dedicated full-observation Mada model was trained and evaluated using all 30 measurements and is used for the complete-observation comparison with the neural baselines. The mask-aware Mada model was trained across variable measurement-availability patterns and is used for the random-missingness, structured-removal, and fixed-observation experiments. Consequently, the complete-input result of the mask-aware model should not be interpreted as the result of the dedicated full-observation model.

### B.1 Cross-Model Parameter-Wise Performance

The complete parameter-wise comparison across the full-observation neural baselines is reported in Table 8. Parameters with relatively low and high recoverability were broadly consistent across architectures, supporting the interpretation that much of the parameter-wise recovery pattern reflects the observation-to-parameter inverse problem rather than a single model architecture.

### B.3 Mada Parameter-Wise Predictive Uncertainty

Table 9 reports the parameter-wise results of the dedicated full-observation Mada model. The table includes the validation-derived recoverability score *q*_*j*_, held-out test-set *R*^2^, NRMSE, validation-derived residual-scale prior *σ*_prior, *j*_, mean normalized predictive standard deviation, raw Std./RMSE ratio, raw PICP_95_, and sample-level error–uncertainty correlation for all 20 inverse targets.

**Table 9.**
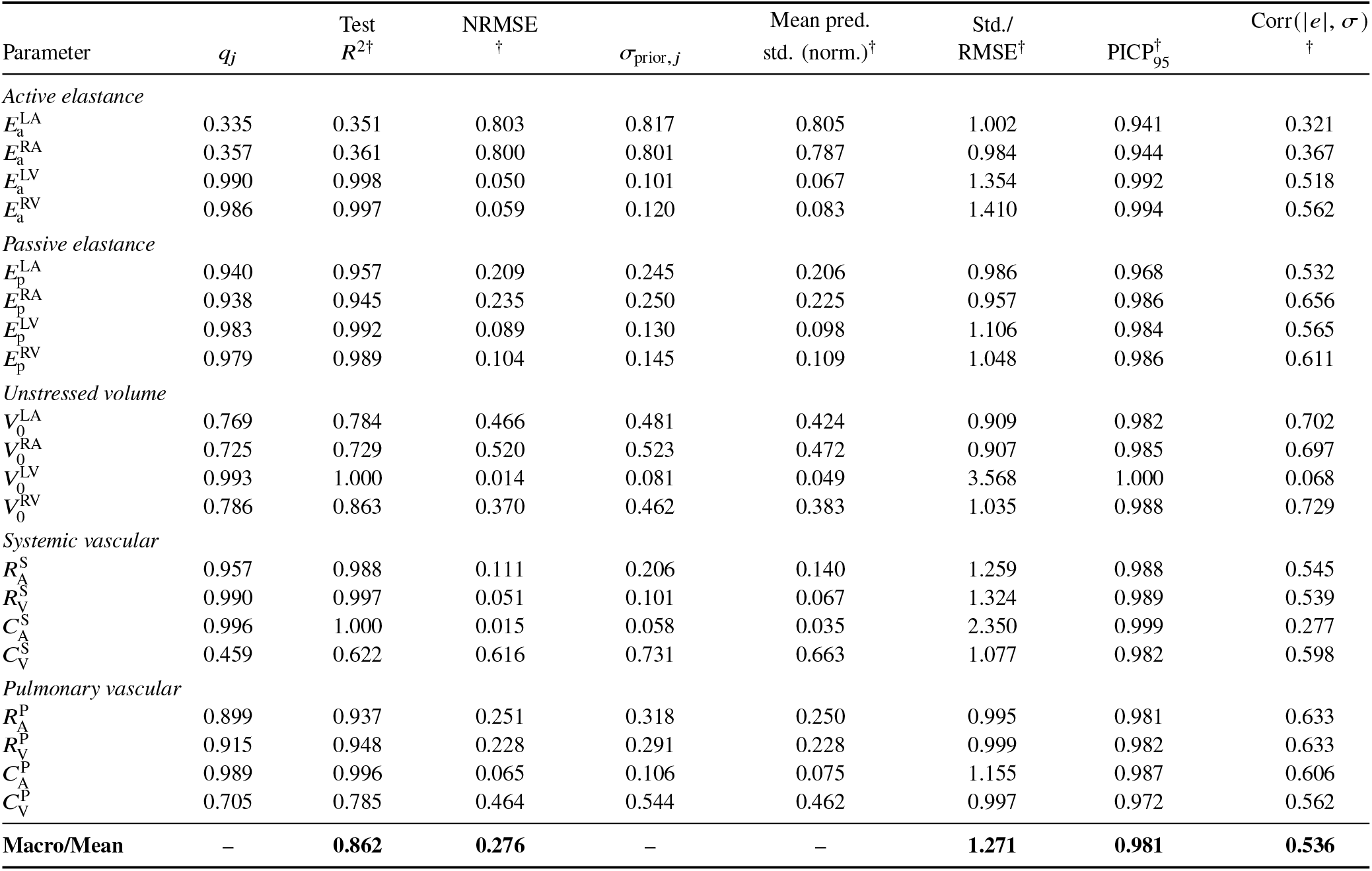
Parameter-wise recoverability and predictive uncertainty of the dedicated full-observation Mada model under complete hemodynamic observations. The recoverability prior and uncertainty prior were derived exclusively from validation predictions. Std./RMSE and PICP_95_ correspond to raw uncertainty before post-hoc calibration. Corr(|*e*|, *σ*) is the Pearson correlation between sample-level absolute prediction error and predicted standard deviation. The dagger identifies metrics computed from held-out test predictions; the recoverability and residual-scale priors are derived from validation data.

The atrial active elastances had low recoverability scores, with *q*_*j*_ = 0.335 for 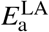 and *q*_*j*_ = 0.357 for 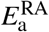, and large mean raw normalized predictive standard deviations of 0.805 and 0.787, respectively. Before calibration, macro Std./RMSE was 1.271 and mean PICP_95_ was 0.981; after parameter-wise validation calibration, they were 1.045 and 0.974, respectively. Mean sample-level error–uncertainty correlation was 0.536. Multiplicative calibration changed uncertainty scale and interval coverage but not predictive means or within-parameter error–uncertainty correlations.

### B.3 Measurement Availability and Clinical Observation Details

All experiments in this subsection used the same frozen mask-aware Mada checkpoint. No missingness-level-specific, measurement-group-specific, or clinical-scenario-specific retraining or fine-tuning was performed. Predictive uncertainty used the validation-only calibration procedures in Sec. 2.3.

#### B.3.1 Parameter-Wise Robustness Under Random Measurement Missingness

Table 10 reports parameter-wise test-set *R*^2^ and mean calibrated normalized predictive standard deviation at missingness levels of 0%, 10%, 30%, 50%, and 70%.

**Table 10.** Parameter-wise inverse-prediction performance and predictive uncertainty of mask-aware Mada under increasing random measurement missingness. Each entry reports the test-set *R*^2^, followed in parentheses by the mean calibrated normalized predictive standard deviation.

| Parameter | 0% missing | 10% missing | 30% missing | 50% missing | 70% missing |
| --- | --- | --- | --- | --- | --- |
| *Active elastance* |  |  |  |  |  |
| $E_a^{LA}$ | 0.333 (0.823) | 0.316 (0.832) | 0.259 (0.865) | 0.176 (0.910) | 0.089 (0.959) |
| $E_a^{RA}$ | 0.334 (0.814) | 0.319 (0.824) | 0.266 (0.857) | 0.182 (0.903) | 0.092 (0.951) |
| $E_a^{LV}$ | 0.996 (0.094) | 0.960 (0.155) | 0.764 (0.392) | 0.457 (0.741) | 0.168 (1.085) |
| $E_a^{RV}$ | 0.992 (0.122) | 0.932 (0.201) | 0.803 (0.386) | 0.636 (0.605) | 0.412 (0.869) |
| *Passive elastance* |  |  |  |  |  |
| $E_p^{LA}$ | 0.947 (0.242) | 0.925 (0.273) | 0.844 (0.377) | 0.688 (0.545) | 0.452 (0.782) |
| $E_p^{RA}$ | 0.940 (0.250) | 0.923 (0.281) | 0.844 (0.386) | 0.671 (0.564) | 0.388 (0.808) |
| $E_p^{LV}$ | 0.989 (0.112) | 0.979 (0.143) | 0.932 (0.247) | 0.832 (0.414) | 0.623 (0.636) |
| $E_p^{RV}$ | 0.984 (0.134) | 0.961 (0.190) | 0.876 (0.344) | 0.693 (0.556) | 0.415 (0.793) |
| *Unstressed volume* |  |  |  |  |  |
| $V_0^{LA}$ | 0.769 (0.487) | 0.748 (0.504) | 0.642 (0.589) | 0.477 (0.719) | 0.281 (0.860) |
| $V_0^{RA}$ | 0.712 (0.541) | 0.694 (0.553) | 0.613 (0.618) | 0.464 (0.725) | 0.281 (0.847) |
| $V_0^{LV}$ | 1.000 (0.027) | 0.998 (0.043) | 0.966 (0.128) | 0.861 (0.361) | 0.624 (0.883) |
| $V_0^{RV}$ | 0.818 (0.432) | 0.713 (0.527) | 0.530 (0.685) | 0.355 (0.806) | 0.199 (0.894) |
| *Systemic vascular* |  |  |  |  |  |
| $R_A^S$ | 0.982 (0.154) | 0.947 (0.213) | 0.879 (0.325) | 0.764 (0.479) | 0.502 (0.751) |
| $R_V^S$ | 0.996 (0.072) | 0.989 (0.095) | 0.977 (0.142) | 0.960 (0.204) | 0.909 (0.320) |
| $C_A^S$ | 1.000 (0.041) | 0.986 (0.072) | 0.867 (0.237) | 0.607 (0.611) | 0.280 (1.253) |
| $C_V^S$ | 0.526 (0.681) | 0.396 (0.774) | 0.230 (0.877) | 0.144 (0.921) | 0.082 (0.948) |
| *Pulmonary vascular* |  |  |  |  |  |
| $R_A^P$ | 0.914 (0.311) | 0.838 (0.396) | 0.670 (0.557) | 0.481 (0.715) | 0.274 (0.876) |
| $R_V^P$ | 0.927 (0.283) | 0.867 (0.357) | 0.733 (0.501) | 0.579 (0.643) | 0.380 (0.809) |
| $C_A^P$ | 0.993 (0.106) | 0.966 (0.170) | 0.824 (0.357) | 0.559 (0.643) | 0.250 (1.000) |
| $C_V^P$ | 0.698 (0.560) | 0.544 (0.669) | 0.287 (0.838) | 0.119 (0.939) | 0.035 (0.991) |
| **Macro/Mean** | **0.842 (0.314)** | **0.800 (0.364)** | **0.690 (0.485)** | **0.535 (0.650)** | **0.337 (0.866)** |

The evaluation used the fixed data partition and test-mask seeds.

For 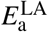, *R*^2^ decreased from 0.333 under complete observations to 0.089 at 70% missingness, while mean normalized calibrated predictive standard deviation increased from 0.823 to 0.959. Parameter-wise differences persisted, demonstrating that missingness fraction alone does not determine prediction reliability.

#### B.3.2 Detailed Physiological Measurement–Parameter Dependencies

For removed physiological measurement group *g* and inverse target *j*, Table 11 reports the recoverability loss 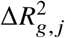 and calibrated predictive uncertainty-inflation factor 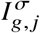 relative to complete observations for all 140 measurement-group–parameter combinations.

**Table 11.**
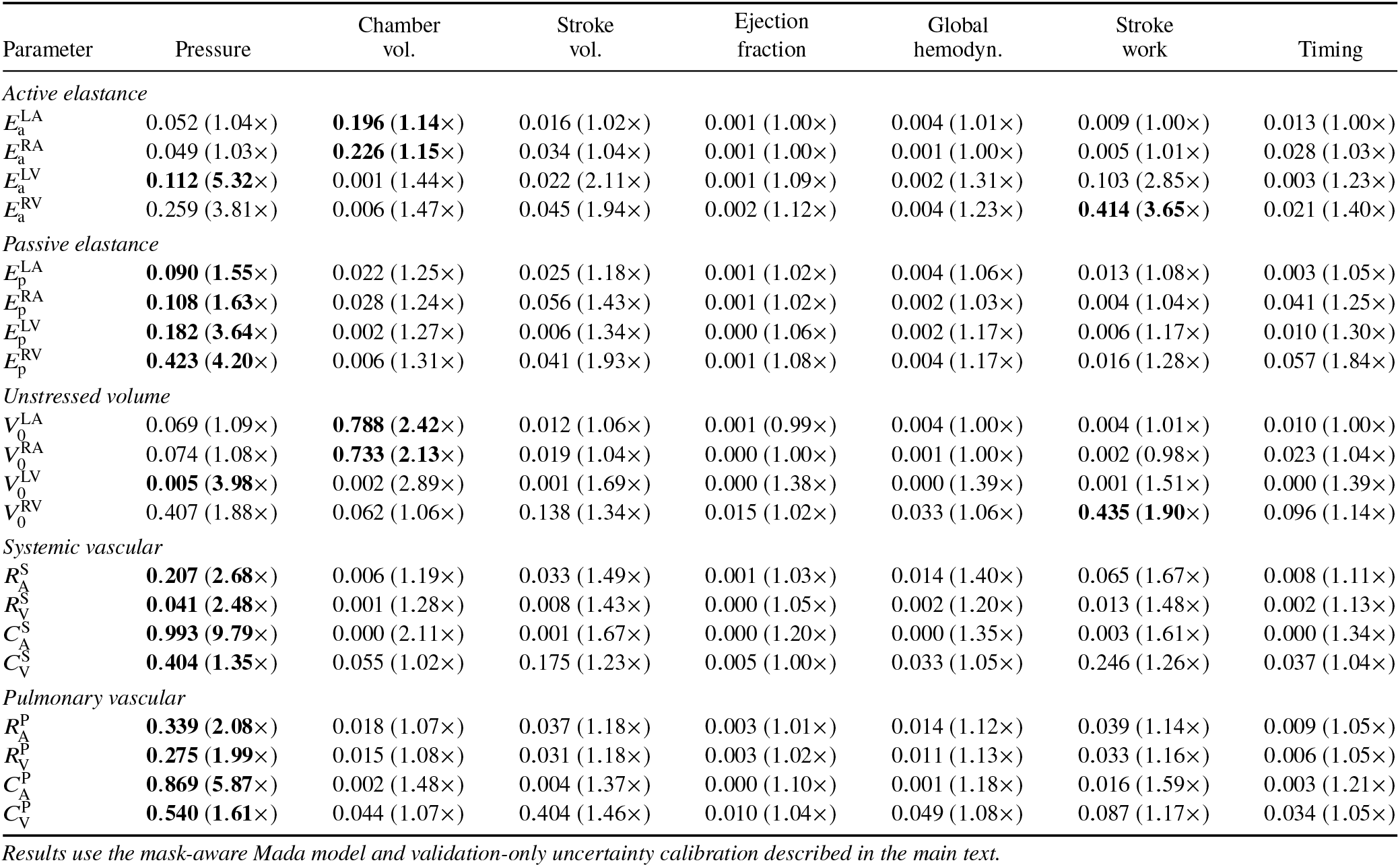
Detailed physiological measurement–parameter dependencies obtained from structured measurement-group masking. For removed measurement group *g* and inverse target *j*, each entry reports recoverability loss 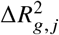, followed in parentheses by calibrated predictive uncertainty inflation 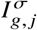.

Pressure removal produced the largest macro-averaged recoverability loss (0.275), followed by chamber-volume removal (0.111) and stroke-work removal (0.076). Pressure and chamber-volume measurements were especially important for vascular parameters and unstressed chamber volumes, respectively. Dependencies also differed within the active-elastance group: pressure removal reduced 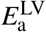 recovery by 0.112, whereas stroke-work removal reduced 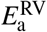 recovery by 0.414. Uncertainty inflation broadly accompanied these losses (Table 11).

#### B.3.3 Uncertainty Calibration Under Structured Measurement Loss

Table 12 summarizes uncertainty calibration under structured measurement removal. The Std./RMSE ratio remained broadly close to one. Chamber-volume removal produced the most conservative average uncertainty scale; global-hemodynamics, ejection-fraction, timing, and stroke-volume removal also produced ratios above one. PICP_95_ remained close to nominal in most conditions, with pressure removal providing the principal undercoverage case.

**Table 12.**
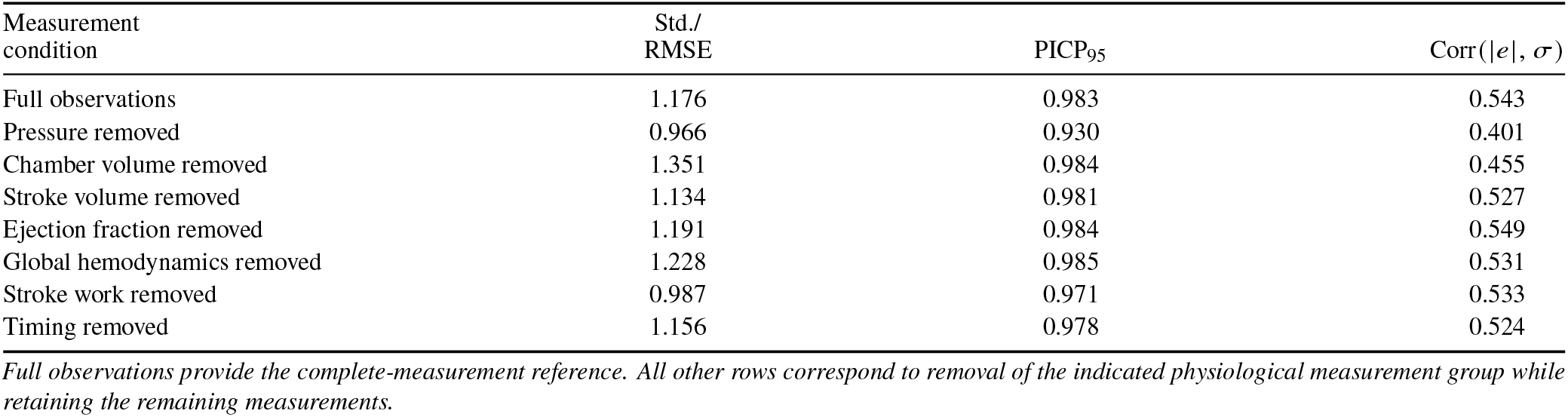
Validation-calibrated uncertainty of mask-aware Mada under structured physiological measurement loss. Metrics are averaged over the 20 targets. Std./RMSE is the parameter-wise mean predicted standard deviation divided by empirical RMSE, with an ideal value of 1; PICP_95_ is the empirical coverage of the nominal 95% prediction intervals, with a nominal value of 0.95; and Corr(|*e*|, *σ*) is the Pearson correlation between sample-level absolute prediction error |*e*| and predicted standard deviation *σ*, where *e* denotes the normalized prediction error.

| Measurement condition | Std./RMSE | PICP<sub>95</sub> | Corr($\vert e\vert $, $\sigma$) |
| --- | --- | --- | --- |
| Full observations | 1.176 | 0.983 | 0.543 |
| Pressure removed | 0.966 | 0.930 | 0.401 |
| Chamber volume removed | 1.351 | 0.984 | 0.455 |
| Stroke volume removed | 1.134 | 0.981 | 0.527 |
| Ejection fraction removed | 1.191 | 0.984 | 0.549 |
| Global hemodynamics removed | 1.228 | 0.985 | 0.531 |
| Stroke work removed | 0.987 | 0.971 | 0.533 |
| Timing removed | 1.156 | 0.978 | 0.524 |

Full observations provide the complete-measurement reference. All other rows correspond to removal of the indicated physiological measurement group while retaining the remaining measurements.

**Table 13.**
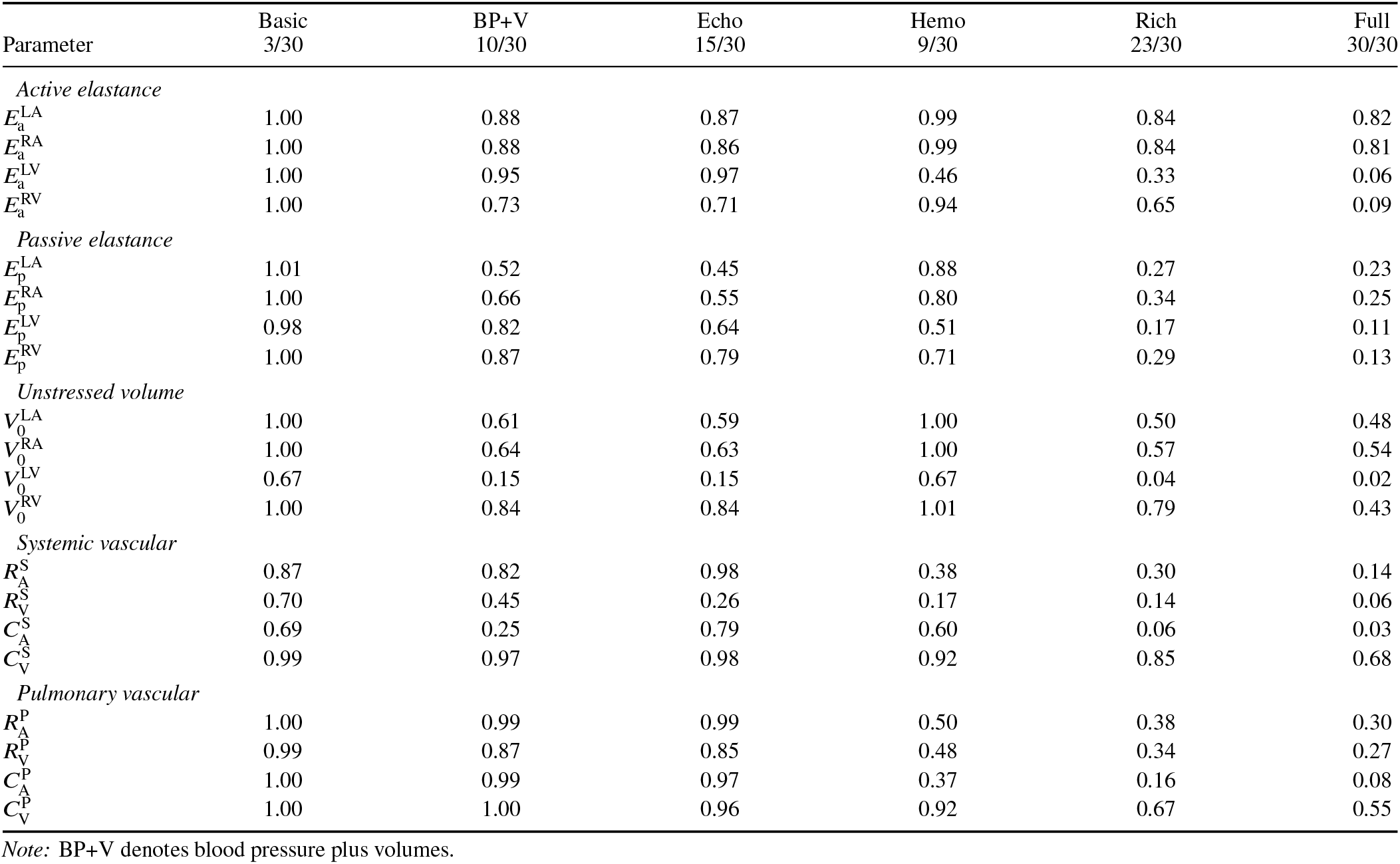
Calibrated parameter-wise predictive uncertainty for the six available-observation sets. Entries are the mean predictive standard deviation normalized by the corresponding training-target standard deviation.

| Parameter | Basic<br>3/30 | BP+V<br>10/30 | Echo<br>15/30 | Hemo<br>9/30 | Rich<br>23/30 | Full<br>30/30 |
| --- | --- | --- | --- | --- | --- | --- |
| *Active elastance* |  |  |  |  |  |  |
| $E_a^{LA}$ | 1.00 | 0.88 | 0.87 | 0.99 | 0.84 | 0.82 |
| $E_a^{RA}$ | 1.00 | 0.88 | 0.86 | 0.99 | 0.84 | 0.81 |
| $E_a^{LV}$ | 1.00 | 0.95 | 0.97 | 0.46 | 0.33 | 0.06 |
| $E_a^{RV}$ | 1.00 | 0.73 | 0.71 | 0.94 | 0.65 | 0.09 |
| *Passive elastance* |  |  |  |  |  |  |
| $E_p^{LA}$ | 1.01 | 0.52 | 0.45 | 0.88 | 0.27 | 0.23 |
| $E_p^{RA}$ | 1.00 | 0.66 | 0.55 | 0.80 | 0.34 | 0.25 |
| $E_p^{LV}$ | 0.98 | 0.82 | 0.64 | 0.51 | 0.17 | 0.11 |
| $E_p^{RV}$ | 1.00 | 0.87 | 0.79 | 0.71 | 0.29 | 0.13 |
| *Unstressed volume* |  |  |  |  |  |  |
| $V_0^{LA}$ | 1.00 | 0.61 | 0.59 | 1.00 | 0.50 | 0.48 |
| $V_0^{RA}$ | 1.00 | 0.64 | 0.63 | 1.00 | 0.57 | 0.54 |
| $V_0^{LV}$ | 0.67 | 0.15 | 0.15 | 0.67 | 0.04 | 0.02 |
| $V_0^{RV}$ | 1.00 | 0.84 | 0.84 | 1.01 | 0.79 | 0.43 |
| *Systemic vascular* |  |  |  |  |  |  |
| $R_A^S$ | 0.87 | 0.82 | 0.98 | 0.38 | 0.30 | 0.14 |
| $R_V^S$ | 0.70 | 0.45 | 0.26 | 0.17 | 0.14 | 0.06 |
| $C_A^S$ | 0.69 | 0.25 | 0.79 | 0.60 | 0.06 | 0.03 |
| $C_V^S$ | 0.99 | 0.97 | 0.98 | 0.92 | 0.85 | 0.68 |
| *Pulmonary vascular* |  |  |  |  |  |  |
| $R_A^P$ | 1.00 | 0.99 | 0.99 | 0.50 | 0.38 | 0.30 |
| $R_V^P$ | 0.99 | 0.87 | 0.85 | 0.48 | 0.34 | 0.27 |
| $C_A^P$ | 1.00 | 0.99 | 0.97 | 0.37 | 0.16 | 0.08 |
| $C_V^P$ | 1.00 | 1.00 | 0.96 | 0.92 | 0.67 | 0.55 |

Note: BP+V denotes blood pressure plus volumes.

**Table 14.** Parameter-wise calibration of the 95% prediction intervals for the six available-observation sets. Entries report validation-calibrated PICP_95_; the nominal target is 0.95.

| Parameter | Basic<br>3/30 | BP+V<br>10/30 | Echo<br>15/30 | Hemo<br>9/30 | Rich<br>23/30 | Full<br>30/30 |
| --- | --- | --- | --- | --- | --- | --- |
| *Active elastance* |  |  |  |  |  |  |
| $E_a^{LA}$ | 0.94 | 0.94 | 0.94 | 0.93 | 0.94 | 0.94 |
| $E_a^{RA}$ | 0.94 | 0.94 | 0.94 | 0.94 | 0.95 | 0.95 |
| $E_a^{LV}$ | 0.94 | 0.94 | 0.94 | 0.97 | 0.99 | 0.96 |
| $E_a^{RV}$ | 0.93 | 0.97 | 0.97 | 0.93 | 0.98 | 0.97 |
| *Passive elastance* |  |  |  |  |  |  |
| $E_p^{LA}$ | 0.93 | 0.96 | 0.97 | 0.95 | 0.98 | 0.97 |
| $E_p^{RA}$ | 0.94 | 0.96 | 0.96 | 0.99 | 0.99 | 0.99 |
| $E_p^{LV}$ | 0.94 | 0.94 | 0.95 | 0.96 | 0.97 | 0.97 |
| $E_p^{RV}$ | 0.93 | 0.94 | 0.94 | 0.95 | 0.98 | 0.98 |
| *Unstressed volume* |  |  |  |  |  |  |
| $V_0^{LA}$ | 0.94 | 1.00 | 0.99 | 0.94 | 1.00 | 1.00 |
| $V_0^{RA}$ | 0.95 | 0.99 | 0.99 | 0.95 | 0.99 | 1.00 |
| $V_0^{LV}$ | 0.85 | 0.96 | 0.96 | 0.85 | 0.94 | 0.97 |
| $V_0^{RV}$ | 0.99 | 1.00 | 0.99 | 0.98 | 0.99 | 1.00 |
| *Systemic vascular* |  |  |  |  |  |  |
| $R_A^S$ | 0.96 | 0.96 | 0.98 | 0.96 | 0.98 | 0.98 |
| $R_V^S$ | 0.96 | 0.97 | 0.96 | 0.96 | 0.97 | 0.96 |
| $C_A^S$ | 0.95 | 0.96 | 0.91 | 0.95 | 0.95 | 0.97 |
| $C_V^S$ | 0.93 | 0.93 | 0.94 | 0.94 | 0.95 | 0.97 |
| *Pulmonary vascular* |  |  |  |  |  |  |
| $R_A^P$ | 0.94 | 0.94 | 0.94 | 0.98 | 0.99 | 0.99 |
| $R_V^P$ | 0.94 | 0.97 | 0.97 | 0.98 | 0.98 | 0.99 |
| $C_A^P$ | 0.95 | 0.95 | 0.96 | 0.96 | 0.98 | 0.97 |
| $C_V^P$ | 0.95 | 0.95 | 0.96 | 0.95 | 0.97 | 0.97 |

Note: BP+V denotes blood pressure plus volumes.

Error–uncertainty correlation was more sensitive to the type of missing information and decreased substantially after pressure removal.

#### B.3.4 Detailed Fixed-Observation Scenario Protocol

The six clinically motivated observation sets were defined as follows:

- *Basic clinical*: systemic systolic and diastolic arterial pressures and heart rate (3 of 30 measurements);
- *BP + chamber volumes*: systemic systolic and diastolic arterial pressures together with the maximum and minimum volumes of the four cardiac chambers (10 of 30);
- *Echo-like*: maximum and minimum chamber volumes, left- and right-ventricular ejection fractions, chamber stroke volumes, and heart rate (15 of 30);
- *Hemodynamic*: the eight pressure measurements and cardiac output (9 of 30);
- *Rich clinical*: pressures, maximum and minimum chamber volumes, cardiac output, chamber stroke volumes, and ventricular ejection fractions (23 of 30); and
- *Full synthetic*: all 30 measurements.

Let *s* = 1, …, 6 index the six fixed observation scenarios. For each scenario *s*, a binary availability mask **m**(*s*) *∈* {0, 1}^30^ was applied to every validation and test sample. An entry of one indicated an available measurement, whereas zero indicated an unavailable measurement. The mask itself was concatenated to the masked normalized input, so unavailable measurements were explicitly identified rather than imputed.

Because the fixed clinical masks differ from the random masking patterns used during training, predictive uncertainty was recalibrated separately for each scenario using validation data only. The validation scale ratio defined in Eq. (24) was clipped to [0.4, 2.5] independently for each fixed scenario and inverse target. For scenario *s* and inverse target *j*,

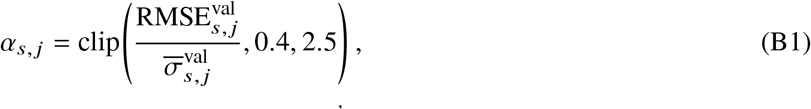

where 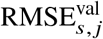 denotes the validation root-mean-square error and 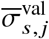 denotes the mean uncalibrated predictive standard deviation for target *j* under scenario *s*. No test-set-dependent adjustment was applied. With all 30 measurements available, the random-missingness and fixed-observation protocols use identical predictive means and raw standard deviations but different calibration factors; their calibrated full-input uncertainty is therefore protocol specific.

For sample *n*, the calibrated predictive standard deviation was then

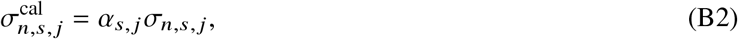

where *n* indexes samples, *σ*_*n,s, j*_ is the uncalibrated predictive standard deviation, and 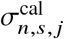 is the corresponding calibrated value. Parameter recovery was reported using *R*^2^, NRMSE, and PCC; uncertainty was summarized using mean normalized standard deviation, Std./RMSE, PICP_95_, and cross-parameter correspondence between raw uncertainty and realized NRMSE.

#### B.3.5 Detailed Parameter-Wise Recovery

Table 15 reports parameter-wise *R*^2^ values across the six fixed observation sets.

**Table 15.** Detailed parameter-wise recovery across six fixed, clinically motivated observation sets. Each entry reports the held-out test-set coefficient of determination, *R*^2^, for one of the 20 inverse LPN targets. All scenarios used the same frozen mask-aware Mada checkpoint.

| *(a) Chamber-mechanics parameters* |  |  |  |  |  |  |  |  |  |  |  |  |
| --- | --- | --- | --- | --- | --- | --- | --- | --- | --- | --- | --- | --- |
| Observation set | Active elastance |  |  |  | Passive elastance |  |  |  | Unstressed volume |  |  |  |
| | $E_a^{LA}$ | $E_a^{RA}$ | $E_a^{LV}$ | $E_a^{RV}$ | $E_p^{LA}$ | $E_p^{RA}$ | $E_p^{LV}$ | $E_p^{RV}$ | $V_0^{LA}$ | $V_0^{RA}$ | $V_0^{LV}$ | $V_0^{RV}$ |
| Basic (3/30) | 0.00 | 0.00 | 0.00 | 0.00 | −0.02 | 0.00 | 0.04 | 0.00 | 0.00 | 0.00 | 0.00 | 0.00 |
| BP + chamber volumes (10/30) | 0.23 | 0.22 | 0.10 | 0.47 | 0.73 | 0.56 | 0.33 | 0.25 | 0.63 | 0.58 | 0.98 | 0.29 |
| Echo-like (15/30) | 0.24 | 0.26 | 0.07 | 0.50 | 0.80 | 0.69 | 0.59 | 0.38 | 0.65 | 0.60 | 0.98 | 0.30 |
| Hemodynamic (9/30) | 0.02 | 0.02 | 0.79 | 0.13 | 0.23 | 0.37 | 0.73 | 0.50 | 0.00 | −0.01 | 0.00 | −0.03 |
| Rich clinical (23/30) | 0.31 | 0.30 | 0.89 | 0.58 | 0.93 | 0.88 | 0.97 | 0.91 | 0.75 | 0.68 | 1.00 | 0.38 |
| Full synthetic (30/30) | 0.33 | 0.33 | 1.00 | 0.99 | 0.95 | 0.94 | 0.99 | 0.98 | 0.77 | 0.71 | 1.00 | 0.82 |

| *(b) Vascular parameters* |  |  |  |  |  |  |  |  |  |
| --- | --- | --- | --- | --- | --- | --- | --- | --- | --- |
| Observation set | Systemic vascular |  |  |  | Pulmonary vascular |  |  |  |  |
| | $R_A^S$ | $R_V^S$ | $C_A^S$ | $C_V^S$ | $R_A^P$ | $R_V^P$ | $C_A^P$ | $C_V^P$ | |
| Basic (3/30) | 0.24 | 0.52 | 0.52 | 0.00 | 0.00 | 0.02 | 0.00 | 0.00 |  |
| BP + chamber volumes (10/30) | 0.33 | 0.80 | 0.94 | 0.06 | 0.02 | 0.25 | 0.00 | 0.01 |  |
| Echo-like (15/30) | 0.04 | 0.93 | 0.00 | 0.03 | 0.01 | 0.29 | 0.05 | 0.08 |  |
| Hemodynamic (9/30) | 0.85 | 0.97 | 0.63 | 0.14 | 0.76 | 0.77 | 0.86 | 0.15 |  |
| Rich clinical (23/30) | 0.91 | 0.98 | 1.00 | 0.27 | 0.86 | 0.88 | 0.98 | 0.56 |  |
| Full synthetic (30/30) | 0.98 | 1.00 | 1.00 | 0.53 | 0.91 | 0.93 | 0.99 | 0.70 |  |

#### B.3.6 Error–Uncertainty Correspondence Under Fixed Clinical Observation Sets

To complement interval coverage, the realized test-set NRMSE of each parameter was compared with its mean raw predictive standard deviation under each fixed observation scenario. Both quantities were expressed in normalized target units and plotted on logarithmic axes.

The cross-parameter log-space Pearson correlations, *r*_log_, were 0.53 for the basic clinical set, 0.89 for BP + chamber volumes, 0.80 for the Echo-like set, 0.77 for the hemodynamic set, and 0.98 for both the rich clinical and full synthetic sets. These associations concern raw uncertainty across parameters and complement, rather than replace, interval coverage and sample-level error–uncertainty correlation.

## C Recoverability and Practical-Ambiguity Diagnostics

Direct optimization, waveform replays, and compensation profiles use the dataset-generation LPN forward solver. The diagnostic cohort comprises 20 fixed held-out cases, with Cases 15, 17, 08, 06, and 07 examined in detail. The target-specific neural analysis itself includes the separately trained multi-output reference and the two fine-tuned single-output ResMLP models described below.

This appendix provides the detailed protocols and results for the direct optimization benchmark, the single-output atrial active-elastance diagnostic, and the profile-based compensation analysis. The complementary local forward-Jacobian sensitivity analysis is reported separately in Appendix D.6.

### C.1 Direct LPN Optimization Benchmark

The unknown vector contains 20 parameters. The 27 hemodynamic outputs define the residual in Eq. (30); BPM, PR interval, and EMD are fixed case-specific timing inputs. Parameter bounds and range-normalized error denominators are the original generating bounds. Output residuals use the training-set standard deviations.

Two fixed numerical starts, A and B, are used for every case and optimizer. Start A is the midpoint of the training-set parameter-extrema intervals, and start B is a seed-20 uniform draw from the same intervals. Both starts lie within the generating bounds used as optimization constraints. The 20-case cohort was selected by a maximin design in the training-standardized coordinates of LA/RA active elastance, LA/RA/RV unstressed volume, and systemic/pulmonary venous compliance. Its space-filling *R*^2^ values are descriptive cohort statistics, not population estimates.

The trained full-observation Mada model reported in Table 2 was also evaluated on the 20-case diagnostic cohort. Its parameter estimates were projected to the generating bounds for the benchmark parameter-error and cross-case *R*^2^ calculations and for forward reconstruction using the original LPN solver. TRF and Dogbox use SciPy 1.11.1 bounded least squares with a two-point numerical Jacobian, Jacobian-based parameter scaling, and max_nfev= 2000. The tolerances are

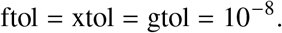

L-BFGS-B minimizes the mean squared standardized residual with numerical gradients, maxiter= 2000, maxfun= 100000, and ftol = gtol = 10−8. Least-squares nfev counts reported residual evaluations, not iterations, and excludes additional finite-difference residual calls. Total forward calls are recorded separately.

The forward solver uses the dataset-generation configuration, initial state, activation timing, integration scheme, and clinical-output extraction. Integration starts with 10 cardiac cycles and 1,000 steps per cycle, so Δ*t* = *T*/1000 for cardiac period *T*. If consecutive-cycle volumes and vascular pressures do not repeat within the original relative/absolute tolerances 10−3 and 0.1, integration continues one cycle at a time up to 50 cycles. The last two cycles define the summary outputs; the final cycle supplies waveform comparisons.

Forward-solver failures were treated as unsuccessful optimization attempts without introducing artificial residuals. For each case and optimizer, the solution with the lower forward NRMSE was selected independently of parameter-recovery accuracy.

### C.2 Detailed Optimization Results

Ground-truth replay converged for all 20 cases. Its maximum standardized forward NRMSE was 1.67 × 10−16 and the largest absolute output difference was 3.64 × 10−12. Forward residuals therefore measure candidate-solution disagreement with the same data-generation model.

Mean cross-case *R*^2^ over the seven emphasized parameters was 0.601 for Mada, 0.588 for selected TRF, 0.194 for Dogbox, and −0.001 for L-BFGS-B. Selected TRF LA/RA *R*^2^ values were 0.236/0.352, versus 0.139/0.441 for Mada. Across the complete 20-case diagnostic cohort, the median forward NRMSE was 0.002 for selected TRF and 0.054 for Mada. The corresponding median mean range-normalized parameter errors were 0.069 and 0.075, respectively. Thus, selected TRF achieved substantially closer hemodynamic reconstruction while providing broadly comparable parameter recovery on this finite diagnostic cohort.

Median range-normalized LA/RA errors were 0.172/0.183 for Mada and 0.187/0.118 for TRF. Selected TRF median errors for RV unstressed volume, systemic venous compliance, and pulmonary venous compliance were 0.023, 0.172, and 0.082.

Within the five cases retained for detailed visualization, rather than the complete 20-case cohort, the median forward NRMSE was 0.100 for Mada and 0.002 for selected TRF; the corresponding median mean range-normalized parameter errors were 0.098 and 0.144. Selected TRF forward NRMSE for Cases 15, 17, 08, 06, and 07 was 1.07 × 10−2, 1.19 × 10−4, 3.75 × 10−3, 1.86 × 10−3, and 5.43 × 10−7, respectively. Across all 20 cases, 1 Mada, 19 TRF-A, and 20 TRF-B fits met NRMSE ≤ 0.02.

Optimizer convergence was 40/40 for TRF, 35/40 for Dogbox, and 31/40 for L-BFGS-B. Five Dogbox runs exhausted their evaluation budget; nine L-BFGS-B runs ended in abnormal line-search termination. Final LPN replays converged periodically for all 120 direct-optimizer runs. Per-run runtime, reported evaluations, and actual forward-call counts are listed in Table 22; the two starts must both be counted when assessing case-wise inversion cost.

### C.3 Detailed LPN Forward Reconstruction

The same original solver and case-specific timing values generated trajectories at the recorded parameters, the Mada estimate, and both optimized TRF vectors. GT traces are forward replays at generating parameters; the stored dataset contains summary observations rather than measured waveforms.

The waveform comparisons retain differences not fully represented by a scalar summary residual. Matching the 27 summary observations does not require agreement at every time point.

### C.4 Single-Output Atrial Active-Elastance Diagnostic

A 20-output ResMLP was trained using seed 20 to provide a reproducible multi-output reference and predictions for the same 20 cases used in the LPN diagnostic benchmark. The model used a 30-dimensional input layer, hidden dimension 256, four fully connected residual blocks, Gaussian error linear unit (GELU) activation, layer normalization (LayerNorm), dropout 0.1, and a 20-dimensional linear output layer (544,020 trainable parameters). Two single-output ResMLPs were then constructed for 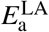 and 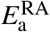. Each retained the same input and four-block backbone but used a one-dimensional output head (539,137 trainable parameters). To reduce diagnostic cost, the single-output models were initialized from the multi-output ResMLP, including the corresponding row of the output head, and then fine-tuned end-to-end using a target-specific mean-squared-error (MSE) loss. For inverse target *j*, let *N*_train_ denote the number of training cases, 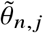 the normalized ground-truth target for training case *n*, and 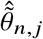 the corresponding normalized single-output prediction. The loss was

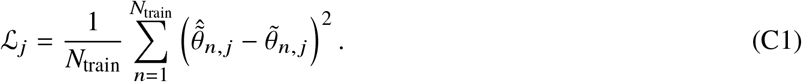

The multi-output model used initial learning rate 10−3, up to 50 epochs, and patience 8. The single-output models used initial learning rate 5 × 10−4, up to 25 epochs, patience 6, learning-rate reduction by 0.5 after validation stagnation, gradient clipping at 5, and batch size 4096. Only seed 20 was used because this was a mechanistic diagnostic rather than a multi-seed architecture comparison.

The diagnostic multi-output ResMLP achieved full-test *R*^2^ = 0.337 for 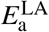 and 0.365 for 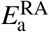. Single-output fine-tuning increased these to 0.339 and 0.372, corresponding to

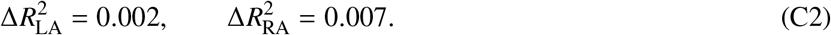

NRMSE changed from 0.812 to 0.811 for LA and from 0.797 to 0.793 for RA. On the 20 optimization cases, LA *R*^2^ changed from 0.166 to 0.152, while RA *R*^2^ changed from 0.430 to 0.487. Thus, improvements were small on the full test set and not uniform on the diagnostic cohort. The 50-epoch multi-output ResMLP used for this diagnostic is distinct from the ResMLP baseline in Table 2.

### C.5 Profile-Based Compensation Diagnostic

Case 07 was used to examine parameter compensation despite the low parameter-recovery error of its selected TRF solution. Let 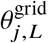 and 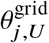 denote the fixed profile-grid endpoints. The profile coordinate is

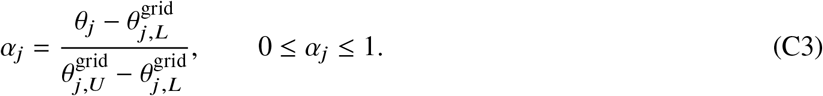

The physical profile-grid endpoints are defined by training-set extrema, while the free-parameter optimization constraints use the generating bounds. At each point, the fixed coordinate is removed from the compensation set

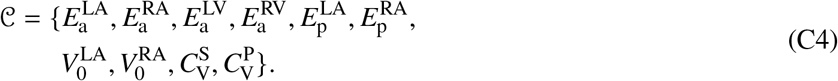

All parameters outside this set remain at generating values. The compensation objective is

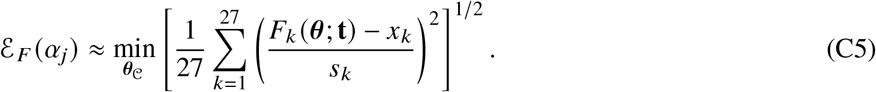

The reported value is the smaller residual found from generating-parameter and selected-TRF warm starts, not a certified global minimum. Seven points each are evaluated for LA, RA, and LV active elastance, plus a 5 × 5 joint LA–RA grid. Free parameters obey the generating bounds and use the same TRF tolerances, 2,000-evaluation limit, and periodic-convergence logic as the benchmark.

A converged profile fit qualifies as a low-error reconstruction when

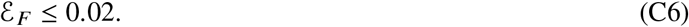

LA qualified at 1/7 positions, with residuals 0.002–0.319; RA qualified at 6/7 positions, with residuals 0.002–0.037; and the LV control qualified at 4/7 positions, with residuals 0.001–0.180. The joint grid contained 4/25 qualifying points and had minimum residual 0.003. All 46 selected profile searches reported optimizer convergence. These values describe the evaluated grid and targeted compensation set.

### C.6 Profile-Solution Waveforms and Compensatory Parameter Paths

#### C.6.1 Pressure and Volume Waveform Envelopes

The four converged low-error joint-profile parameter sets were replayed with the original LPN forward solver. Pointwise percentiles summarize variation within this finite solution set.

#### C.6.3 Compensatory Parameter Trajectories

For compensating parameter ℓ, displacement is normalized by the generating range:

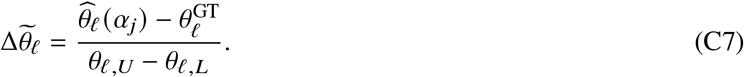

The displayed parameter changes must be read alongside the forward residuals: only 1/7 LA and 6/7 RA positions satisfy the stated tolerance. A large displacement at a high-residual point does not establish effective compensation.

## D Robustness and Model Diagnostics

This appendix provides the detailed protocols and results for five related analyses: modality-dependent measurement noise, the five-case noisy direct-optimization diagnostic, comparison with a simple probabilistic ResMLP, the recoverability-weighted reference-regularization sweep, and local forward-sensitivity analysis. The population-level noise experiments and the five-case forward-replay experiment use the dedicated full-observation Mada model reported in Table 2.

### D.1 Measurement-Noise Protocol

Let *i* index a hemodynamic observation to which measurement noise is applied, and let *x*_*i*_ denote its value in physical units. For each perturbed observation, *ϵ*_*i*_ *∼ N*(0, 1) denotes a standard-normal random draw. Let 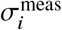 denote the additive measurement-noise standard deviation and let *c*_*i*_ denote the dimensionless multiplicative noise amplitude. Additive perturbations were defined as

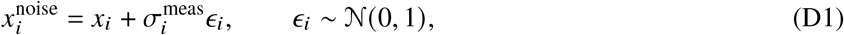

whereas multiplicative perturbations were defined as

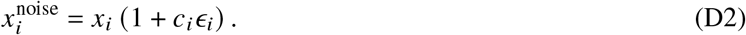

Systemic systolic pressure, systemic diastolic pressure, and mean arterial pressure used an additive measurement-noise standard deviation of 5 mmHg. Pulmonary systolic and diastolic pressure, mean pulmonary arterial pressure, central venous pressure, and pulmonary arterial wedge pressure used an additive standard deviation of 2 mmHg. Maximum and minimum chamber volumes used 5% multiplicative variation, with physiological ordering of extrema enforced after perturbation. Heart rate used an additive standard deviation of 2 beats/min, whereas the PR interval and electromechanical delay used 0.005 s. Stroke volumes and ejection fractions were recomputed from the perturbed volume extrema. Cardiac output was recomputed from the perturbed LV stroke volume and heart rate and then received an additional 7.5% multiplicative flow perturbation. Systemic and pulmonary vascular resistances were recomputed from the corresponding perturbed pressures and cardiac output. LV and RV stroke work received 7.5% multiplicative perturbation. A single seed-20 noise realization was used for reproducibility.

The frozen full-observation Mada checkpoint was evaluated on 65,000 held-out cases under clean and noisy observations. No noisy-data training, fine-tuning, or test-time recalibration was performed; clean validation-derived calibration factors were retained. Predicted means were projected to the training-set parameter extrema before population-level neural scoring:

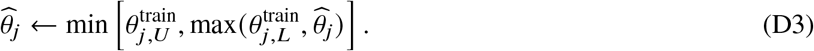

The five-case physics diagnostic uses the seed-20 noisy observations, including perturbed timing and derived outputs, together with predictions from the dedicated full-observation Mada model. Noisy neural estimates are projected to the training-set parameter extrema before forward replay, as in the population-level noise analysis. Bounded TRF optimization uses the two fixed starts and the generating parameter bounds. Let 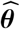 denote the parameter vector recovered from the noisy observations, and let **t**noise denote the corresponding perturbed timing vector. For the original LPN forward-solver outputs, *k* = 1, …, 27 indexes the modeled hemodynamic quantities, *F*_*k*_ denotes modeled output *k*, and *s*_*k*_ denotes its training-set standard deviation.

The noisy-data fitting error measures agreement with the perturbed observations:

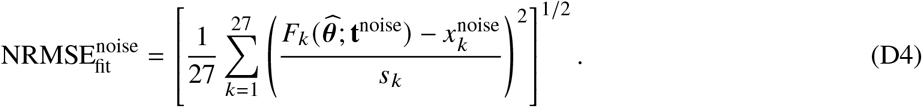

In contrast, the clean-reference forward error measures agreement of the same recovered parameter vector with the original unperturbed observations:

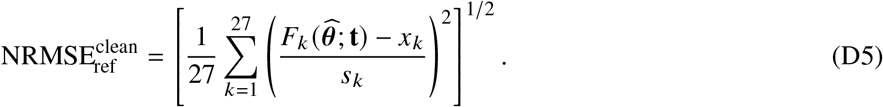

For each case, the TRF run with the lower noisy-data fitting error was selected; ground-truth parameter error was not used in this selection.

Table 16 reports the parameter-wise performance under clean and perturbed observations, complementing the aggregate results presented in the main manuscript.

**Table 16.**
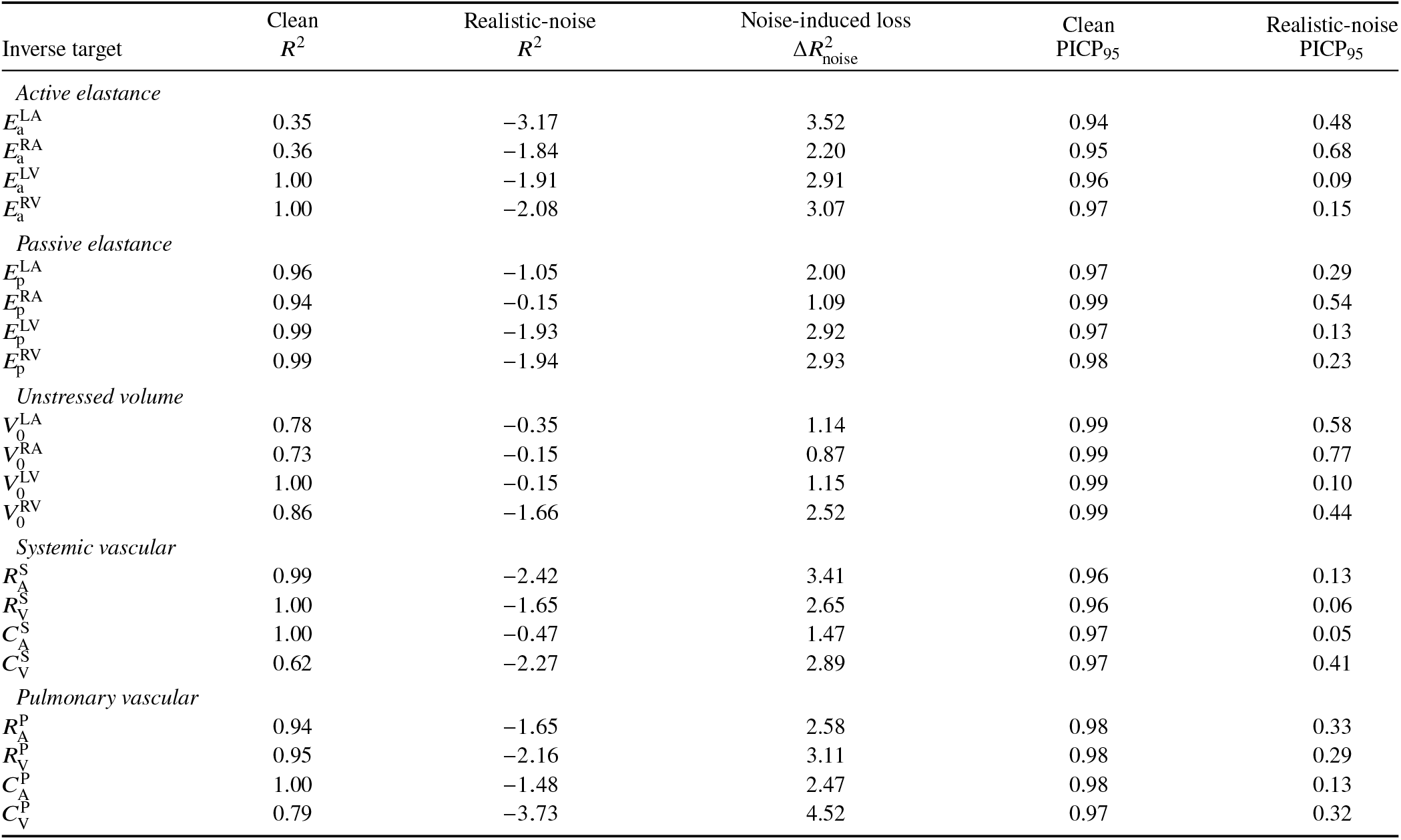
Parameter-wise robustness of the frozen full-observation Mada model to the prescribed modality-dependent measurement perturbations (*N*_test_ = 65,000). The noise-induced loss is defined 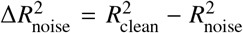. Estimates are projected to training extrema. PICP_95_ denotes the empirical coverage of the nominal 95% predictive intervals with frozen clean-validation calibation.

### D.2 Detailed Measurement-Noise Results

Across the five retained diagnostic cases, the median mean range-normalized parameter error for selected TRF increased from 0.144 under clean inversion to 0.223 under noisy inversion. Median noisy-fit NRMSE was 0.146, and median clean-reference forward NRMSE increased from 0.002 to 0.264. These five-case statistics are distinct from the 65,000-case population-level neural noise results reported above. Clean and noisy TRF solutions were selected separately according to their respective fitting residuals, without using generating-parameter error. All ten noisy TRF runs converged. All five noisy Mada replays and their clean-reference replays converged periodically.

Fitting noisy summaries can move the estimated parameters away from the clean generating state. The initialization with the lower noisy fitting residual need not have the lower parameter or clean-reference error.

### D.3 Probabilistic ResMLP Architecture Comparison

Because model size and training budget were not matched, this experiment tests whether the main trends persist in a simpler architecture; it is not a strict component ablation.

The simple probabilistic ResMLP used the same mask-aware normalized input as Mada,

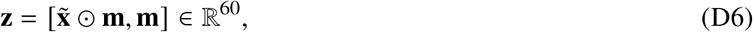

where 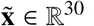 is the normalized measurement vector and **m** *∈* {0, 1}^30^ is the corresponding availability mask.

The model used the same mask-aware backbone as Mada: hidden dimension 256, two fully connected residual blocks, GELU activation, LayerNorm, and dropout 0.1. All Mada-specific prior operations were removed. In particular, the simple model did not use the frozen Stage 1 reference, recoverability-weighted reference regularization, or the *σ*_prior, *j*_ -centered uncertainty formulation.

Let **h** *∈* ℝ^256^ denote the final hidden representation. The simple model used direct mean and log-variance heads,

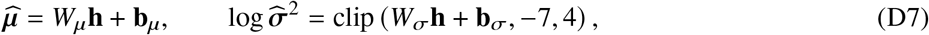

where *W*_*µ*_ and *W*_*σ*_ are trainable output-weight matrices, **b**_*µ*_ and **b**_*σ*_ are the corresponding bias vectors, and ***µ*** and 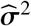 denote the predicted 20-dimensional mean and variance vectors, respectively. The clipping operation is applied element-wise. The model used the same heteroscedastic Gaussian negative log-likelihood (NLL).

All data-dependent aspects were held fixed: train/validation/test split, normalization, 30 inputs, 20 targets, mask distribution, and evaluation masks. The simple model was trained with AdamW, initial learning rate 3 × 10−4, weight decay 10−4, batch size 256, gradient clipping at 5, maximum 100 epochs, patience 30, and learning-rate reduction by 0.5 after 10 validation epochs without improvement to a minimum of 10−6. Only seed 20 was used. The evaluated mask-aware Mada Stage 2 predictor contained 428,348 trainable parameters; the pretrained Stage 1 reference remained frozen during Stage 2 optimization. The simple probabilistic ResMLP contained 291,624 trainable parameters, 31.9% fewer than the Mada Stage 2 predictor.

Both models were evaluated on the same 65,000 held-out cases under complete observations, 10%, 30%, 50%, and 70% random missingness, and the six fixed observation sets. Random-missingness calibration followed the same missingness-conditioned validation procedure used for Mada. For the fixed observation scenarios, the calibration factor in Eq. (B1), clipped to [0.4, 2.5], was estimated separately for each model, scenario, and inverse target using validation data only. The primary metrics were macro *R*^2^, macro NRMSE, PICP_95_, Std./RMSE, calibrated Gaussian NLL, and raw error–uncertainty correlation. Parameter-wise Mada advantage was

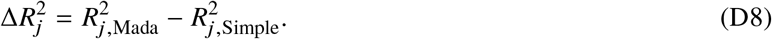

An absolute macro-*R*^2^ difference below 0.01 was used as a descriptive practical-similarity margin rather than a formal equivalence test.

Under complete observations, macro *R*^2^ was approximately 0.842 for mask-aware Mada and 0.794 for the simple model. At 70% missingness, the corresponding values were 0.337 and 0.312. Across the six fixed observation sets, both models showed the same broad trend from sparse to rich observations, although the ordering of the intermediate observation sets was not identical. Overall, the two formulations showed similar degradation with increasing measurement loss, while the magnitude and direction of their point-prediction differences varied by observation condition. Uncertainty metrics did not show a consistent advantage for either model.

### D.4 Recoverability-Weighted Reference Regularization

#### D.4.1 Motivation

The Stage 1 predictor provides both an empirical reference estimate and a parameter-wise characterization of inverse recoverability. Parameter-specific recoverability scores determine the strength of regularization toward the fixed Stage 1 reference prediction.

Let

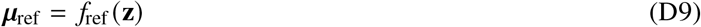

denote the prediction of the frozen Stage 1 reference model, and let ***µ*** denote the final predictive mean produced by the fully trainable Stage 2 probabilistic model.

This formulation gives the recoverability prior a direct interpretation. Parameters that are predicted reliably by the Stage 1 model are encouraged to remain closer to the reference estimate, whereas parameters with weaker empirical recoverability retain greater freedom to be relearned by the Stage 2 predictor.

#### D.4.2 Recoverability-weighted objective

The predictive negative log-likelihood is

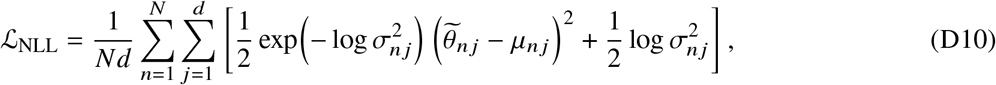

where *N* is the number of samples, *d* = 20 is the number of inverse targets, and all quantities are evaluated in normalized parameter space.

The validation-derived recoverability scores are normalized to unit mean,

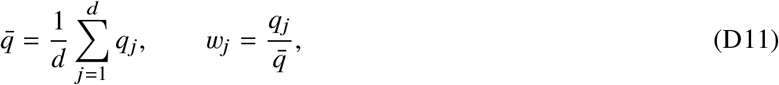

and the reference regularization is defined as

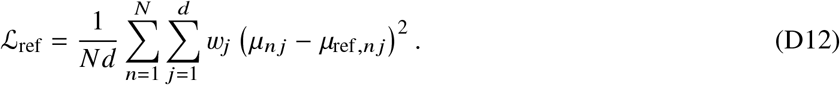

The penalty directly constrains the Stage 2 predictive mean relative to the fixed Stage 1 reference. The complete Stage 2 objective is

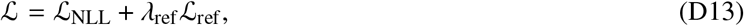

where *λ*_ref_ ≥ 0 controls the strength of the reference regularization.

#### D.4.3 Experimental protocol

The regularization sweep used the full-observation dataset and the same training, validation, and test partition as the primary full-observation experiments. All input and target normalization statistics were estimated from the training subset.

For every candidate value of *λ*_ref_, the same pretrained Stage 1 reference predictor, validation-derived recoverability scores, and residual-scale priors were used. The Stage 1 predictor remained fixed and was excluded from Stage 2 optimization.

To isolate the effect of reference regularization, all Stage 2 candidates were initialized from the same model state and used the same random seed, training samples, mini-batch ordering, stochastic sequence, optimizer settings, and checkpoint-selection rule. Optimization used AdamW with an initial learning rate of 10−3, weight decay of 10−4, batch size 256, and a maximum of 100 epochs. The learning rate was reduced on validation plateaus, and for every candidate the checkpoint with the lowest *unregularized* validation NLL was retained.

The following reference-regularization strengths were evaluated:

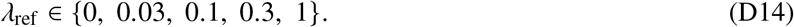

The *λ*_ref_ = 0 model therefore provides a strictly matched unregularized Stage 2 control.

#### D.4.4 Full-data results

The matched validation sweep selected *λ*_ref_ = 0.1, which achieved the lowest validation NLL (−1.453). On the held-out test set, this model achieved macro *R*^2^ = 0.862, NRMSE = 0.276, and PCC = 0.921.

The matched unregularized model (*λ*_ref_ = 0) achieved macro *R*^2^ = 0.859, NRMSE = 0.280, and PCC = 0.919. Thus, weak reference regularization increased macro *R*^2^ by 0.003 and reduced NRMSE by approximately 1.6% relative to the matched unregularized model.

A smaller regularization weight (*λ*_ref_ = 0.03) produced a smaller improvement, whereas stronger regularization did not further improve generalization. At *λ*_ref_ = 0.3, macro *R*^2^ decreased to 0.860, and at *λ*_ref_ = 1 it decreased further to 0.856. These results indicate that the Stage 1 reference is most useful as a weak prior rather than as a strong constraint on the Stage 2 predictor.

Table 17 summarizes the regularization sweep.

**Table 17.** Matched reference-regularization sweep for Mada over *λ*_ref_. The macro performance metrics and the best validation negative log-likelihood (NLL) are reported for each configuration. Boldface identifies the selected *λ*_ref_ = 0.1, which has the lowest unregularized validation NLL (Gaussian constant omitted).

| Model | $\lambda_{\text{ref}}$ | Macro $R^2$ | Macro NRMSE | Macro PCC | Best validation NLL |
| --- | --- | --- | --- | --- | --- |
| Mada | 0 | 0.859 | 0.280 | 0.919 | -1.434 |
| Mada | 0.03 | 0.860 | 0.280 | 0.919 | -1.436 |
| **Mada** | **0.1** | **0.862** | **0.276** | **0.921** | **-1.453** |
| Mada | 0.3 | 0.860 | 0.280 | 0.919 | -1.440 |
| Mada | 1 | 0.856 | 0.284 | 0.917 | -1.427 |

**Table 18.** Normalized forward-Jacobian conditioning at *h* = 0.01. Effective rank uses the relative singular-value cutoff 10^−3^.

| Case | Condition number | Effective rank |
| --- | --- | --- |
| 15 | $3.28 \times 10^7$ | 16 |
| 17 | $1.33 \times 10^9$ | 15 |
| 08 | $2.38 \times 10^7$ | 17 |
| 06 | $1.91 \times 10^7$ | 17 |
| 07 | $2.54 \times 10^7$ | 16 |

**Table 19.**
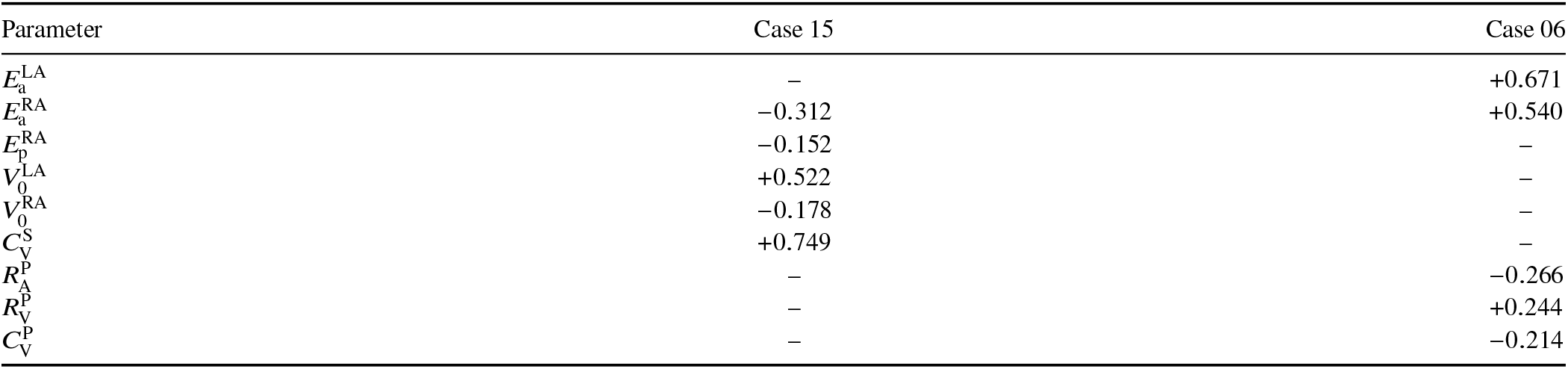
Five largest-magnitude weakest-direction loadings for cases 15 and 06 at *h* = 0.01. A dash denotes a parameter outside that case’s top five. The largest-magnitude loading is oriented positive.

| Parameter | Case 15 | Case 06 |
| --- | --- | --- |
| $E_a^{LA}$ | – | +0.671 |
| $E_a^{RA}$ | –0.312 | +0.540 |
| $E_p^{RA}$ | –0.152 | – |
| $V_0^{LA}$ | +0.522 | – |
| $V_0^{RA}$ | –0.178 | – |
| $C_V^S$ | +0.749 | – |
| $R_A^P$ | – | –0.266 |
| $R_V^P$ | – | +0.244 |
| $C_V^P$ | – | –0.214 |

**Table 20.** Local conditioning and selected-TRF recovery. Both selected runs converged; small forward residuals coexist with substantial parameter errors.

| Metric | Case 15 | Case 06 |
| --- | --- | --- |
| $\kappa_2(\bar{\mathbf{J}})$ | $3.28 \times 10^7$ | $1.91 \times 10^7$ |
| Forward NRMSE | $1.07 \times 10^{-2}$ | $1.86 \times 10^{-3}$ |
| Mean parameter error | 0.159 | 0.144 |
| Largest atrial error | 0.460 | 0.858 |
| Optimizer converged | Yes | Yes |

**Table 21.**
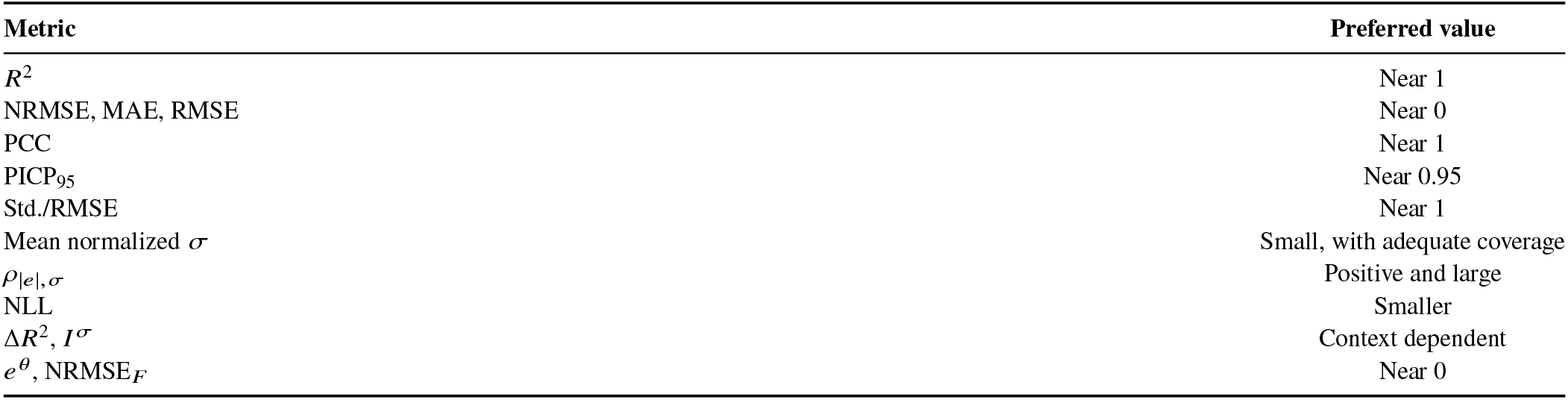
Preferred direction or reference value for the principal evaluation metrics.

**Table 22.** Direct-optimizer convergence and elapsed wall time over both starts (40 runs per method). All 120 final LPN replays converged periodically.

| Method | Optimizer Converged | Runtime (h)<br>median / max | Reported nfev<br>median / max | Forward calls<br>median / max |
| --- | --- | --- | --- | --- |
| TRF | 40/40 | 0.808 / 6.931 | 40.5 / 296 | 533 / 5297 |
| Dogbox | 35/40 | 1.969 / 92.046 | 68.5 / 2000 | 1399.5 / 41981 |
| L-BFGS-B | 31/40 | 5.149 / 35.375 | 4126.5 / 14805 | 4127.5 / 14806 |

Five Dogbox runs reached the evaluation limit; nine L-BFGS-B runs ended in line-search failure.

#### D.4.5 Reference-regularization behavior

The magnitude of the reference penalty provides an additional view of how *λ*_ref_ influences Stage 2 learning. At the selected *λ*_ref_ = 0.1 checkpoint, L_ref_ = 0.023, corresponding to a contribution

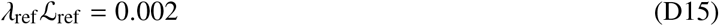

to the total objective.

Increasing *λ*_ref_ beyond the selected value more strongly constrained the final prediction toward the Stage 1 reference. For example, L_ref_ decreased to 0.020 at *λ*_ref_ = 0.3 and to 0.018 at *λ*_ref_ = 1. This stronger constraint, however, was accompanied by reduced test-set predictive accuracy.

Notably, the weakly regularized model did not simply produce uniformly smaller departures from the Stage 1 reference than the unregularized model. The improvement at *λ*_ref_ = 0.1 should therefore be interpreted as a modest regularization effect on Stage 2 learning rather than as evidence that prediction improves merely by remaining closer to the Stage 1 estimate.

Figure 20 summarizes the predictive performance, validation behavior, and strength of reference regularization across the matched sweep.

**Figure 16.**
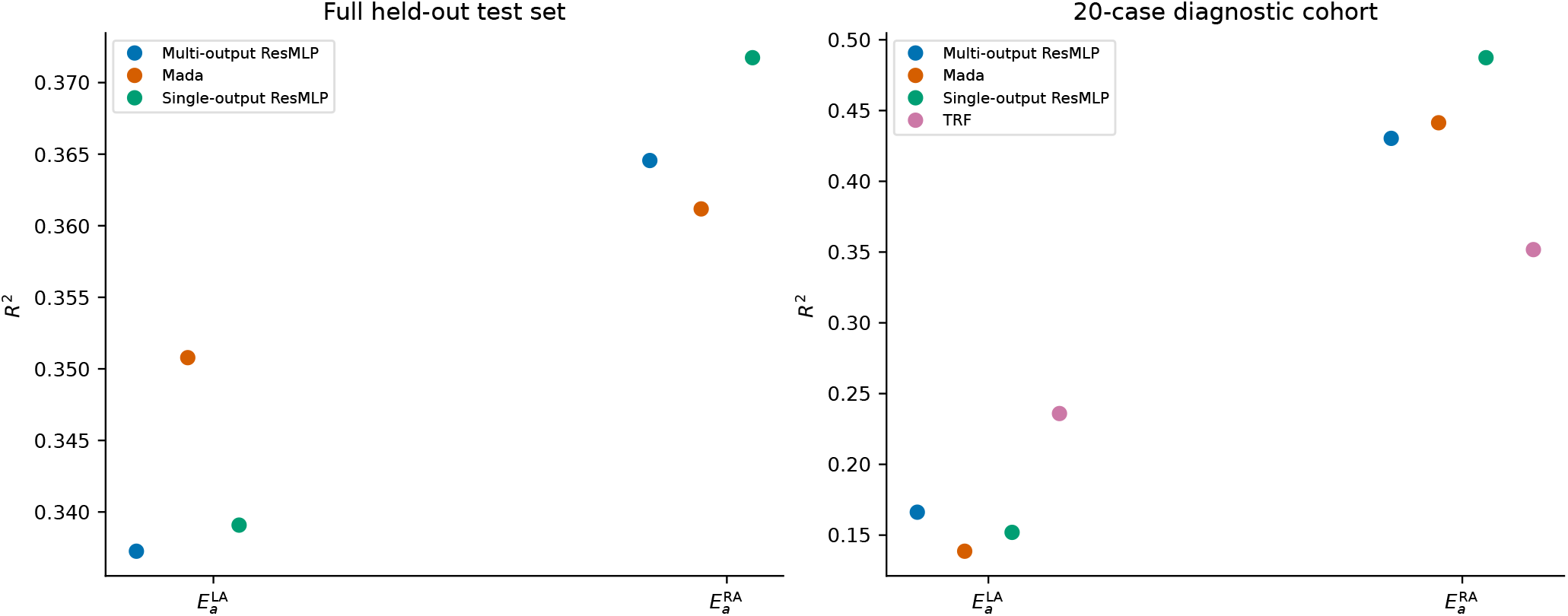
Single-output diagnostic of atrial active-elastance learnability. (a) Held-out test-set *R*^2^ for the multi-output ResMLP, Mada, and single-output ResMLPs. (b) Cross-case *R*^2^ for the same neural models and selected TRF on the 20 optimization cases. Neural results use unprojected predictions; Mada is the dedicated full-observation model reported in Table 2.

**Figure 17.**
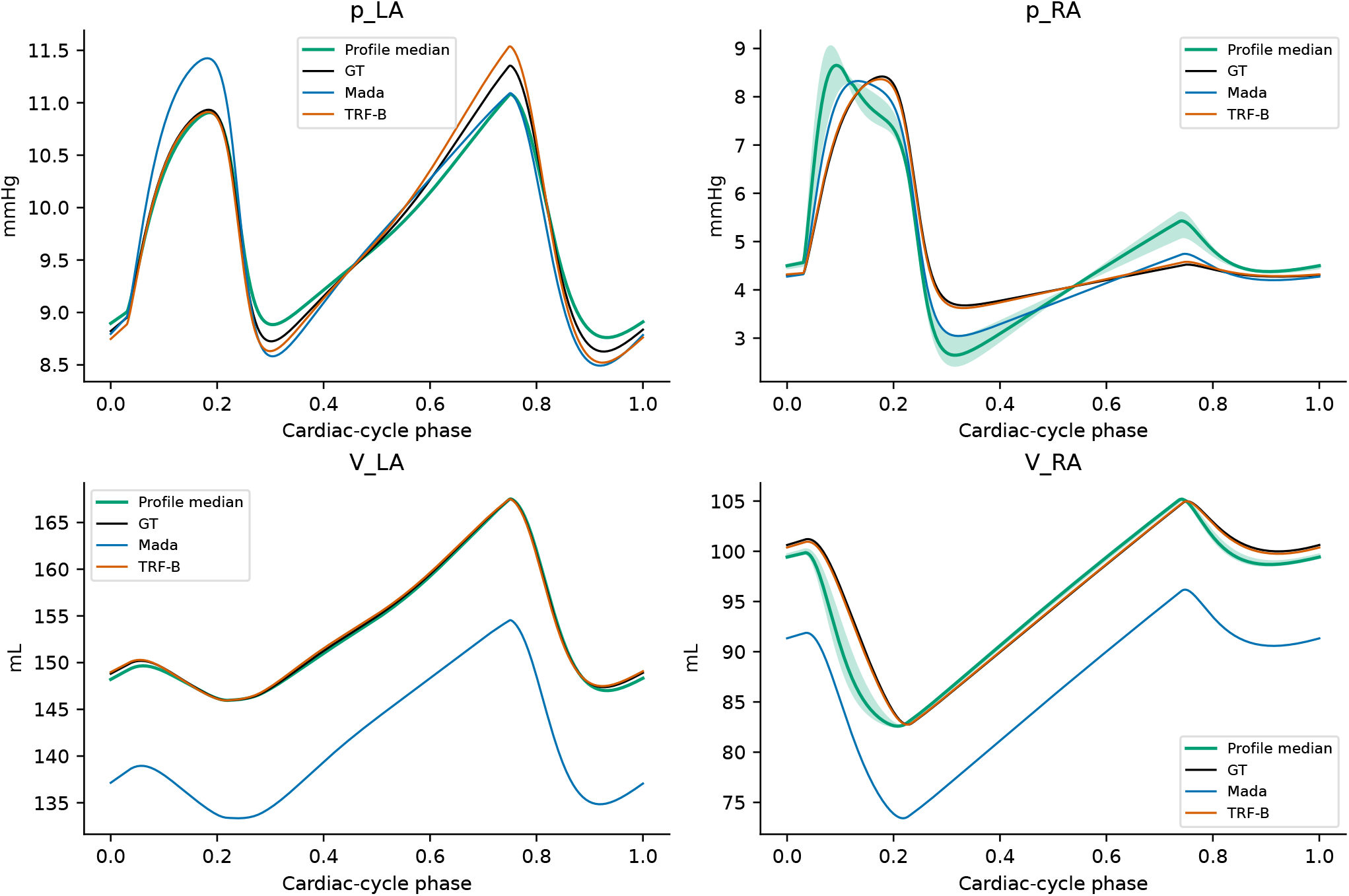
Case 07 waveforms: median and 5th–95th percentile envelope over the four qualifying joint-profile solutions, with GT, Mada, and selected TRF. This envelope is not a predictive or posterior interval.

**Figure 18.**
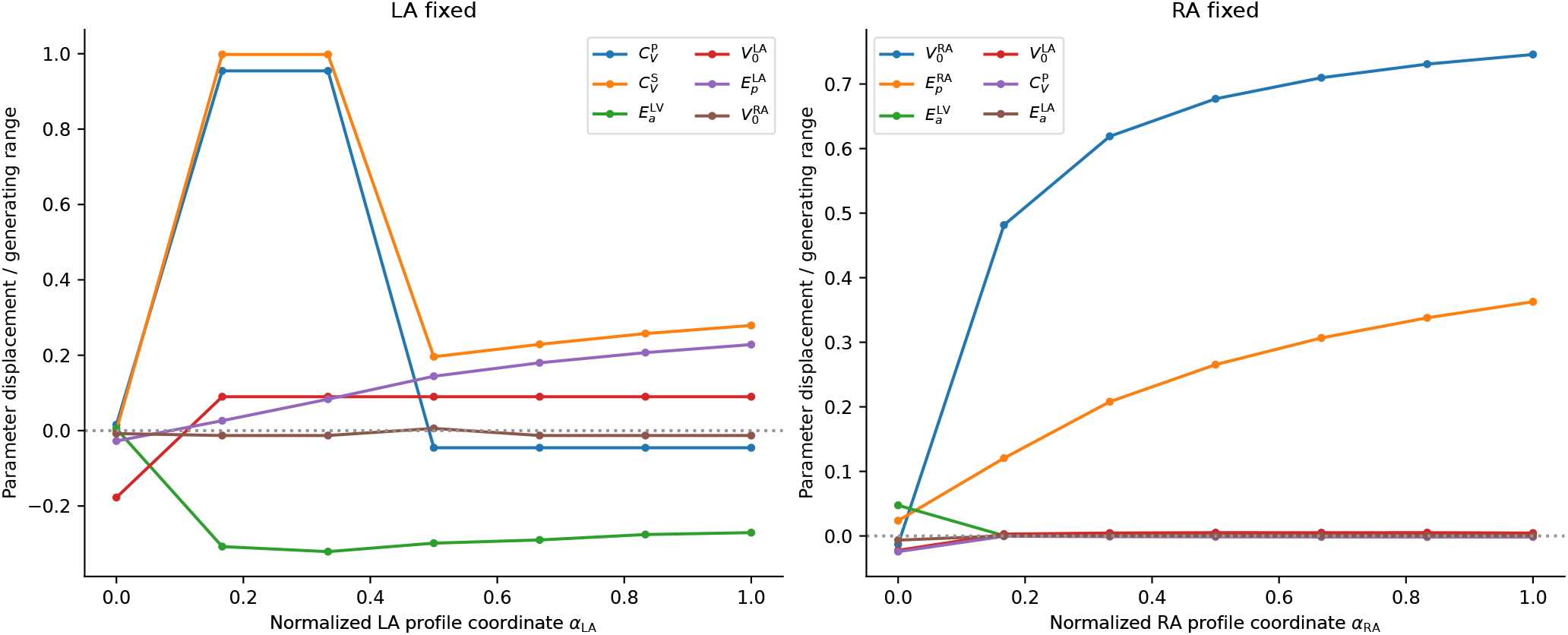
Compensatory displacements along the LA and RA profiles versus normalized profile coordinate *α*_*j*_ . The six largest peak-to-peak displacements are shown for each profile, excluding its fixed coordinate. Points are independently optimized; connecting lines do not imply a continuous optimization trajectory.

**Figure 19.**
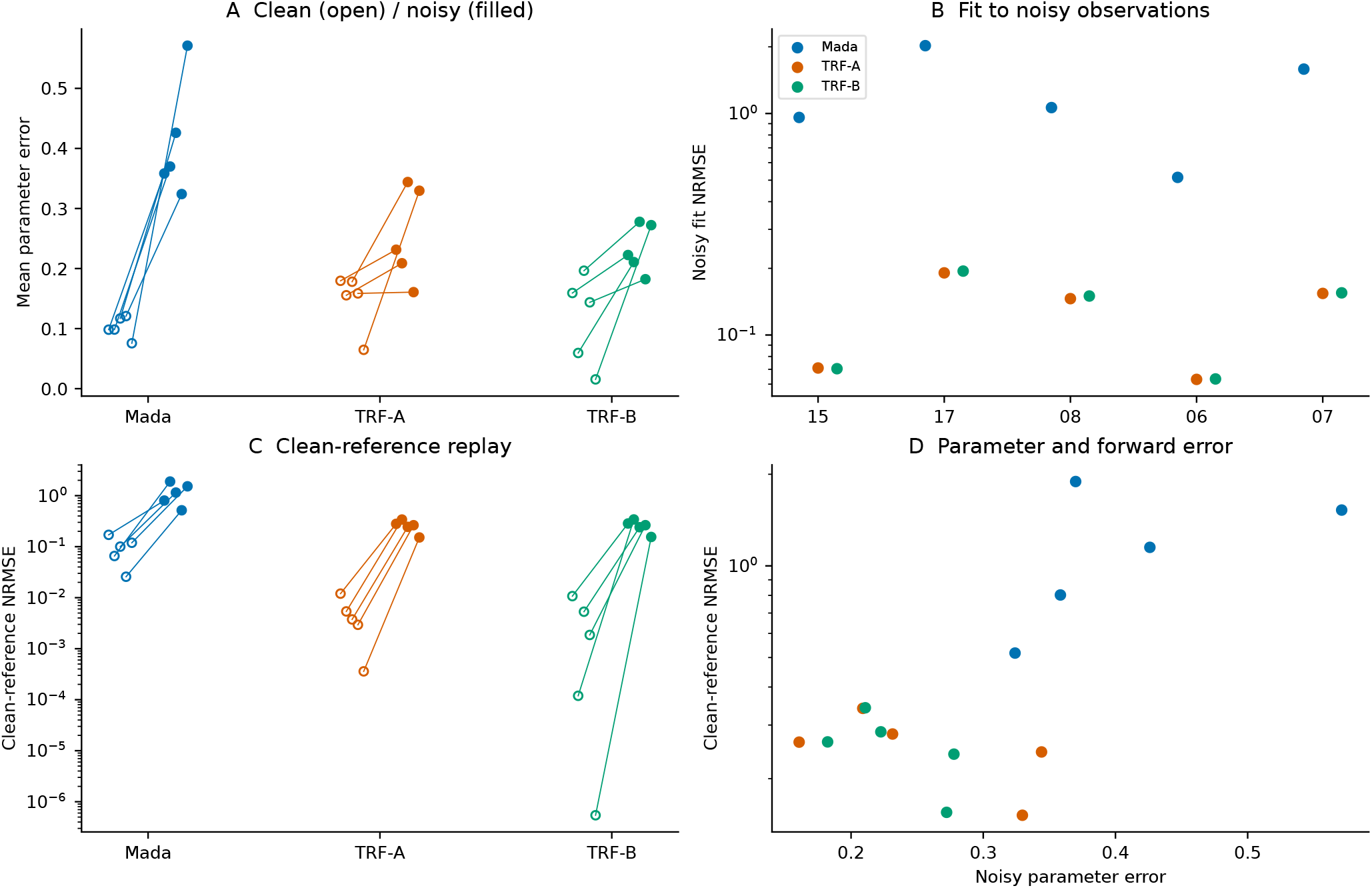
Five-case measurement-noise diagnostic with the original LPN forward solver. A: clean/noisy mean parameter errors. B: noisy-observation fitting residual. C: clean-reference replay error. D: noisy parameter error versus clean-reference error. All five Mada estimates are shown; noisy-fit and clean-reference replays converged periodically.

**Figure 20.**
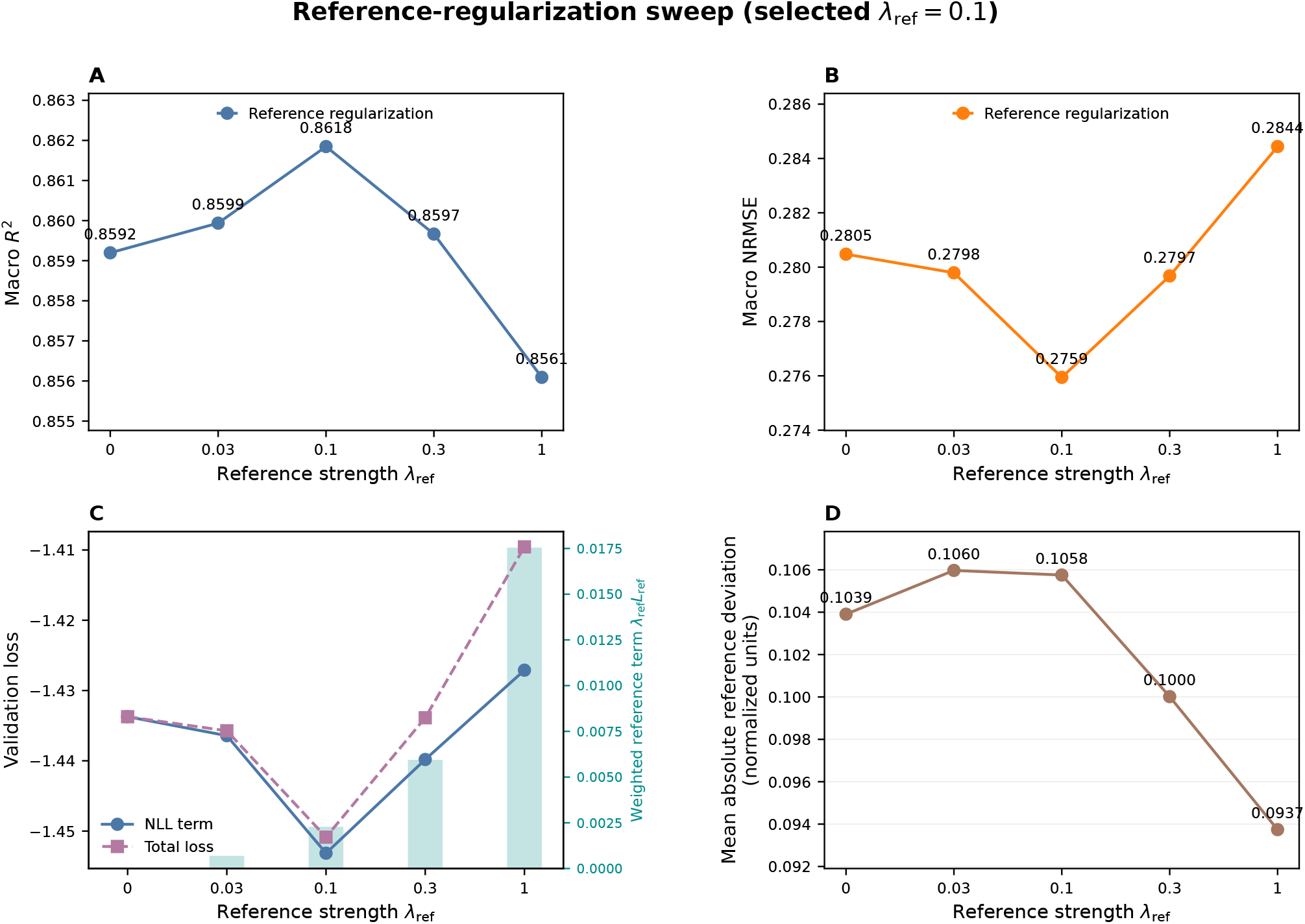
Extended full-data reference-regularization sweep. (A) Test-set macro *R*^2^ across *λ*_ref_ *∈* {0, 0.03, 0.1, 0.3, 1}. (B) Test-set macro NRMSE. (C) Unregularized validation NLL, total validation loss, and the weighted reference-regularization contribution at the selected checkpoint for each *λ*_ref_. (D) Mean absolute deviation of the Stage 2 prediction from the frozen Stage 1 reference. The validation sweep selected *λ*_ref_ = 0.1, which achieved the best test-set macro *R*^2^ among the matched candidates; stronger regularization reduced predictive accuracy.

#### D.4.6 Interpretation

The matched sweep supports a modest benefit from weak recoverability-weighted reference regularization in Stage 2 training. The weights *q*_*j*_ were held fixed; the experiment did not compare recoverability weighting with uniform reference weighting. The Stage 1 recoverability scores remain useful as empirical descriptors of parameter-wise inverse predictability.

The selected *λ*_ref_ = 0.1 model improved macro *R*^2^ only modestly relative to the strictly matched unregularized Stage 2 control. Accordingly, the reference penalty should be interpreted as a weak inductive prior rather than as the primary source of predictive performance. Stronger regularization reduced accuracy, indicating that the Stage 2 model must retain sufficient freedom to depart from the deterministic Stage 1 reference when supported by the observations.

Because the regularization comparison was performed using a single matched random seed, the small performance improvement should not be interpreted as evidence of a large or universal architectural advantage. Its principal value is that it incorporates validation-derived recoverability information through a well-defined constraint on the final prediction while preserving the flexibility of the probabilistic Stage 2 model.

### D.5 Measurement-Noise Amplitude Sweep

#### D.5.1 Noise-scaling protocol

The measurement-noise analysis evaluated the frozen full-observation Mada model across a common-amplitude sweep of the prescribed modality-dependent perturbations. The full prescribed condition corresponds to *γ*_noise_ = 1, while the lower amplitudes test whether degradation begins progressively before the full noise level is reached. The common noise multiplier is denoted by *γ*_noise_.

For every measurement subject to additive noise, the scaled perturbation was defined as

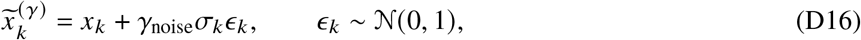

whereas measurements subject to multiplicative noise were perturbed as

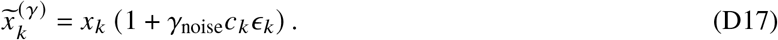

The modality-specific values of *σ*_*k*_ and *c*_*k*_, together with the rules used to recompute derived hemodynamic quantities, were identical to those described in Appendix D.1. The four evaluated noise scales were

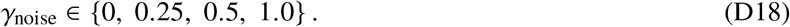

Thus, *γ*_noise_ = 0 corresponds to the clean test set, whereas *γ*_noise_ = 1 reproduces the full prescribed measurement-noise condition. The same seed-20 standard-normal realization was scaled only through *γ*_noise_, so differences across conditions reflect changes in noise amplitude rather than stochastic realization.

The full-observation Mada checkpoint and clean-validation calibration factors remained fixed throughout the sweep. Performance was evaluated using macro test *R*^2^, macro NRMSE, the mean predicted-standard-deviation-to-RMSE ratio, and the mean 95% prediction-interval coverage probability 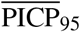.

The aggregate results for all four noise amplitudes are summarized in Table 5.

#### D.5.2 Noise-amplitude dependence of prediction and calibration

Table 5 shows a monotonic deterioration in point-prediction performance as measurement-noise amplitude increased. At noise multipliers *γ*_noise_ = 0, 0.25, 0.5, and 1.0, macro *R*^2^ was 0.862, −0.053, −0.749, and −1.610, respectively; macro NRMSE was 0.276, 0.959, 1.279, and 1.586. Mean Std./RMSE decreased from 1.046 to 0.281, 0.226, and 0.190, while mean PICP_95_ decreased from 0.974 to 0.474, 0.386, and 0.310. Thus, even one-quarter of the prescribed perturbation amplitude produced negative macro *R*^2^ and substantial interval undercoverage.

#### D.5.3 Interpretation

The amplitude sweep shows that the full prescribed noise condition is not an isolated threshold at which the model suddenly fails. Instead, both inverse-prediction accuracy and uncertainty calibration worsen systematically as measurement noise increases. However, the especially large change between *γ*_noise_ = 0 and 0.25 indicates that uncertainty calibration is highly sensitive even to relatively small departures from the clean synthetic-data distribution.

These findings reinforce the interpretation that calibration derived from clean validation data is distribution specific. It can correct the average uncertainty scale for clean or similarly distributed observations, but it does not automatically account for measurement error that was absent during model training and calibration. Robust application to measured clinical data will therefore require the relevant device- and modality-specific error distributions to be represented during training, uncertainty calibration, or both.

The present sweep remains a controlled robustness diagnostic. It uses one fixed stochastic realization and scales all modality-specific perturbations by a common factor; it does not represent correlated device bias, calibration drift, temporal misalignment, disease-dependent heteroscedasticity, or repeated independent noise realizations. These effects should be examined before the reported uncertainty estimates are interpreted as clinically calibrated prediction intervals.

### D.6 Local Forward-Sensitivity Diagnostics

#### D.6.1 Motivation

The five-case analysis evaluates local observational directions at the generating parameters using the same LPN forward solver. It characterizes sensitivity rather than establishing global structural identifiability.

#### D.6.2 Normalized Forward Jacobian

The 27 outputs and 20 parameters are standardized using the training-set means and standard deviations:

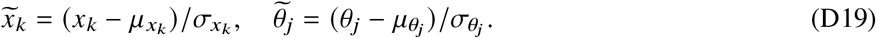

Timing is fixed. At the generating vector,

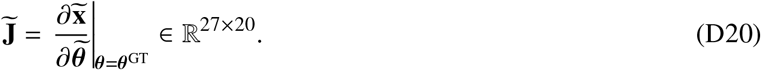

Centered differences perturb physical parameter *j* by 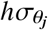, without clipping the stencil:

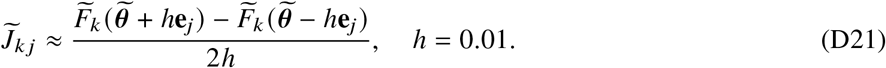

The same calculation is available at *h* = 0.005 and 0.02. The singular-value decomposition, spectral condition number, and effective rank are

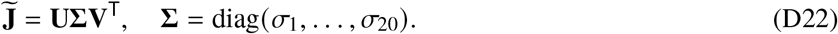

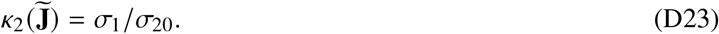

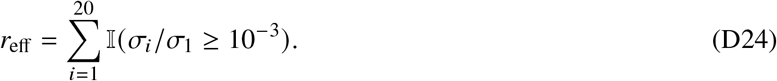

The last right singular vector describes the weakest local standardized direction. Its sign is fixed by making the largest-magnitude loading positive.

#### D.6.3 Local Sensitivity Results

At *h* = 0.01, condition numbers for Cases 15, 17, 08, 06, and 07 were 3.28 × 10^7^, 1.33 × 10^9^, 2.38 × 10^7^, 1.91 × 10^7^, and 2.54 × 10^7^, respectively. Effective ranks were 16, 15, 17, 17, and 16 and remained unchanged across all three steps. Figure 21 shows the relative spectra.

**Figure 21.**
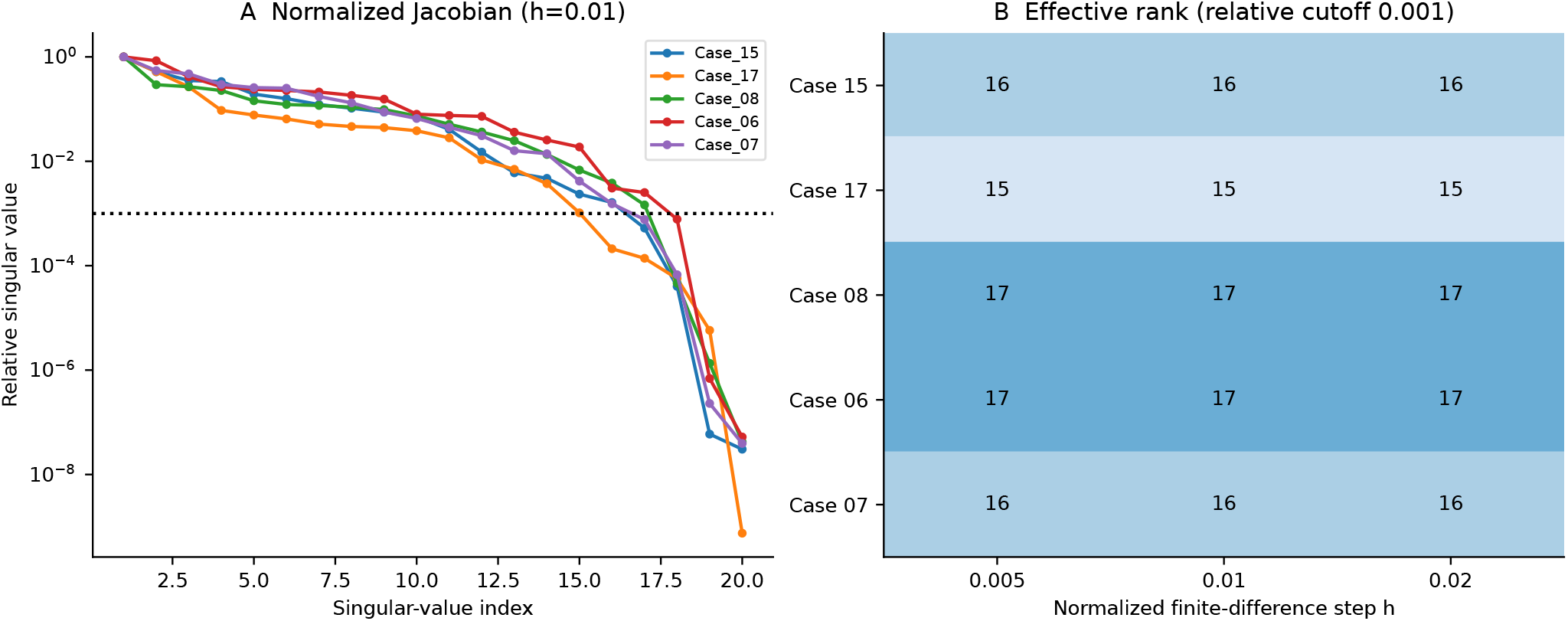
Original LPN normalized-Jacobian spectra at generating parameters. A: relative singular values at *h* = 0.01 and rank cutoff 10^−3^. B: effective ranks at all three finite-difference steps.

The largest five loadings for Cases 15 and 06 give the truncated directions

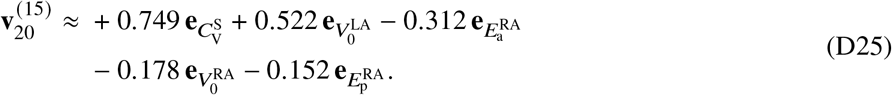

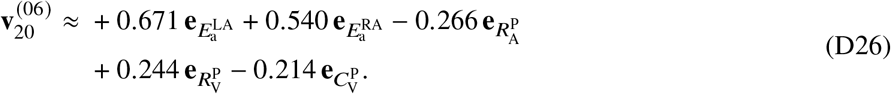

Systemic venous compliance has the largest Case 15 loading, whereas both atrial active elastances dominate Case 06. These directions are local and depend on numerical step. At *h* = 0.02, absolute cosine similarity to the *h* = 0.01 weakest vector falls to 0.227 for Case 15 and 0.050 for Case 08. Thus, unchanged effective rank does not imply unchanged weakest-vector composition; Table 23 reports the full step comparison.

**Table 23.** Step-size sensitivity of local Jacobian conditioning and weakest directions. Cosine is the absolute inner product with the unit weakest vector at *h* = 0.01.

| Case | $h$ | Condition number | Rank | Absolute cosine |
| --- | --- | --- | --- | --- |
| 15 | 0.005 | $1.38 \times 10^8$ | 16 | 1.000 |
| 15 | 0.01 | $3.28 \times 10^7$ | 16 | 1.000 |
| 15 | 0.02 | $1.14 \times 10^7$ | 16 | 0.227 |
| 17 | 0.005 | $5.30 \times 10^9$ | 15 | 1.000 |
| 17 | 0.01 | $1.33 \times 10^9$ | 15 | 1.000 |
| 17 | 0.02 | $3.50 \times 10^8$ | 15 | 1.000 |
| 08 | 0.005 | $3.15 \times 10^7$ | 17 | 1.000 |
| 08 | 0.01 | $2.38 \times 10^7$ | 17 | 1.000 |
| 08 | 0.02 | $7.15 \times 10^5$ | 17 | 0.050 |
| 06 | 0.005 | $9.18 \times 10^7$ | 17 | 1.000 |
| 06 | 0.01 | $1.91 \times 10^7$ | 17 | 1.000 |
| 06 | 0.02 | $4.82 \times 10^6$ | 17 | 1.000 |
| 07 | 0.005 | $1.09 \times 10^8$ | 16 | 0.999 |
| 07 | 0.01 | $2.54 \times 10^7$ | 16 | 1.000 |
| 07 | 0.02 | $7.43 \times 10^6$ | 16 | 0.977 |

#### D.6.4 Comparison With Direct Optimization

Selected TRF in Case 15 achieved forward NRMSE 0.011, mean parameter error 0.159, and largest atrial error 0.460. Case 06 achieved 0.002, 0.144, and 0.858, respectively. Both selected optimizers converged. Mada in Case 06 had forward NRMSE 0.026 and mean parameter error 0.121: lower forward residual therefore accompanied a larger parameter error in this example. Conversely, Case 07’s selected TRF mean parameter error was only 0.015. These heterogeneous cases do not define a condition-number threshold for recovery.

Local LPN sensitivities describe weak observational directions, whereas the validation-derived recoverability scores in Appendix D.4 characterize empirical neural prediction difficulty.

## E Evaluation Metrics and Interpretation

This section summarizes the evaluation metrics used throughout the study. The purpose is to distinguish three related but different questions: (i) whether the inferred LPN parameters are accurate, (ii) whether the reported predictive uncertainty is calibrated, and (iii) whether the inferred parameters reproduce the target hemodynamics. Unless stated otherwise, metrics are first computed separately for each inverse target and are then macro-averaged over the *d* = 20 LPN parameters.

Let *N* denote the number of test cases and let *d* = 20 denote the number of inverse targets. The index *n* = 1, …, *N* denotes a test case, whereas *j* = 1, …, *d* denotes an inverse target. Let *θ*_*n j*_ denote the generating value of parameter *j* in case *n*, and let 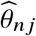 denote its predicted mean. Let 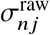 and 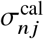 denote the corresponding raw and validation-calibrated predictive standard deviations, respectively, in physical parameter units. Unless explicitly stated otherwise, interval coverage, Std./RMSE, and mean predictive-standard-deviation results use 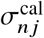. Tables explicitly labeled as reporting raw uncertainty instead use 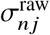. The indicator function **1**(·) equals one when its argument is true and zero otherwise.

### E.1 Point-Prediction Metrics

The root-mean-square error (RMSE) and mean absolute error (MAE) for parameter *j* are

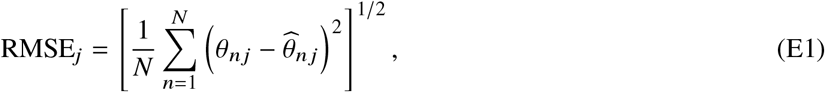

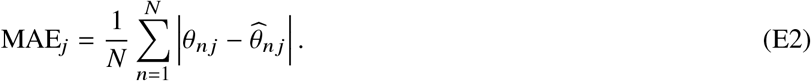

Both metrics have the physical units of the corresponding parameter, and lower values indicate more accurate predictions. RMSE weights large errors more strongly than MAE.

To permit comparisons among parameters with different units and scales, let *s*_*j*,train_ denote the standard deviation of inverse target *j* in the training subset *D*_train_,

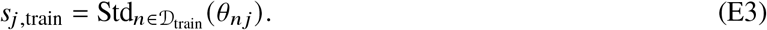

The normalized root-mean-square error is then

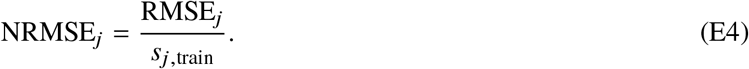

Thus, NRMSE_*j*_ = 0 indicates exact prediction, while smaller values indicate better recovery. The normalization is distribution dependent: the same absolute error can produce a different NRMSE if the spread of the target population changes.

The coefficient of determination is

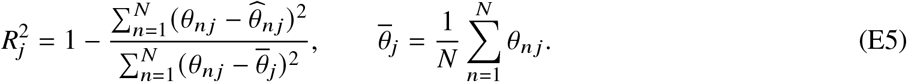

An 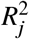 value near one indicates accurate case-to-case recovery. A value of zero means that the predictions are no better, in squared-error terms, than using the test-set mean for every case; 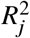 can be negative when the model performs worse than that reference. In this work, 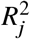 is interpreted as an empirical recoverability measure for the specified population, observation set, and inference method, not as a structural-identifiability index.

The Pearson correlation coefficient (PCC) measures linear association between predicted and recorded generating values,

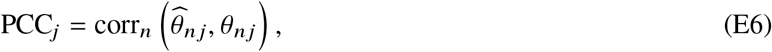

where corr_*n*_ (·, ·) denotes the Pearson correlation computed across the *N* test cases. A PCC near one indicates strong positive linear association, but PCC alone does not detect a constant offset or positive multiplicative scaling bias. It is therefore reported together with *R*^2^ and NRMSE.

Macro values are unweighted averages across the *d* = 20 inverse targets:

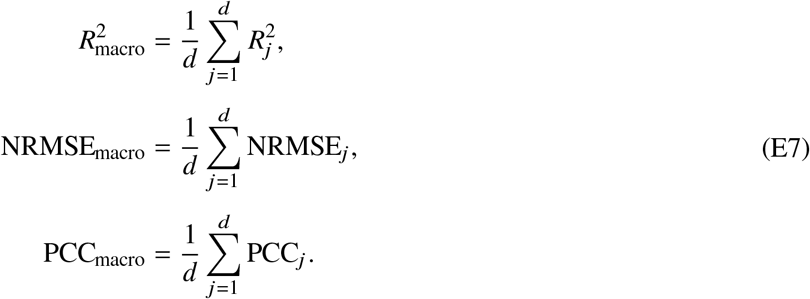

Macro averaging gives each physiological parameter equal weight, regardless of its physical units or numerical range, but can conceal difficult individual targets. Parameter-wise results are therefore also reported.

### E.2 Predictive-Uncertainty Metrics

For test case *n* and inverse target *j*, let 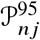 denote Mada’s nominal 95% Gaussian prediction interval. It is defined as

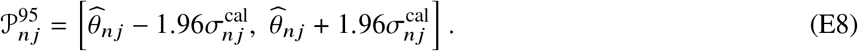

This is a prediction interval for the unknown parameter of an individual test case. It is not a confidence interval for a population mean and does not, by itself, establish uniqueness or structural identifiability of the inverse problem.

The 95% prediction-interval coverage probability (PICP_95_) is the observed fraction of recorded generating values contained within these intervals,

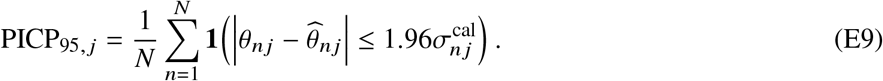

The nominal target is 0.95. Values appreciably below 0.95 indicate undercoverage: the intervals are too narrow, the predictive mean is biased, or the assumed error distribution is inadequate. Values above 0.95 indicate conservative coverage, which may result from unnecessarily wide intervals. Therefore, a larger PICP_95_ is not automatically better; proximity to the nominal level must be considered together with interval scale. For a finite test set, ordinary binomial sampling variation around 0.95 is also expected.

The average predicted uncertainty scale is compared with the realized RMSE using

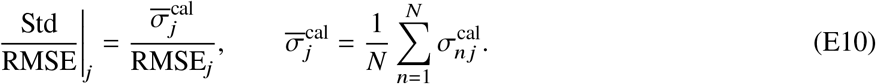

A ratio near one indicates that the average predicted standard deviation is similar to the empirical RMSE. A value below one indicates that predictive uncertainty is too small on average, whereas a value above one indicates a conservative average scale. This ratio assesses only global uncertainty magnitude and can therefore be near one even when individual prediction intervals are poorly calibrated.

Because the multiplicative calibration factor is defined from validation RMSE and mean validation predictive standard deviation, Std./RMSE primarily assesses whether this calibrated scale relationship transfers from validation to test data. It should not be interpreted as an independent validation of uncertainty quality. PICP_95_ and sample-level error–uncertainty correlation provide complementary assessments of interval coverage and case-specific uncertainty ranking.

The mean normalized predictive standard deviation reported in several figures is

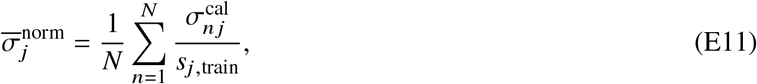

where *s*_*j*,train_ is the training-set standard deviation of inverse target *j* defined above. It measures the typical predicted uncertainty relative to the population scale. Smaller values indicate sharper predictions only when coverage remains adequate.

To assess whether Mada assigns larger uncertainty to cases with larger prediction errors, define the signed parameter-prediction error as

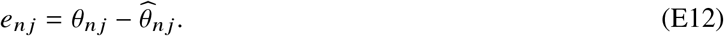

The sample-level error–uncertainty correlation for target *j* is then

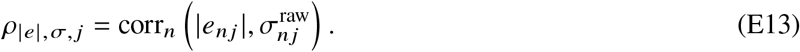

For the dedicated full-observation model and fixed or structured masks, calibration multiplies every predicted standard deviation for a given target and condition by the same positive constant, so the within-target correlation is unchanged:

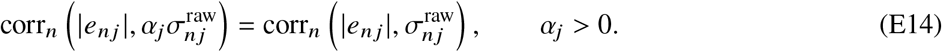

This invariance does not generally hold under random missingness, where the calibration factor depends on each sample’s actual missing fraction. Error–uncertainty correlations are therefore reported using raw standard deviations throughout.

A positive value indicates useful case-specific ranking of predictive uncertainty. However, this correlation does not guarantee correct interval coverage or uncertainty scale and is therefore interpreted jointly with PICP_95_ and Std./RMSE.

For probabilistic model comparison, the Gaussian negative log-likelihood (NLL) is evaluated in the normalized target space used for neural-network training. Let 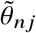 denote the normalized generating value, *µ*_*n j*_ the predicted normalized mean, and *σ*_*n j*_ the predicted normalized standard deviation for case *n* and target *j* . The NLL is

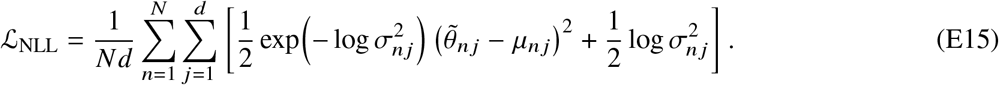

The parameter-independent Gaussian constant is omitted, as in the training objective. Lower NLL is better. It jointly penalizes inaccurate means and inappropriate uncertainty, including both overconfident errors and excessively broad distributions. NLL values are comparable only when the same target normalization and additive-constant convention are used.

### E.3 Measurement-Loss and Forward-Reconstruction Metrics

For removal of measurement group *g*, the recoverability loss and uncertainty inflation are

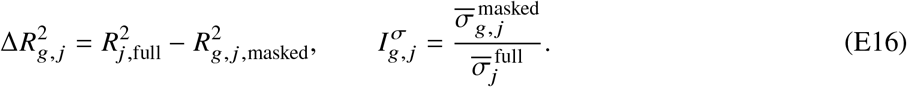

A positive 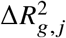 indicates that removing group *g* reduces recovery of parameter *j* . An uncertainty-inflation value greater than one means that the model reports greater uncertainty after that information is removed. These are empirical measurement–parameter dependency measures and should not be read as formal sensitivity or identifiability indices.

In the direct-optimization diagnostics, let 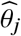 denote the recovered value of inverse target *j*, 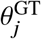 its recorded generating value, and *θ*_*j,L*_ and *θ*_*j,U*_ its original lower and upper generating bounds. Parameter recovery is summarized by the range-normalized absolute error

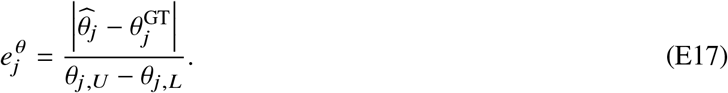

Hemodynamic reconstruction is evaluated separately. Let *K* = 27 denote the number of modeled hemodynamic outputs, *k* = 1, …, *K* index those outputs, 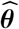 denote the recovered parameter vector, **t** the known case-specific timing vector, 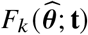 the corresponding forward-LPN prediction for output *k, x*_*k*_ the target hemodynamic quantity, and *s*_*k*_ its training-set standardization scale. The normalized forward error is

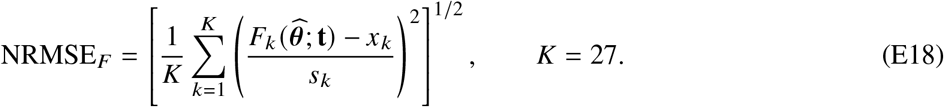

Low NRMSE_*F*_ indicates consistency with the target hemodynamic summaries, whereas low 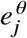 indicates accurate recovery of the recorded generating parameter. A solution may have very low forward error but substantial parameter error when different parameter combinations produce similar observed hemodynamics.

In the measurement-noise diagnostic, this forward metric is evaluated in two distinct ways. The noisy-fit error uses the perturbed observations and perturbed timing vector,

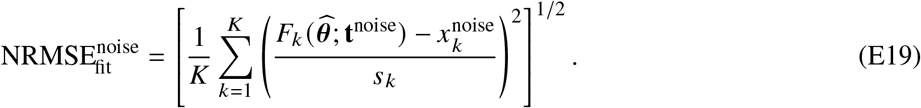

The clean-reference error evaluates the same recovered parameter vector against the original observations and timing,

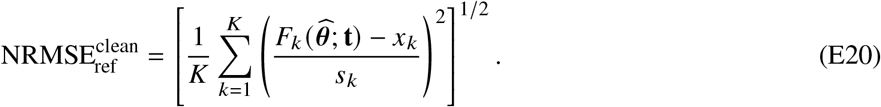

A parameter vector can therefore fit the noisy observations while showing substantially poorer agreement with the clean generating observations.

### E.4 Computational and Numerical Diagnostic Details

The following tables summarize optimizer runtime and convergence and the finite-difference sensitivity of the local forward Jacobian.

